# Reducing the Bacterial Lag Phase Through Methylated Compounds: Insights from Algal-Bacterial Interactions

**DOI:** 10.1101/2023.06.06.543872

**Authors:** Martin Sperfeld, Delia A. Narváez-Barragán, Sergey Malitsky, Veronica Frydman, Lilach Yuda, Jorge Rocha, Einat Segev

## Abstract

The bacterial lag phase is a key period for resuming growth. Despite its significance, the lag phase remains underexplored, particularly in environmental bacteria. Here, we explore the lag phase of the model marine bacterium *Phaeobacter inhibens* when it transitions from starvation to growth with a microalgal partner. Utilizing transcriptomics and ^13^C-labeled metabolomics, our study reveals that methylated compounds, which are abundantly produced by microalgae, shorten the bacterial lag phase. Our findings underscore the significance of methyl groups as a limiting factor during the lag phase and demonstrate that methyl groups can be harvested from algal compounds and assimilated through the methionine cycle. Furthermore, we show that methylated compounds, characteristic of photosynthetic organisms, induce variable reductions in lag times among bacteria associated with algae and plants. These findings highlight the adjustability of the bacterial lag phase and emphasize the importance of studying bacteria in an environmental context.

**One-Sentence Summary:** Bacteria use algal compounds as a metabolic shortcut to transition from starvation to growth.

## Introduction

Marine heterotrophic bacteria rely on organic carbon synthesized by photosynthetic microorganisms, and consume approximately half of the carbon fixed by these primary producers^1–3^. The primary producers, namely microalgae, produce organic carbon by fixing CO_2_ in a process that is subjected to temporal dynamics^4,5^. For example, photosynthetic productivity is poor at night and in winter, when light intensities decline, or nutrients are depleted. Consequently, heterotrophic bacteria must endure prolonged phases of starvation^6–8^. However, with the onset of microalgal productivity, bacteria must rapidly activate their metabolism. A rapid metabolic response is key for bacteria to outgrow co-occurring competitors^9–11^.

The transition from starvation^12,13^ and preparation for cell division occur during the lag phase^14^. Recent studies highlight that bacteria undergo a global metabolic reorganization during the lag phase^15^ and synthesize essential building blocks (such as nucleotides and amino acids), which are then incorporated into larger macromolecules including DNA, RNA and proteins^16–18^. Some building blocks can be acquired from external sources and alleviate metabolic needs during the bacterial lag phase^18^. While the crucial preparatory lag period has been well-studied in human pathogens and in food-spoiling bacteria^15^, much is unknown about the bacterial lag phase and its controls in the marine environment.

Several groups of marine heterotrophic bacteria, such as the common Roseobacter group^19^, are often found in association with microalgae^1^. Much knowledge exists regarding the metabolic pathways through which bacteria utilize algal metabolites for growth^20,21^. However, there remains a significant knowledge gap regarding the early physiological response of bacteria upon encountering algae after a period of starvation. As bacteria often endure periods of starvation in nature^22^, strategies to optimize the initial interaction with algal hosts could prove beneficial for the ecological success of marine bacteria. In this study, we sought to understand the post-starvation response of a model Roseobacter species, *Phaeobacter inhibens*, to microalgal metabolites and to determine the underlying mechanisms.

We analyzed the metabolism of *P. inhibens* while it grows with microalgae or is supplemented with selected microalgal metabolites. We discovered that methyl groups represent a metabolic “bottleneck” to bacterial growth even when ample carbon and nutrients are made available. Nano-to micromolar concentrations of methylated compounds, which are abundantly produced by microalgae^23–25^, significantly shorten the lag phase of bacteria. This observation marks methylated compounds as a class of metabolites that reduces lag times. Specifically, compounds that can donate one-carbon groups to the bacterial metabolism induce reduction of the lag phase with no influence on the bacterial growth rate. To elucidate the mechanisms that underly lag phase shortening by methylated compounds, we explored the bacterial transcriptome during the lag phase, estimated cellular requirements for methyl groups, and analyzed the incorporation of ^13^C-labeled methyl groups. Our results suggest that methyl group synthesis is a constraint during the bacterial lag phase, a limitation that can be alleviated by abundant algal metabolites. Our findings reveal the overlooked tunability of the bacterial lag phase and underscore the importance of studying bacterial physiology within an ecological context.

## Results

### Bacterial methyl group metabolism genes are highly expressed across growth phases during co-cultivation with algae

We first set out to investigate the metabolic response that is activated in *Phaeobacter inhibens* bacteria during co-cultivation with *Emiliania huxleyi* microalgae (also known as *Gephyrocapsa huxleyi*^26^). This algal-bacterial pair naturally co-occurs in the environment and was previously studied by us^27,28^ and by others^29–31^. We analyzed the transcriptome of bacteria in co-cultures with algae—a condition under which bacteria rely entirely on algal-secreted metabolites for growth^27^. As a reference condition, we analyzed bacterial pure cultures that received glucose as a sole carbon source (figs. S1-S3, tables S1-S2, and data S1). Notably, comparing pure bacterial cultures under nutrient replete conditions with glucose as a sole carbon source, and bacteria in co-cultures that receive a continuous, complex, yet limited supply of algal metabolites, contains inherent limitations. Yet, such a comparison can highlight potentially important differences and serve as a hypothesis generator for further exploration.

Based on KEGG metabolic pathways definitions^32^ we identified catabolic and anabolic processes that are highly expressed in bacteria during co-cultivation with algae. We found that genes involved in utilizing methylated compounds were among the 20 highest transcribed bacterial metabolic genes (Fig. 1A and data S1). Examples include a gene encoding a cobalamin-dependent methyltransferase (gene 3 in Fig. 1A), a gene encoding a formate—THF ligase (gene 29), and genes involved in spermidine synthesis (genes 1, 2 and 21). Methyl group metabolism emerged as a central process during bacterial growth with algae. Consequently, to identify relevant genes that might have escaped annotation in the KEGG database, we conducted a manual search for bacterial genes involved in metabolizing methylated compounds, culminating in a comprehensive list of 83 genes (table S3 and fig. S4). Differential gene expression (DE) analysis indicated that the identified genes were markedly upregulated in the presence of algae, compared to bacterial pure cultures grown with glucose (Fig. 1B, and data S1 for DE results of all genes). To portray the possible routes by which bacteria utilize methyl groups during interactions with algae, we overlaid the reactions encoded by the methyl group metabolism genes with our DE results (Fig. 1C). This schematic overview of methyl group metabolism, also termed one-carbon metabolism, highlights that bacteria harvest, dissimilate, and assimilate methyl groups during interactions with algae, suggesting that methylated compounds are a relevant resource for algal-associated bacteria.

**Fig. 1:**
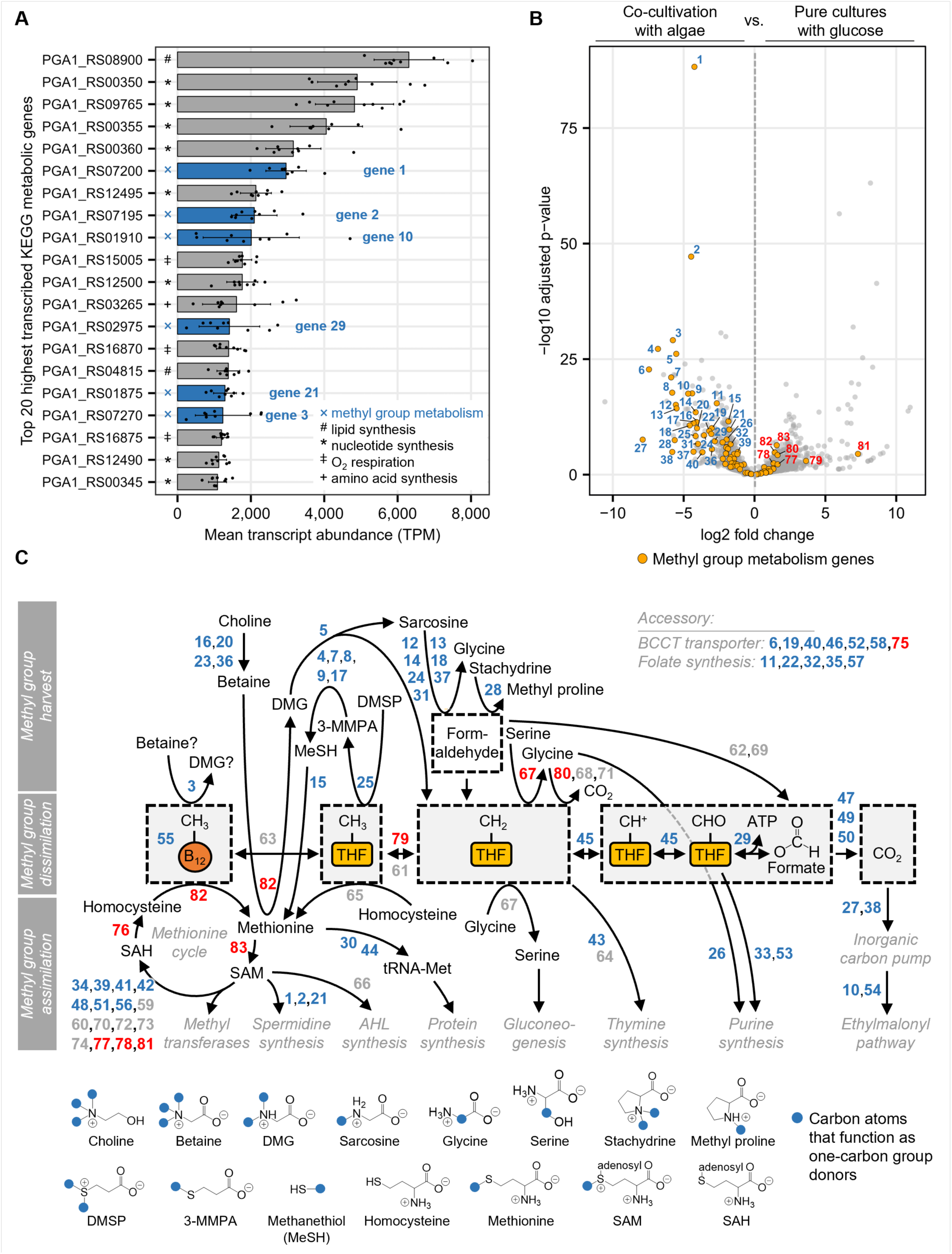
Identification of metabolic genes that are transcribed in *P. inhibens* bacteria during co-cultivation with *E. huxleyi* microalgae. (**A**) Co-culture RNA-sequencing revealed that bacterial genes involved in methyl group metabolism were among the highest transcribed KEGG metabolic genes in the presence of algae (blue bars). Bars represent the mean of nine algal-bacterial co-cultivation samples taken from three different time points (see fig. S1 for sampling points). Error bars depict standard deviations. Labels on the y-axis: RefSeq accession numbers. KEGG annotations of respective methyl group metabolism genes: *S*-adenosylmethionine decarboxylase (gene 1), trimethylamine—corrinoid protein Co-methyltransferase (gene 3), spermidine synthase (gene 2), crotonyl-CoA carboxylase/reductase (gene 10), formate—tetrahydrofolate ligase (gene 29), ornithine decarboxylase (gene 21). See data S1 for transcript abundances per sample. (**B**) Methyl group metabolism genes (orange dots) were upregulated in bacteria that were co-cultivated with algae (left; blue numbers) compared to bacteria in pure cultures that grew exponentially with glucose (right; red numbers). Samples used for differential gene expression analysis are presented in fig. S1. (**C**) Bacterial methyl group metabolism genes include reactions involved in harvesting, dissimilating and assimilating methyl groups from methylated donor compounds. Included are genes that are either directly involved in transforming methyl groups, or which are accessory and encode for transporters or tetrahydrofolate (THF) synthesis. Blue and red numbers indicate up-and downregulated genes, respectively (thresholds for coloring: adjusted *p*-value < 0.05; log2 fold change <-0.585 and > +0.585). Gene numbers are identical throughout the text and figures, with accession numbers and functional annotations listed in table S3. Blue dots in chemical structures represent carbon atoms (mainly methyl groups) that function as one-carbon group donors. BCCT—Betaine/Carnitine/Choline Transporter family.

### The methylated compound DMSP triggers the shortening of the bacterial lag phase

To investigate the influence of methylated compounds on bacteria, we conducted a series of growth experiments using the methylated compound dimethylsulfoniopropionate (DMSP) as a model substrate. DMSP is naturally produced by microalgae and was extensively studied in the context of bacterial metabolism and growth^33,34^. In our experiments, bacterial pure cultures were pre-cultivated with glucose as a sole carbon source until reaching stationary phase. Then, stationary phase bacteria were used to initiate fresh cultures which contained the same glucose medium, but that were additionally supplemented with DMSP (or with water as control). We found that micromolar concentrations of DMSP stimulated the growth of bacteria (Fig. 2A). Plotting the growth curves semilogarithmically revealed that the low levels of added DMSP affected specifically the bacterial lag period; lag times were shortened while growth rates were not affected (Fig. 2B and fig. S5A). We further quantified the lag phase shortening effect by calculating △lag times, which is the time difference between the onset of growth in control cultures, compared to supplemented cultures (Fig. 2B and fig. S5B). By analyzing △lag times under different DMSP concentrations, we found that lag times were shortened by up to 2.5 hours, and that already nanomolar concentrations of DMSP resulted in significantly lower lag times (Fig. 2C and fig. S5C). Interestingly, above 20 μM a higher concentration did not increase the delta lag phase. This suggests that there may be a threshold at which the demethylation machinery becomes saturated, leading to diminished sensitivity to DMSP, as previously suggested^35^.Our results show that in the presence of a utilizable carbon source such as glucose (provided at a concentration of 1 mM in our experiments), exposure to low levels of the methylated compound DMSP leads to the shortening of the bacterial lag phase.

**Fig. 2:**
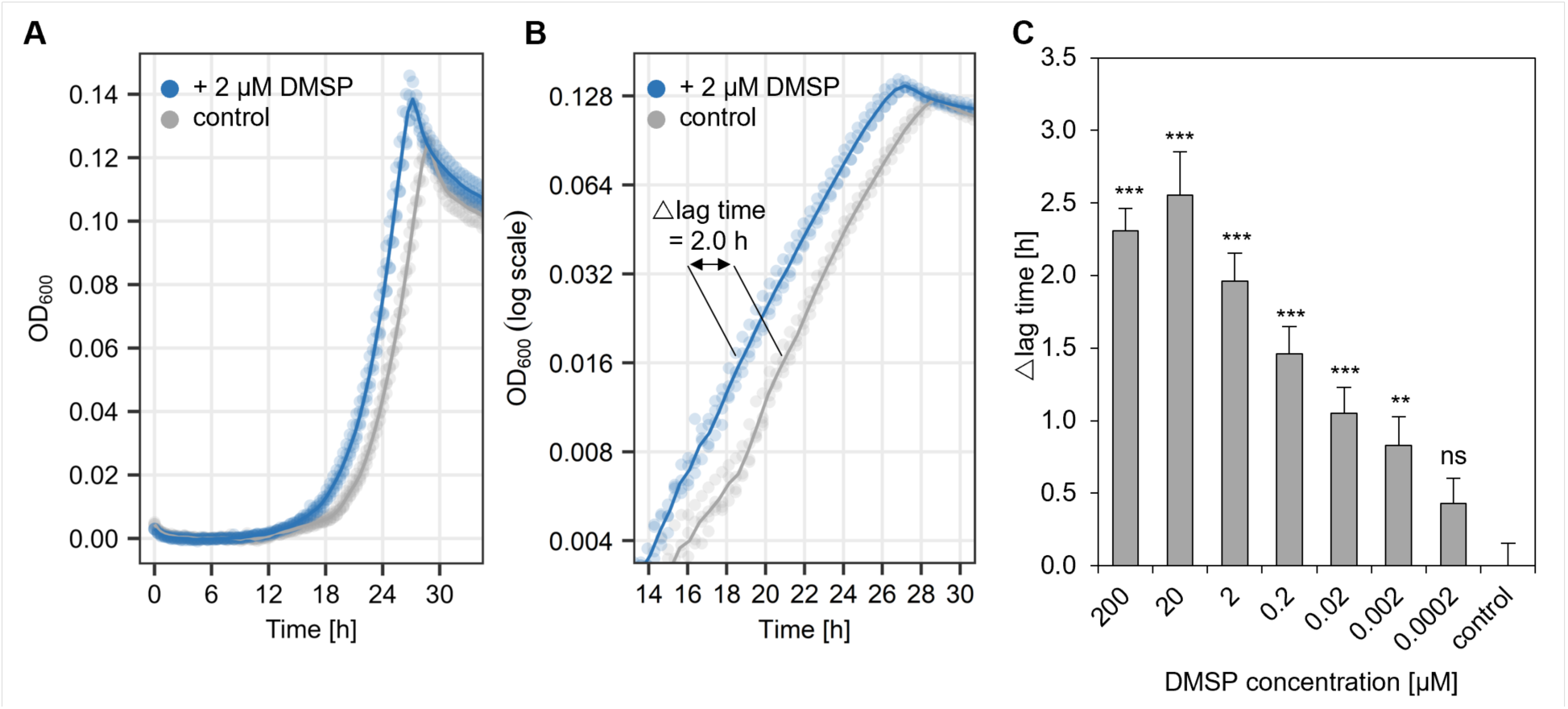
The methylated compound DMSP induces lag phase shortening in *P. inhibens* bacteria. (**A**) Micromolar concentrations of DMSP stimulated earlier growth of bacteria (blue) compared to control cultures supplemented with water (grey). (**B**) Semi-log2 transformed growth curves show that doubling times were unchanged, as evident from the identical slopes, while lag times were shortened. Lag phase shortening is expressed as the difference between mean lag times measured for supplemented cultures compared to control cultures (△lag time; see *Method details* for explanations). (**C**) Exposure to nanomolar concentrations of DMSP resulted in significant bacterial lag phase shortening. All growth experiments were conducted in artificial seawater medium (ASW_b_) with 1 mM glucose as carbon source, using glucose-grown, stationary phase pre-culture bacteria as inoculum. For each condition, four biological replicates were used (four wells of a 96-well plate were inoculated with the same pre-culture). In panels **A** and **B**, dots represent individual measurements for each of the four replicates per time point, and lines depict a smoothed average. Error bars in panel **C** indicate the standard deviation of the estimated difference between means. ANOVA and Dunnett’s test were used to identify significant differences between control and supplemented cultures (adjusted *p*-value thresholds: *** ≤ 0.001, ** ≤ 0.01, ns [not significant] > 0.05).

### Methyl groups are involved in lag phase shortening

To determine whether the methyl groups of DMSP are involved in lag phase shortening, we used analogues of DMSP that harbor different numbers of methyl groups (Fig. 3A and figs. S6-S7). We found that the extent of lag phase shortening correlated with the number of methyl groups per molecule, while non-methylated analogues did not have a measurable impact on the lag period (Fig. 3A and table S4 for statistical significance). This finding suggests that the methyl groups of DMSP drive lag phase shortening.

**Fig. 3:**
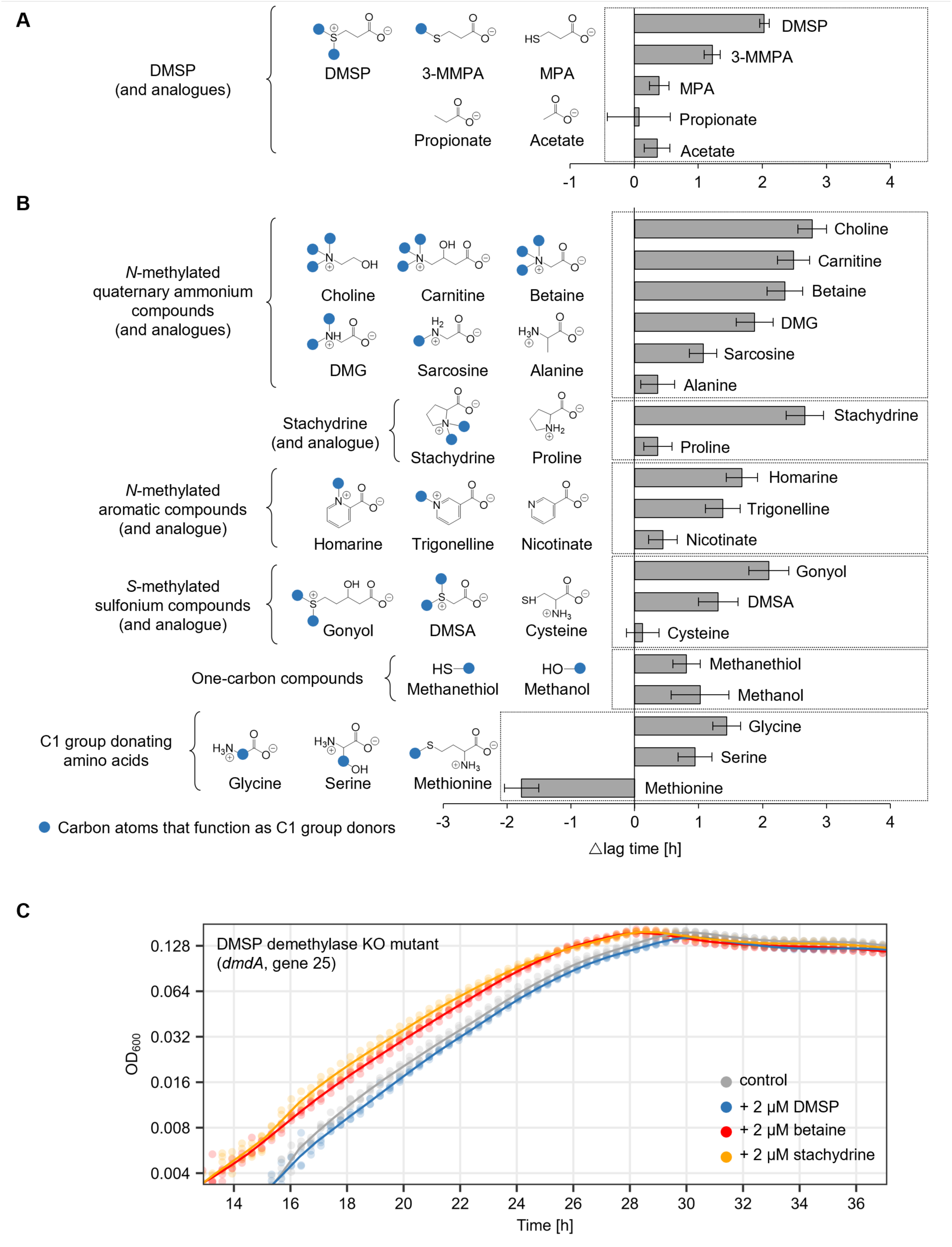
Methyl groups and other one-carbon moieties induce lag phase shortening in *P. inhibens* bacteria. (**A**) Growth experiments with analogues of DMSP (2 μM) revealed that methyl groups are required for lag phase shortening. The number of methyl groups per molecule correlated with the extent of lag phase shortening. (B) Growth experiments further revealed that multiple methylated compounds (2 μM) trigger lag phase shortening, while non-methylated analogues at the same concentration were less or not effective. In panels (A) and (B), blue dots in chemical structures represent carbon atoms (mainly methyl groups) that function as one-carbon group donors. Bars represent differences between mean lag times measured for supplemented cultures compared to control cultures (△lag time). Each condition was analyzed with three biological replicates, and error bars indicate standard deviation of the difference between estimated means. (C) Lag phase shortening was no longer evident in a *P. inhibens dmdA* knock-out mutant upon exposure to DMSP, while other methylated compounds (betaine and stachydrine) still triggered lag phase shortening. Dots represent individual measurements for four biological replicates, and lines depict a smoothed average.

To substantiate the role of methyl groups, we investigated whether other methylated compounds that are commonly produced by microalgae induce lag phase shortening (Fig. 3B). We selected methylated compounds with different chemical properties including *N*-methylated, *S*-methylated, small one-carbon compounds and amino acids (which are themselves not methylated but that can donate one-carbon groups to the cellular methyl group metabolism). All compounds were added at a concentration of 2 μM. The results show that supplementation with all tested methylated compounds resulted in shortening of the lag phase (Fig. 3B and table S4 for statistical significance), while the doubling times were largely unchanged (figs. S6-S7). The extent of lag phase shortening correlated with the number of methyl groups per molecule, in line with our results with the DMSP analogues (Fig. 3A). Interestingly, methionine was an exception and prolonged the lag phase. Growth inhibitions by methionine were reported for other bacteria and may result from feedback loops that regulate intracellular methionine concentrations^36^. In summary, our findings demonstrate that different methylated compounds prompt a reduction in the bacterial lag phase and suggest a correlation between the quantity of methyl groups present in a compound and the degree of lag reduction. Furthermore, the data also reveal that additional characteristics of the tested compounds, such as carbon count or the nature of chemical moieties (*N*-methylated versus *S*-methylated), do not influence the shortening of the lag phase. This underscores the centrality of methyl groups in the process of lag phase shortening.

### Harvesting the methyl group from a methylated compound is required for lag phase shortening

The involvement of methyl groups in lag phase shortening could be explained by three possible scenarios: 1) methyl groups are a limiting resource, 2) methylated compounds are sensed by receptor proteins that trigger a signal cascade, and 3) methylated compounds act as intracellular solutes with protective functions. To distinguish between these possibilities, we generated a *P. inhibens* knock-out (KO) mutant that is unable to harvest methyl groups from a respective donor compound. We deleted the *dmdA* gene (Fig. 1C; gene 25), which encodes for the well-characterized DMSP demethylase^37^. The deletion of the demethylase gene is expected to hamper the harvest of methyl groups from DMSP, while it is not expected to affect extracellular receptor binding or internalization of the compound. Our data show that DMSP-dependent lag phase shortening was abolished in the *dmdA* KO mutant (Fig. 3C). Importantly, the mutant still demonstrated lag phase shortening when supplemented with other methylated compounds (Fig. 3C). Interestingly, we observed a more pronounced difference in the reduction of the lag phase due to altered amounts of methyl groups on DMSP (Fig. 3A) compared to changes in the concentration of the added DMSP (Fig. 2C). While the molecular mechanisms underlying this difference are not fully understood, a possible explanation previously suggested is the insensitivity of DMSP catabolic enzymes to increasing levels of DMSP^35^. Additionally, DMSP and other methylated compounds are compatible solutes and are transported into the cell via high-affinity transport systems^38^. These transporters function effectively at low substrate concentrations but become saturated at higher concentrations, potentially contributing to diminished lag reductions with increasing DMSP concentrations. While the DMSP demethylase is a well-studied enzyme, its role in lag phase dynamics has been overlooked. These results suggest that methyl groups are a limiting resource during the lag phase and that bacteria can overcome this bottleneck by harvesting methyl groups from external donor compounds.

### Stoichiometric calculations highlight assimilatory methyl group requirements

Methyl groups may function as a limiting resource either by being assimilated as a one-carbon group (C1) substrate, or by being dissimilated for ATP generation (Fig. 1C). To explore the possible fate of methyl groups during the lag phase, we estimated the C1 group requirements per bacterial cell needed for one cell duplication. In this process, C1 groups are utilized as a substrate to synthesize cellular building blocks for biomass formation (i.e., purines, thymine, methionine and histidine; table S5). Based on numbers determined for exponentially growing *E. coli*^39^, we calculated that a single *P. inhibens* cell requires 190.1 amol of C1 groups to synthesize the building blocks needed for one cell duplication (table S5). If a freshly initiated *P. inhibens* culture, at a density of 25,000 CFU/mL, assimilates all available DMSP methyl groups during the lag phase, then the bacterial population requires 2.4 nM of DMSP to duplicate. Indeed, similar DMSP concentrations—2 nM—induced significant lag phase shortening (Fig. 2C).

Methyl groups can also undergo dissimilation to generate ATP, producing one ATP per methyl group (Fig. 1C; gene 29). According to the provided growth-associated maintenance cost (GAM) in the metabolic model of *P. inhibens*^40^, a single *P. inhibens* cell requires 23,800 amol ATP to conduct one cell duplication during exponential growth with glucose. If a freshly initiated *P. inhibens* culture (25,000 CFU/mL) were to rely entirely on the dissimilation of DMSP methyl groups during the lag phase to meet its ATP requirements, the bacterial population would need 298 nM of DMSP to fulfill 100% of the ATP costs for one duplication. In our experiments, 2 nM DMSP already induced significant lag phase shortening (Fig. 2C). This concentration can cover only 0.7% of the bacterial ATP needs. It is important to acknowledge that the presented values in this calculation are estimations and may likely underestimate bacterial requirements. However, it is noteworthy that the difference in assimilatory versus dissimilatory needs, which can be both fulfilled by methyl groups, spans two orders of magnitude. Despite potential biases in our estimations, this calculation highlights the considerable potential of methyl groups to alleviate assimilatory needs and accelerate the lag phase, rather than substantially affecting the high demand of dissimilatory processes. Based on these calculations, it appears more plausible that methyl groups are assimilated during the lag phase to cover C1 group requirements, while the required ATP is generated from the glucose in the medium.

### Methyl group assimilation genes are upregulated during the lag phase

To experimentally elucidate the intracellular fate of methyl groups, we analyzed the bacterial transcriptome in response to DMSP specifically during the lag phase. If methyl groups are indeed assimilated during the lag phase, functions associated with methyl group assimilation are anticipated to be transcriptionally regulated during this phase. The transcriptome of *P. inhibens* was previously demonstrated to be remarkably dynamic, revealing rapid responses occurring within minutes following treatment^41^. Therefore, RNA was sampled 15 min and 40 min after initiation of fresh cultures, using glucose-grown stationary phase bacteria as inoculum. The cultures contained either glucose as sole carbon source or were supplemented with DMSP (figs. S8-S10, tables S6-S7 and data S2). Our data show that the addition of DMSP resulted in the upregulation of 43 methyl group metabolism genes during the lag phase and the downregulation of 12 genes (figs. S11-S13)—an effect that was most pronounced 15 min post initiation of fresh cultures.

Specifically, genes involved in DMSP catabolism were upregulated (fig. S11; genes 4, 7, 8, 9, 17, 25), including the DMSP demethylase gene (*dmdA*: gene 25). The most upregulated genes were involved in synthesizing spermidine (fig. S11; genes 1, 2 and 21), which is a metabolite associated with cell growth^42^. The most downregulated gene was involved in methyl group oxidation—a regulation that may impede methyl group dissimilation (fig. S11; gene 79). Interestingly, DMSP demethylation (*dmdA*: gene 25), spermidine synthesis (*speD*: gene 1, *speE*: gene 2, and gene 21) and methyl group oxidation (*metF*: gene 79) are all linked via the methionine cycle (Fig. 4A). This link is further supported by the upregulation of *metK* (gene 83)—a gene that is involved in converting methionine into *S*-adenosylmethionine (SAM; Fig. 4A). In summary, transcriptome analyses suggest that DMSP methyl groups are assimilated during the bacterial lag phase through the methionine cycle.

**Fig. 4:**
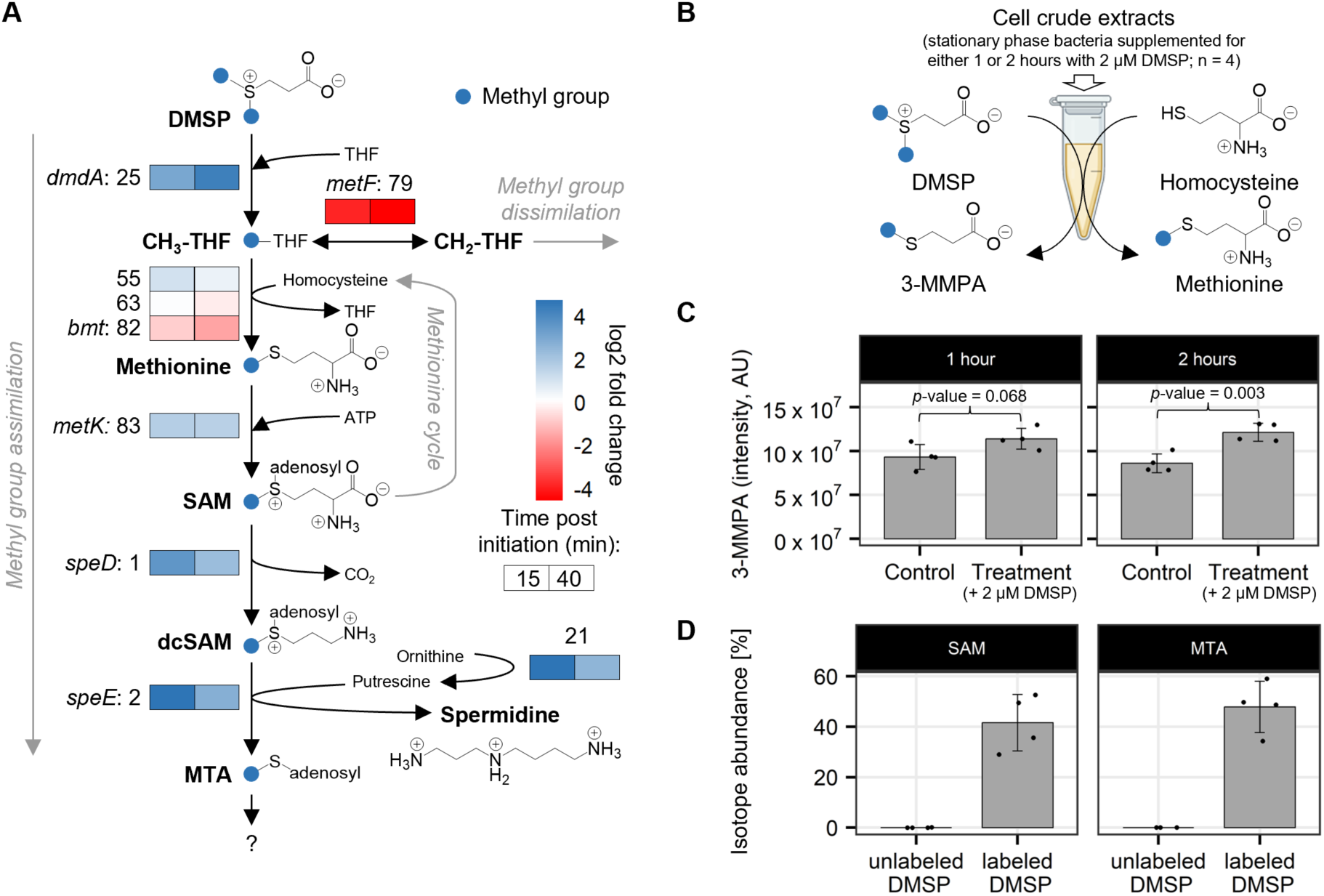
Transcriptional, metabolic and enzymatic response of *P. inhibens* bacteria towards DMSP during the lag phase. (**A**) The transcriptional response was analyzed in freshly initiated bacterial cultures that were supplemented with 50 µM DMSP, compared to control cultures without DMSP (all cultures contained 1 mM glucose; fig. S8). Gene expression changes (log2 fold change) are indicated as colored boxes that correspond to the age of the culture post initiation— 15 min (left box) and 40 min (right box). Colors show upregulation (blue) and downregulation (red) in response to DMSP. The three most upregulated methyl group metabolism genes were involved in spermidine synthesis (*speD*: gene 1, *speE*: gene 2, and gene 21; fig. S11). Spermidine synthesis utilizes *S*-adenosylmethionine (SAM), which is the main product of the methionine cycle. Likewise, genes involved in DMSP demethylation (*dmdA*: gene 25; fig. S11) and SAM synthesis (*metK*: gene 83; fig. S11) were highly upregulated in response to DMSP. A gene that mediates between methyl group assimilation and dissimilation was the most downregulated gene in response to DMSP (*metF*: gene 79; fig. S11). Numbers left to boxes or above designate gene numbers (see table S3 for accession numbers and functional annotations). RNA-sequencing was conducted using three biological replicates per condition. (**B**) Evaluating methionine synthesis activity in response to DMSP. Stationary phase bacteria were supplemented with either 2 µM DMSP or water as control (four biological replicates per condition) and sampled for cell disruption one and two hours after supplementation. Resulting cell crude extracts were used for *in vitro* enzymatic reactions (volume = 1.5 mL; 0.35 mg protein per reaction) with added homocysteine (2 mM) and DMSP (200 µM) to catalyze the formation of 3-*S*-methylmercaptopropionate (3-MMPA), which is the demethylated form of DMSP. Product formation was measured by LC-MS analysis. (**C**) Cell crude extracts of DMSP-supplemented stationary phase bacteria produced slightly more 3-MMPA after 1 hour (*p*-value = 0.068), and significantly more 3-MMPA after 2 hours (*p*-value = 0.003), compared to non-supplemented control cultures. Bars represent the mean of four biological replicates (dots), with error bars showing the standard deviation. The Student’s t-Test was used to calculate *p*-values. (**D**) Tracking the metabolic fate of DMSP methyl groups during the lag phase. Freshly initiated bacterial cultures were supplemented with 50 µM [^13^C-methyl]DMSP, in which both methyl groups are ^13^C-labeled. The intracellular formation of ^13^C-labeled metabolites was monitored 2 hours after initiation by LC-MS analysis—a time point at which higher DMSP demethylation activity was observed (shown in **C**). The method identified ^13^C-labeled SAM (isotope abundance for M+1: 41.6%) and ^13^C-labeled ^13^C-labeled *S*-methyl-5’-thioadenosine (MTA; isotope abundance for M+1: 47.9%).

### Methyl groups are harvested from DMSP and assimilated into bacterial metabolites during the lag phase through the methionine cycle

To verify the transcriptomics results suggesting the involvement of the methionine cycle in processing harvested methyl groups, we sought direct experimental validation. First, we examined whether bacteria in the stationary phase, before transitioning to the lag phase upon transferring to fresh media, possess the necessary enzymatic machinery to demethylate DMSP, thereby generating methyl groups for use in the methionine cycle. We therefore generated crude cell extracts from stationary phase bacteria supplemented with DMSP (or water as control) for one or two hours. These extracts were then incubated with the substrates required for DMSP demethylation and the utilization of methyl groups in the methionine cycle—homocysteine and DMSP (Fig. 4B). The formation of the demethylated product of DMSP, 3-MMPA, was measured using LC-MS analysis (Fig. 4C). Our results indicate that crude extracts from DMSP-supplemented stationary cells produced more 3-MMPA, with higher levels observed after a longer incubation period with DMSP (2 hours compared to 1 hour). This suggests that exposure to DMSP during the stationary phase influenced the enzymatic demethylation machinery that was subsequently activated in the crude extract (Fig. 4C). Importantly, analysis of the stationary bacteria used to prepare the crude extract showed significantly lower levels of 3-MMPA, regardless of DMSP supplementation (fig. S14). These findings confirm that stationary *P. inhibens* bacteria possess the enzymatic machinery necessary for DMSP demethylation. Furthermore, as bacteria transition from stationary phase to lag phase, this machinery has the potential to become activated and generate available methyl groups.

To follow the cellular fate of the harvested methyl groups, we wished to identify metabolites that are synthesized from the methyl groups of DMSP during the lag phase. Therefore, bacteria were supplemented with [^13^C-methyl]DMSP in which both methyl group carbons were labeled. In this experiment, glucose-grown stationary phase bacteria were used to initiate fresh cultures that contained 1 mM glucose as a sole carbon source, and to which either 50 µM [^13^C-methyl]DMSP or 50 µM unlabeled DMSP were added. Cells were harvested two hours after initiation of fresh cultures (fig. S8)—a time point after which DMSP demethylation activity was higher (Fig. 4C)— and then subjected to metabolite extraction and LC-MS analysis. Our results show that DMSP methyl groups were incorporated into SAM and *S*-methyl-5’-thioadenosine (MTA) with isotope abundances of 41.6% and 47.9%, respectively (values given for M+1 isotopologues with single ^13^C label; Fig. 4D, fig. S15 and data S4). SAM is formed via the methionine cycle and is required, among others, as a substrate for spermidine synthesis. MTA is a methylated side product of spermidine synthesis (Fig. 4A). The formation of ^13^C-labeled SAM and MTA supports that DMSP-derived methyl groups are assimilated via the methionine cycle. The percent of isotope abundance was similar for both compounds, suggesting that the majority of MTA is produced from SAM and therefore possesses a similar isotope abundance. We cannot exclude that methyl groups are also used for other processes, such as purine synthesis (fig. S12), however, results of [^13^C-methyl]DMSP LC-MS analysis are in accordance with the transcriptional upregulation of spermidine synthesis genes in response to DMSP (Fig. 4A). These results demonstrate that DMSP methyl groups are assimilated during the bacterial lag phase through the methionine cycle. These findings corroborate the conclusions drawn from our prior stoichiometric calculations and offer an empirical demonstration of methyl group assimilation through the methionine cycle for the synthesis of essential building blocks.

### Methylated compounds differentially shorten the lag phase of various bacteria that are associated with algae and plants

The capacity of *P. inhibens* bacteria to alleviate the cellular demands for methyl groups during the lag phase using algal methylated compounds enables them to initiate earlier growth. This rapid response holds potential significance in environmental settings when various starved bacteria encounter an algal host. Therefore, we hypothesized that the capability to utilize methylated compounds from an external source would be common among algal-associated bacteria. However, for this ability to be advantageous, it should be differentially demonstrated by various bacteria. Our findings indicate that certain algal-associated bacteria exhibit lag phase reduction in response to the methylated compounds DMSP and betaine, though not universally across all species. Additionally, the degree of lag phase reduction differs among bacteria and varies in response to distinct methylated compounds (Table 1, fig. S16).

**Table 1.** Bacteria that are naturally associated with algae and plants exhibit various lag reduction capabilities in response to DMSP and betaine. Values represent the △lag time of bacteria exposed to 2 µM of DMSP or betaine (compared to controls with water). Values in brackets indicate standard deviation of the difference between estimated means, using three biological replicates per condition. ND: Not detected.

| Bacterial species | Lag reduction [hours] |  |
| --- | --- | --- |
|  | DMSP | Betaine |
| **Algal-associated bacteria** |  |  |
| *Sulfitobacter pontiacus* iR4 | 1.2 (±0.1) | 1.6 (±0.1) |
| *Labrenzia* sp. DT40 | 4.2 (±0.3) | 5.6 (±0.3) |
| *Vibrio* sp. DT22 | 1.7 (±0.1) | 1.7 (±0.2) |
| *Phaeobacter* sp. DT104 | 2.7 (±0.9) | ND |
| *Pseudoalteromonas* sp. DT35 | ND | 1.4 (±0.1) |
| *Labrenzia* sp. DT112 | ND | ND |
| *Vibrio* sp. DT18 | ND | ND |
| *Marinomonas* sp. DT74 | ND | ND |
| **Plant-associated bacteria** |  |  |
| *Bacillus subtilis* 168 | 0.8 (±0.2) | 2.1 (±0.1) |
| *Pseudomonas aeruginosa* PAO1 | 1.2 (±0.7) | 1.5 (±0.2) |
| *Pseudomonas koreensis* PK | 1.5 ($\pm 0.6$) | ND |
| *Ewingella americana* EA | ND | ND |
| *Lelliottia amnigena* LA | ND | ND |
| *Enterobacter aerogenes* Ea | ND | ND |

The synthesis of methylated compounds is a hallmark of photosynthetic organisms, including both algae and plants^43,44^. Therefore, we tested whether plant-associated bacteria exhibit responses similar to algal-associated bacteria. Our results show that plant-associated bacteria exhibit different lag phase shortening in response to betaine and DMSP (Table 1, fig. S17). We extended our investigation by analyzing the genomes of the tested bacterial strains to determine whether they possess the 83 methyl metabolism genes that exhibited differential expression in *P. inhibens* (Figure 1). However, our analysis did not reveal a correlation between the presence of these genes in the examined genomes and the documented shortening of lag phase (fig. S18). The absence of correlation may be attributed to technical factors such as inadequate annotation and the unavailability of precise genomes for these specific strains. For example, our findings reveal that *Vibrio* bacteria shorten their lag phase in response to DMSP. However, the key DMSP demethylase gene, *dmdA* (gene 25), is absent from the *Vibrio* genome (fig. S18). While previous studies have demonstrated DMSP catabolism in *Vibrio*, the specific enzymes involved remain unidentified^45^. These uncertainties emphasize the need for further investigation to elucidate the underlying mechanisms responsible for lag phase shortening across diverse bacterial strains. Our data demonstrate that various bacteria that interact with photosynthesizing organisms can commence earlier growth using methylated compounds of their host. The exact response of bacteria to specific compounds may be influenced by the natural environments in which these bacteria have evolved. These findings illustrate how previously unrecognized bacterial physiologies, such as the mechanism for lag reduction, become apparent when bacteria are studied within an ecological context encompassing factors like starvation and a host.

## Discussion

### The lag phase in the lab and in the environment

More than a century ago, the scientific community identified a non-replicative period when starved bacterial cultures encountered fresh nutrients. This period was termed the “lag phase”, and it was observed and characterized in controlled laboratory cultures. However, our understanding of the molecular details of this phase remains limited. The challenge lies primarily in the technical demands of traditional laboratory methodologies, which require high bacterial densities, while the lag phase occurs at the onset of cultures when bacterial populations are exceedingly sparse. Recent advancements have provided insights into the cellular events characterizing the lag phase^15^, yet much is unknown about this period in both model and environmental strains.

The complexities of studying the lag phase using laboratory cultures underscore the difficulty in understanding and characterizing this phenomenon in natural microbial populations. Although direct observation of bacterial lag phase in natural settings is impractical, analogies can be drawn between the laboratory-based lag phase and potential occurrences in the environment. In natural settings, bacteria commonly face periods of nutrient scarcity and exist in a phase of nongrowth^46^. Upon encountering a nutrient source, bacteria can resume growth. The transient phase between growth and nongrowth can represent the lag phase. Furthermore, heterotrophic bacteria rely on organic compounds from photosynthetic organisms. These bacteria experience fluctuations in available resources due to diurnal and seasonal photosynthesis variations. The fluctuations lead to cycles of nutrient scarcity followed by surges of organic compounds, potentially triggering a bacterial lag phase during these transitions.

Laboratory model systems cannot fully replicate real-world conditions but can help in assessing the possible environmental relevance of the lag phase. Model system can simulate critical factors such as low bacterial density, nutrient-deprived conditions, low substrate concentrations, and relevant microbial interactions. The utilization of model systems, including simple co-cultures and more complex synthetic communities, can significantly contribute to our understanding of how the lag phase might manifest within natural ecosystems.

### Challenges in determining the exact duration of the lag phase

Investigating the lag phase requires the precise definition of its initiation and conclusion. In laboratory settings, the onset of the lag phase is easily determined at the starting of the culture during the inoculation stage. However, pinpointing the termination of the lag phase, specifically when bacteria transition into the growth phase, presents a challenge. Typically, at the population level, the common method for demarcating the lag phase involves plotting bacterial growth semilogarithmically and identifying the intersection between the initial inoculum level and the extrapolated slope of the logarithmic growth phase. At the single-cell level, the lag phase culminates when a cell reaches its first division. Recent advances in single-cell analysis have unveiled the heterogeneity within microbial populations, with distinct lag phase phenotypes observed within these populations^47^.

The common techniques for monitoring bacterial population growth and define their lag phase are further complicated when studying environmental strains. Various environmentally relevant bacteria, such as *P. inhibens*, have mechanisms that enable them to attach onto algal hosts. When multiple bacteria adhere to an algal cell, conventional methods like live counts and optical density measurements may be inadequate for accurately estimating bacterial cell numbers. Therefore, technological advancements to enable the tracking of single bacterial cells while they are attached to their host will be imperative for precision in such studies. In addition, better understanding of the cellular processes that occur during the lag phase may help in developing metabolic and genetic markers for the study of populations and single cells.

### Spermidine synthesis and the potential link to translation initiation during the lag phase

Analyses of *P. inhibens* bacteria in the context of their algal host, revealed the centrality of methyl group metabolism during algal-bacterial interactions, and specifically methyl group assimilation during the lag phase. Our data highlights that spermidine synthesis, a downstream metabolite of the methionine cycle, can be a notable cellular sink for methyl groups during the lag phase. In DMSP-supplemented bacteria, pronounced transcriptional upregulation was observed of the *metK* gene that is involved in transforming methionine into SAM^48^—the precursor for spermidine synthesis (Fig. 4A). Additionally, the three most upregulated methyl group metabolism genes were associated with the conversion of SAM into spermidine—a pathway that yields MTA as a side product (genes 1, 2 and 21; fig. S11). Moreover, ^13^C-labeled methyl groups were incorporated into SAM and MTA during the lag phase (Fig. 4D). Spermidine is a polycationic polyamine that interacts with negatively charged macromolecules and has various cellular functions^42,49^. The benefits of polyamines are often associated with protein synthesis^50^, and externally supplemented polyamines can increase protein synthesis by up to 2-fold^49^.

Additionally, our results suggest that translation initiation, the crucial step in protein synthesis, could be a destination of methyl groups during lag phase. Translation initiation heavily relies on formylated methionine (fMet) as the initial amino acid. The production of fMet requires at least two C1 groups, emphasizing the importance of methyl groups during the lag phase. In our transcriptome analyses, the *metG* gene that is responsible for charging tRNA^Met^ with methionine, was markedly upregulated in DMSP-supplemented *P. inhibens* bacteria during the lag phase (gene 44; fig. S11). Moreover, we found that various genes involved in adding C1 modifications to tRNA and rRNA components of the translation apparatus were upregulated during the lag phase (e.g., genes 39, 41, 42, 43; fig. S11 and table S3). Thus, while requiring further substantiation, our data suggest a link between methyl group assimilation and translation initiation.

Since protein synthesis is a primary cellular activity during the preparatory lag phase, the functional connection between translation initiation and spermidine synthesis in the context of methyl metabolism warrants additional investigation.

### Bacterial lag phase reduction in an environmental context

The metabolism of algal-associated bacteria is fueled by algal-produced metabolites. Thus, the bacterial diet naturally includes methylated compounds that are a hallmark for photosynthetic organisms^23–25,43^. Our findings suggest that algal-associated bacteria can efficiently harvest and assimilate methyl groups from algal donor compounds. While bacteria are capable of synthesizing methyl groups from glucose, harvesting methyl groups from host metabolites can provide a cost-effective metabolic shortcut. In the marine environment, expediting the transition from lag phase to exponential growth is critical to compete with other marine bacteria for valuable algal resources. Specifically, we observed that algal produced methylated compounds shortened the lag time by up to 2.5 h in *P. inhibens* (Fig. 2C), which is similar to the doubling time of these bacteria (fig. S7). Consequently, under optimal conditions, bacteria may yield twice as many cells in response to pulses of algal metabolites, compared to bacteria that are not adapted to harvest methyl groups from external methylated compounds. Maximizing the efficiency of resource utilization when resources are available presents a selective advantage to bacteria^51^. Given that different algal hosts synthesize and release a unique composition of metabolites^52,53^, including methylated compounds^54,55^, the differential bacterial response may represent host-specific adaptations. It remains to be determined how closely the spectrum of methylated compounds generated by specific microalgae corresponds to the demethylation abilities of the associated bacteria. Nevertheless, our study offers an initial demonstration of the tunability of the lag phase, highlighting the existing gap in our understanding of non-growing bacteria and the mechanisms facilitating their transition into growth.

## Materials and Methods

### Key Resources Table

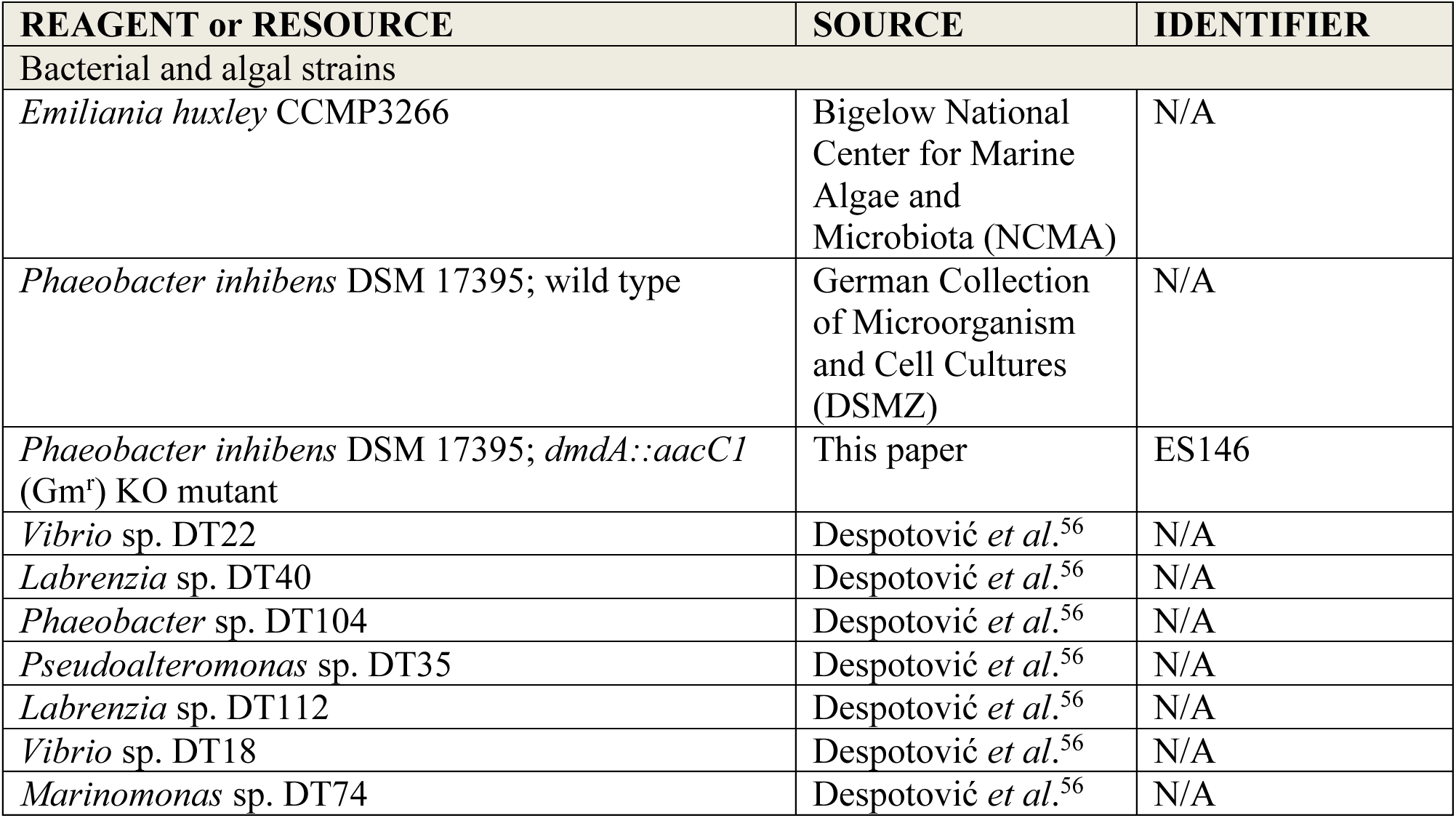
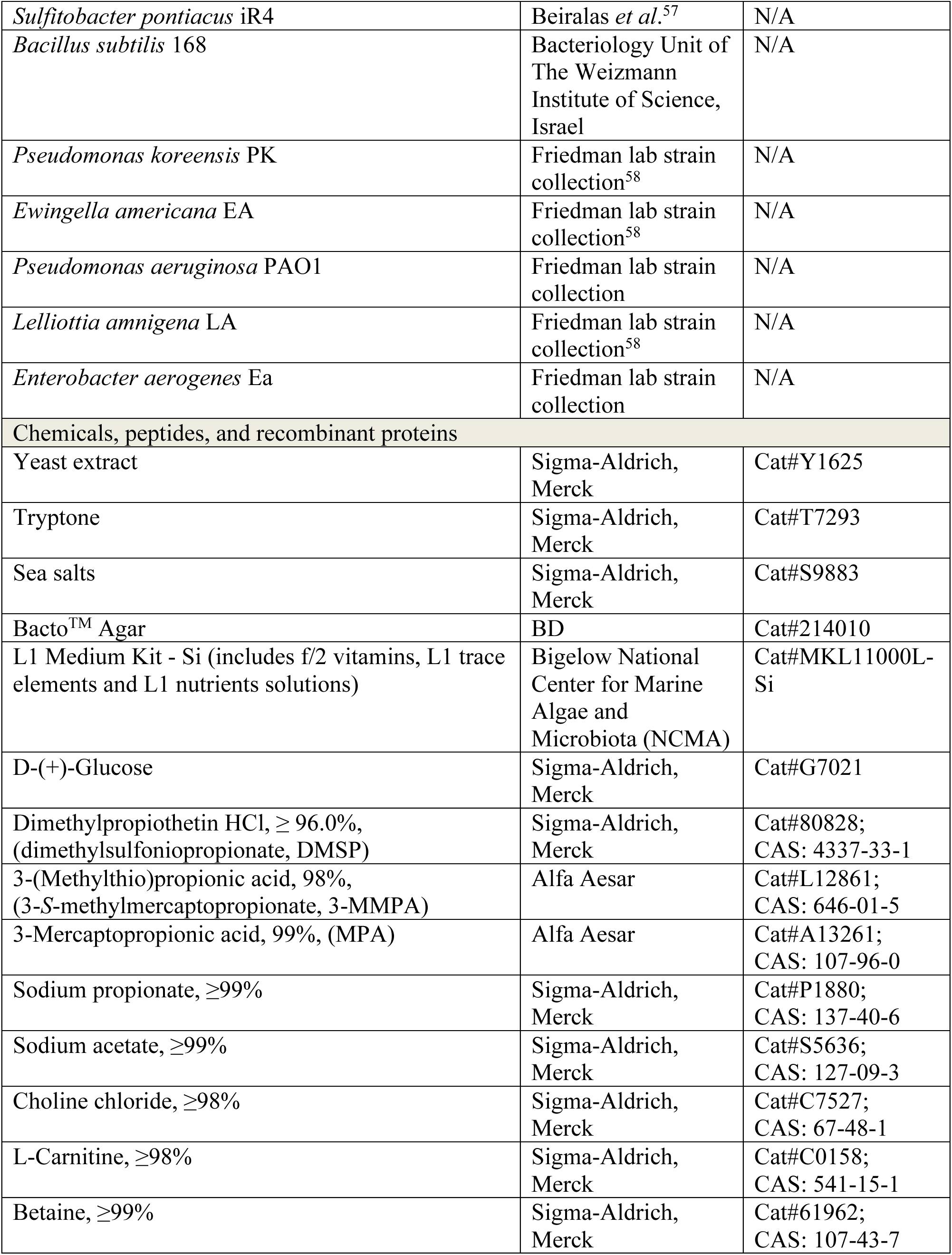
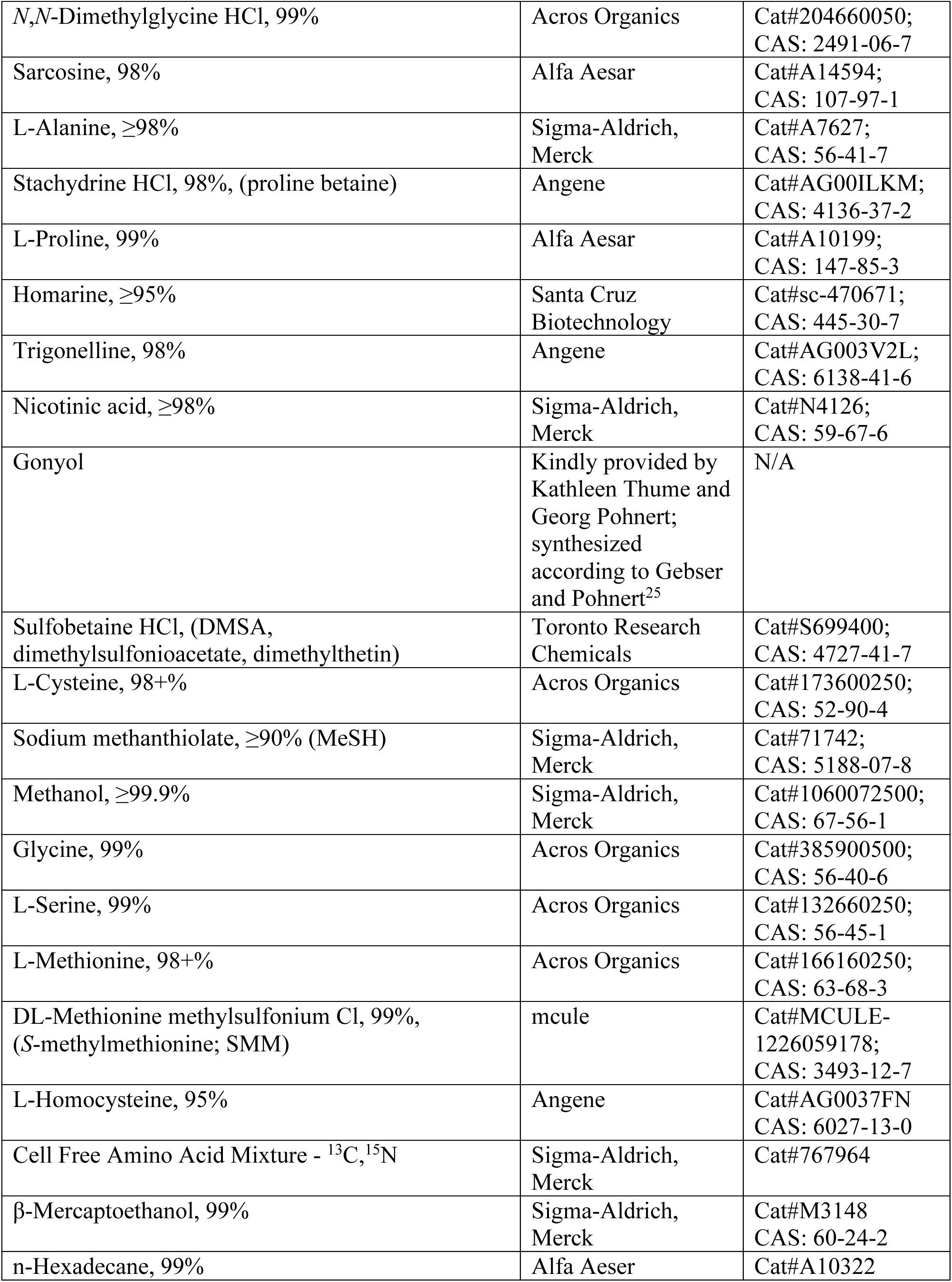
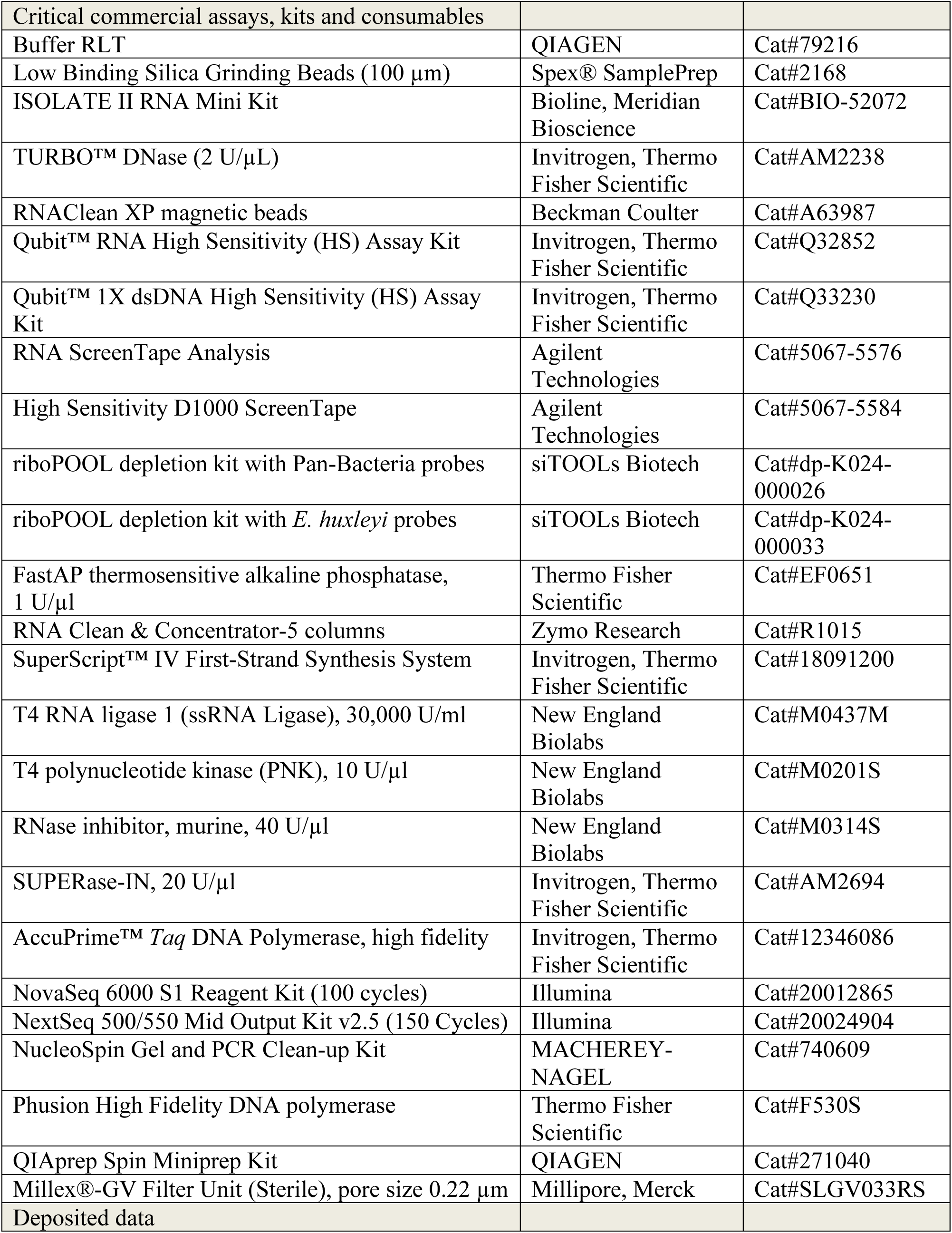
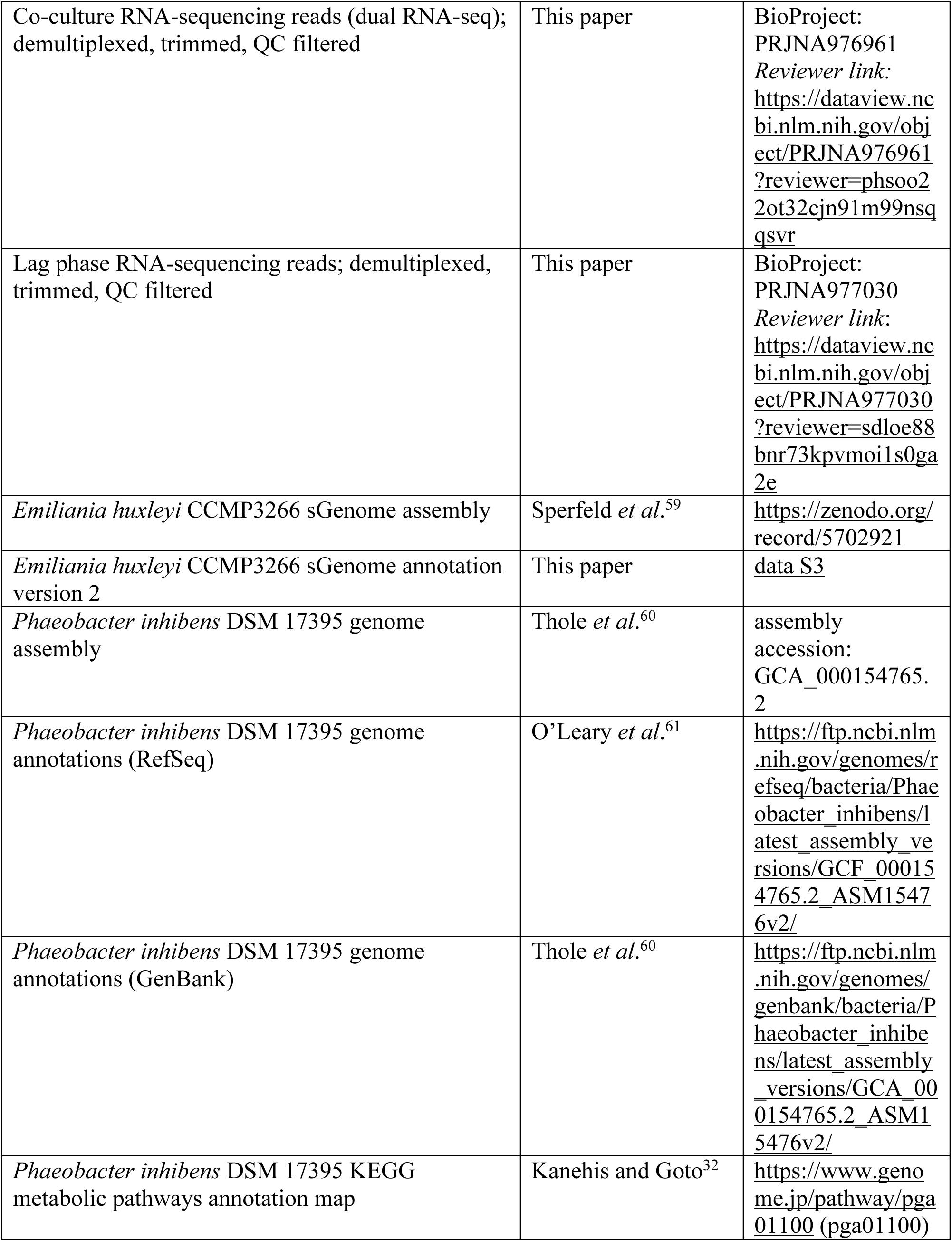
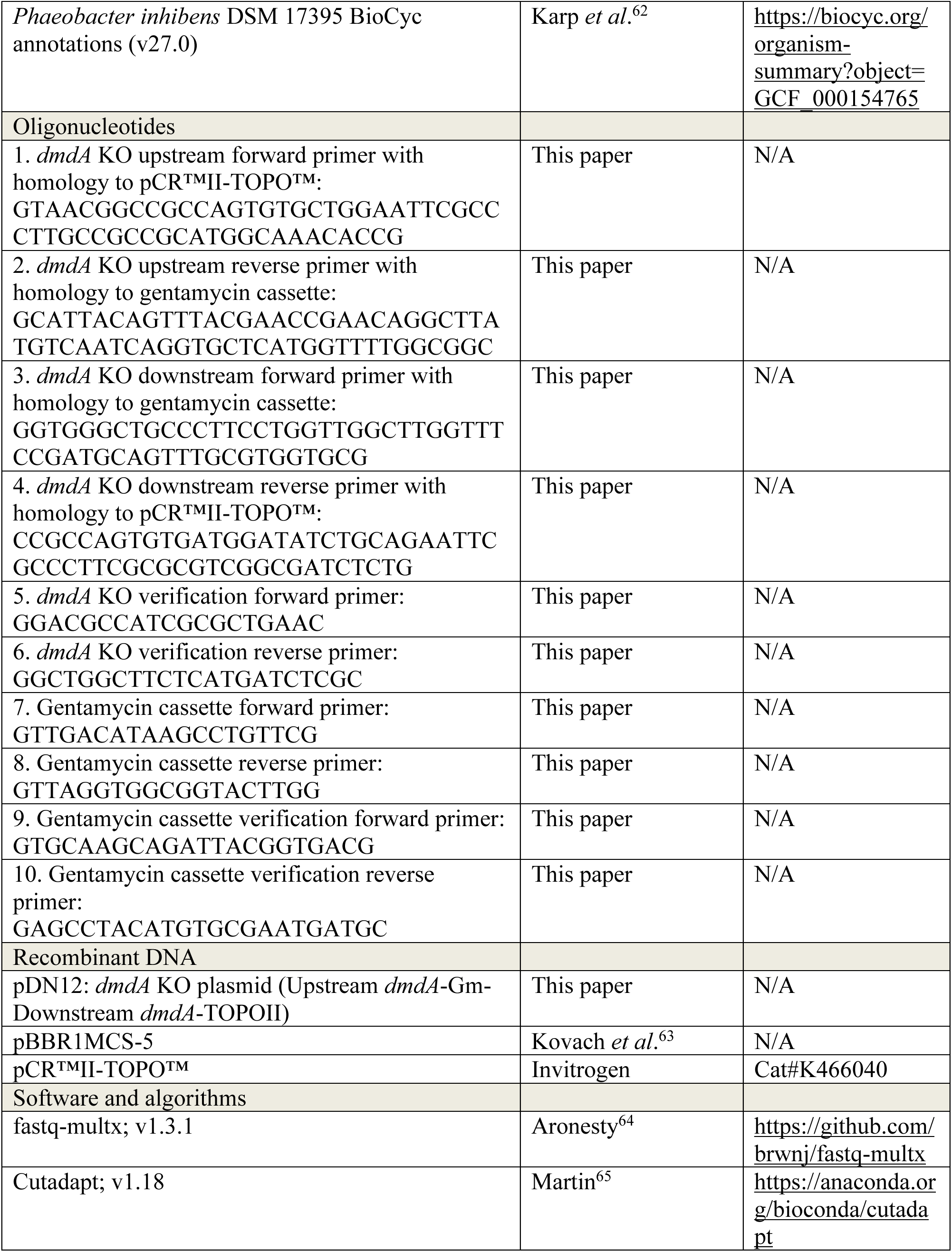
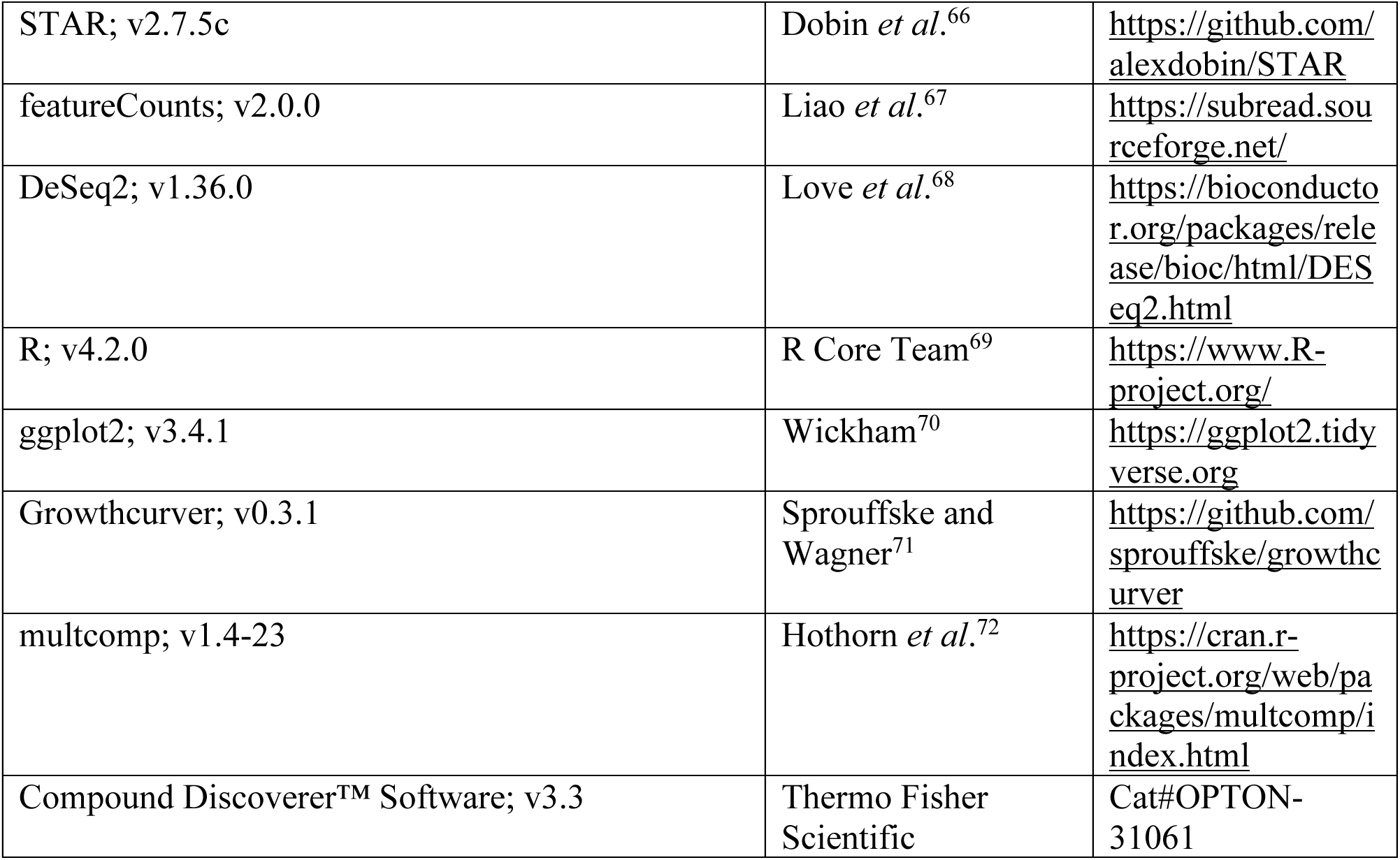

### Microalgae, media composition and general culture conditions

The algal strain *Emiliania huxleyi* CCMP3266 was purchased as an axenic culture from the National Center for Marine Algae and Microbiota (Bigelow Laboratory for Ocean Sciences, ME, USA). Absence of bacteria in axenic algal cultures was monitored periodically both by plating on ½ YTSS agar plates (yeast extract, 2 g/L; trypton, 1.25 g/L; sea salts, 20 g/L; agar, 16 g/L) and under the microscope. The absence of bacteria was additionally confirmed by recent transcriptome sequencing^59^. Algae were maintained in ASW_a_ medium (artificial seawater) at 18°C with a light/dark cycle of 18/6 h and an illumination intensity of 150 mmoles/m^2^/s. The ASW_a_ medium was based on the protocol of Goyet and Poisson^73^ and contained mineral salts (NaCl, 409.41 mM; Na_2_SO_4_, 28.22 mM; KCl, 9.08 mM; KBr, 0.82 mM; NaF, 0.07 mM; Na_2_CO_3_, 0.20 mM; NaHCO_3_, 2 mM; MgCl · 6 H_2_O, 50.66 mM; CaCl_2_, 10.2 mM, SrCl_2_ · 6 H_2_O, 0.09 mM), f/2 vitamins (thiamine HCl, 100 μg/L; biotin, 0.5 μg/L; vitamin B_12_, 0.5 μg/L), L1 trace elements (Na_2_EDTA · 2H_2_O, 4.36 mg/L; FeCl_3_ · 6H_2_O, 3.15 mg/L; MnCl_2_ · 4H_2_O, 178.1 μg/L; ZnSO_4_ · 7H_2_O, 23 μg/L; CoCl_2_ · 6H_2_O, 11.9 μg/L; CuSO_4_ · 5H_2_O, 2.5 μg/L; Na_2_MoO_4_ · 2H_2_O, 19.9 μg/L; H_2_SeO3, 1.29 μg/L; NiSO_4_ · 6H_2_O, 2.63 μg/L; Na_3_VO_4_, 1.84 μg/L; K_2_CrO_4_, 1.94 μg/L) and L1 nutrients (NaNO_3_, 882 μM; NaH_2_PO_4_ · 2H_2_O, 36.22 μM). The components were dissolved in Milli-Q water (IQ 7003; Merck, Darmstadt, Germany) and the pH adjusted to 8.2 with HCl. Stock solutions of f/2 vitamins, L1 trace elements and L1 nutrients were purchased from the Bigelow Laboratory for Ocean Sciences (Boothbay, ME, USA).

### Marine bacteria and media composition

The bacterial strain *Phaeobacter inhibens* DSM 17395 was purchased from the German Collection of Microorganism and Cell Cultures (DSMZ, Braunschweig, Germany). The bacterial strain *Sulfitobacter pontiacus* iR4 was isolated from a xenic culture of *Emiliania huxleyi* CCMP1516^57^. The bacterial strains *Vibrio* sp. DT22, *Labrenzia* sp. DT40, *Phaeobacter* sp. DT104, *Pseudoalteromonas* sp. DT35, *Labrenzia* sp. DT112, *Vibrio* sp. DT18, and *Marinomonas* sp. DT74 were isolated from the Mediterranean Sea and Red Sea^56^, and belong to the Tawfik lab marine bacteria collection, which is now part of the Segev lab strain collection at the Weizmann Institute of Science, Israel. The marine bacteria were cultivated in ASW_b_ medium (identical to the ASW_a_ medium, but with additional 5 mM NH_4_Cl and 33 mM Na_2_SO_4_; f/2 vitamins were omitted, and glucose added as indicated), filtered sea water medium (Mediterranean Sea water enriched with L1 trace elements, f/2 vitamins, and L1 nutrients in the same concentrations as used for ASW_a_) or in ½ YTSS medium (yeast extract, 2 g/L; trypton, 1.25 g/L; sea salts; 20 g/L), as indicated. Bacterial isolates were kept at-80°C in ½ YTSS medium with 20% glycerol for long-term storage.

### Non-marine bacteria and media composition

The bacterial strains *Pseudomonas koreensis* PK, *Pseudomonas aeruginosa* PAO1, *Ewingella americana* EA, *Lelliottia amnigena* LA, and *Enterobacter aerogenes* Ea were kindly provided by Jonathan Friedman (The Hebrew University of Jerusalem, Israel)^58^. The *Bacillus subtilis* strain 168 was obtained from the Bacteriology Unit of the Weizmann Institute of Science, Israel. The strains were cultivated in M9 medium (Na_2_HPO_4_ · 7H2O, 64 g/L; KH_2_PO_4_, 15 g/L; NH_4_Cl, 5 g/L; NaCl, 2.5 g/L) with 20 mM glucose. Bacterial isolates were kept at-80°C in LB medium (tryptone, 10 g/L; yeast extract, 5 g/L; NaCl, 10 g/L NaCl) with 20% glycerol for long-term storage.

### Co-culture RNA-sequencing: Experimental design

To analyze the transcriptome of *P. inhibens* DSM 17395 bacteria during co-cultivation with the *E. huxleyi* CCMP3266 algae, a dual RNA-sequencing approach was applied that allows probing prokaryotic and eukaryotic RNA in the same samples, without separating cells by e.g. filtration^74^. For cultivation, 250 mL Erlenmeyer flasks with 60 ml ASW_a_ medium were inoculated with either both organisms (co-cultures), or with one of the two organisms (pure cultures). Specifically, 24 flasks were inoculated with both organisms (20,000 *E. huxleyi* cells and 2,000 *P. inhibens* CFUs; bacteria were pre-cultivated for 48 h in ASW_b_ medium with 2 mM glucose and f/2 vitamins, and added to algae four days after inoculation). Another 24 flasks were inoculated with a pure culture of algae (20,000 *E. huxleyi* cells). All 48 flasks were sealed with aluminum foil and cultivated in standing cultures in a growth room at 18 °C under a light/dark cycle of 16/8 hr. Illumination intensity during the light period was 150 mmoles/m^2^/s. At each sampling point, three random flasks were sacrificed per condition (3 algal-bacterial co-culture flasks, and 3 algal pure culture flasks) and sampled to enumerate algal cells, bacterial colony forming units (CFU) and to harvest cells for RNA extraction (starting from day 0; see fig. S1 for sampling points). Nine additional replicate flasks were inoculated with pure cultures of bacteria (2,000 *P. inhibens* CFUs; 0.88 mM NH_4_Cl and 2 mM glucose were added to the ASW_a_ medium to facilitate bacterial growth). The nine bacterial flasks were cultivated under the same conditions as the flasks with algae. Three flasks were repeatedly sampled to enumerate bacterial CFUs, while three flasks were sacrificed per sampling point to harvest cells for RNA extraction (see fig. S1 for growth curves and RNA sampling points).

Algal cell numbers were determined for each sampled flask as technical triplicates, using a CellStream flow cytometer (Merck, Darmstadt, Germany). Bacterial CFU numbers were determined for each flask as technical duplicates, using serial plating on ½ YTSS medium agar plates. To harvest cells for RNA extraction, 50 mL culture was centrifuged for 5 min at 18 °C and 3,220 g (5810 R swing-out centrifuge; Eppendorf, Hamburg, Germany) and the supernatant was removed by vacuum suction. Cell pellets were resuspended in 450 µl RLT buffer (QIAGEN, Hilden, Germany) with 1% β-mercaptoethanol, transferred into a 2 mL screw cap tube with 300 mg of 100 µm acid washed Low Binding Silica Beads (SPEX SamplePrep, Metuchen, NJ, USA), plunged into liquid nitrogen and stored at-80 °C for one month until use.

### Co-culture RNA-sequencing: RNA extraction and library preparation

RNA extracts were generated from 30 selected cell pellets (3 biological replicates per sampling point) that originate from algal-bacterial co-cultures, axenic algae and bacteria grown in pure culture with 2 mM glucose (see fig. S1 for sampling points and sample designations). Cell pellets were disrupted by 5 min bead beating at 30 s^−1^ in a mixer mill MM 400 (Retsch, Haan, Germany). RNA was extracted from disrupted cells using the ISOLATE II RNA Mini Kit, which integrates a step for an on-column DNase treatment (Meridian Bioscience, OH, USA). The extracted RNA was subjected to a second DNAse treatment using the TURBO DNase kit (Thermo Fisher Scientific, Waltham, MA, USA) and subsequently purified with 2x concentrated RNAClean XP magnetic beads (Beckman Coulter, Brea, CA, USA). RNA quantity and integrity was evaluated with the Qubit RNA HS Assay (Invitrogen, Thermo Fisher Scientific, Waltham, MA, USA) and the Tapestation RNA ScreenTape analysis (Agilent Technologies, Santa Clara, CA, USA), respectively (fig. S2). Ribosomal RNA was depleted using the riboPOOL rRNA depletion kit in combination with Pan-Bacteria probes and custom probes generated for *E. huxleyi* rRNA from the nucleus, plastid and mitochondrion (siTOOLs Biotech, Planegg, Germany). For ribosomal RNA depletion, 0.9 µg of RNA was used as an input with 50 pmol of each of the two depletion probes. After rRNA depletion, RNA was purified with 2x concentrated RNAClean XP magnetic beads (Beckman Coulter, Brea, CA, USA). Subsequent library preparation was performed as described by Avraham *et al*.^74^, which is a stranded, poly(A)-independent protocol that targets total RNA. Briefly, depleted RNA was fragmented by heat, treated with DNase and FastAP, and then 3’-tagged by ligating the barcoded adapters 1 to 30^74^ (RNase-free HPLC purified; ordered from Integrated DNA Technologies, Coralville, IA, USA). After barcoding, we implemented a calibration sequencing run to quantify the relative abundance of bacterial RNA in each sample. This quantification allows to determine the optimal sample-specific sequencing depths, and prevents over-sequencing of samples that contain either dead algae, or only one of the two organisms. For the calibration RNA-sequencing run, we proceeded with half of the barcoded RNA and stored the rest at-80 °C. All 30 samples were pooled in an equivolume and purified with RNA Clean & Concentrator-5 columns (Zymo Research, Irvine, CA, USA). RNA was reverse transcribed with the SuperScript IV First-Strand Synthesis System (Invitrogen, Thermo Fisher Scientific, Waltham, MA, USA), using the AR2 primer^74^. The resulting cDNA was 3’-ligated to the 3Tr3 adaptor^74^ (HPLC purified; ordered from Integrated DNA Technologies, Coralville, IA, USA). The 3’-ligated cDNA was then amplified with 9 PCR cycles (see fig. S19A for PCR cycle optimization). Finally, the amplified cDNA was purified with a double-sided clean-up step (right: 0.5x, left: 1.5x; RNAClean XP beads, Beckman Coulter, Brea, CA, USA), followed by a left-sided clean-up step (0.7x). The size distribution of the final calibration sequencing library is shown in fig. S19B. After cleanup, the library was sequenced on a NextSeq 500 instrument with a 150 cycles Mid Output Kit (Illumina, San Diego, CA, USA) in paired-end mode (Read 1: 88 bp, Read 2: 78 bp, Index 1: 0 bp, Index 2: 0 bp). Sequencing data were analyzed for the relative abundance of algal and bacterial RNA per sample, using bioinformatics methods described in the section below. The library preparation protocol was then repeated for the final co-culture RNA-sequencing run, starting from the stored, barcoded RNA. The barcoded RNA was pooled in a way that co-culture samples with algae are expected to yield equal amounts of bacterial sequencing reads (fig. S1; co-culture day 04, 06 and 09; expected bacterial feature counts per sample: 250,000-500,000), while fewer sequencing reads were allocated to algal pure cultures (axenic day 04, 06, 09 and 11/12; expected algal feature counts per sample: 40,000,000), co-cultures with bleached algae (co-culture day 11/12; expected algal feature counts per sample: 25,000,000) and bacterial pure cultures (glucose 1 and glucose 2; expected bacterial feature counts per sample: 5,000,000). After reverse transcription and ligation, the 3’-ligated cDNA was amplified with 11 PCR cycles (see fig. S20A for PCR cycle optimization). The final co-culture RNA-sequencing library (fig. S20B) was deep-sequenced on a NovaSeq 6000 instrument with a 100 cycles S2 Kit (Illumina, San Diego, CA, USA) in paired-end mode (Read 1: 64 bp, Read 2: 54 bp, Index 1: 0 bp, Index 2: 0 bp; table S1). The expected distribution of algal and bacterial sequencing reads per sample was confirmed as described below (see table S2 for resulting algal and bacterial feature counts per sample).

### Co-culture RNA-sequencing: Data analysis

Illumina raw reads were demultiplexed with fastq-multx^64^, allowing one mismatch per barcode (v1.3.1;-m 1). Raw reads were then quality filtered and trimmed with cutadapt^65^ (v1.18;-a AGATCGGAAGAGCACACGTCTGAACTCCAGTCAC-A AGATCGGAAGAGCGTCGTGTAGGGAAAGAGTGT-g T{20}-A A{20}--times 2--nextseq-trim=20-m 20--pair-filter=both; see table S2 for summary; reads where deposited under the BioProject accession: PRJNA976961). The quality filtered and trimmed reads were mapped to a reference fasta file that included the previously generated “synthetic genome” assembly (sGenome) of *E. huxleyi* CCMP3266^59^ and the genome of *P. inhibens* DSM 17395 (accession: GCF_000154765.2). Read mapping was conducted with STAR^66^ (v2.7.5c;--outSAMattributes All--alignIntronMax 5000--alignMatesGapMax 5000--outSAMtype BAM SortedByCoordinate--outFilterMultimapNmax 100--outSAMmultNmax 1). For compatibility with STAR, empty lines generated by cutadapt were replaced with N’s (sed’s/^$/N/’). Reads that mapped to *E. huxleyi* features were counted using featureCounts from the subread package^67^ (v2.0.0;-p-C-a Ehux3266_sGenome_v2.gff-F GTF-O--fraction; see data S3 for annotation file). Of note, a slightly modified version of the previously generated *E. huxleyi* CCMP3266 annotation file was used with featureCounts^59^, in which plastid and mitochondrion annotations were replaced by public annotations (accessions: JN022705.1 and JN022704.1, respectively). For compatibility with featureCounts, the NH:i attribute of STAR bam files was changed to “1” (sed-r’s/NH:i:[0-9]+/NH:i:1/g’; STAR reports the number of alternative alignments in the NH:i attribute, which interferes with featureCounts fractional counting). Reads that mapped to *P. inhibens* features were counted with the same featureCounts options but using the *P. inhibens* RefSeq annotations. The output of featureCounts is summarized in table S2. To evaluate the efficiency of ribosomal RNA depletion, also non-rRNA features were summarized (table S2), in which algal rRNA from the nucleus (*E. huxleyi* CCMP3266 sGenome gene ID: G8853), the plastid (Plas_rrn5, Plast_rrnL, Plast_rrnS) and the mitochondrion (Mito_rrnL, Mito_rrnS), as well as bacterial 23S rRNA (PGA1_RS00070, PGA1_RS12745, PGA1_RS14220, PGA1_RS16070), 16S rRNA (PGA1_RS00055, PGA1_RS12760, PGA1_RS14235, PGA1_RS16085) and 5S rRNA (PGA1_RS00075, PGA1_RS12740, PGA1_RS14215, PGA1_RS16065) were excluded.

Bacterial gene transcription was analyzed only in co-culture RNA-sequencing samples that contained *P. inhibens* (table S2; fig. S1). Since sample-specific rRNA depletion efficiencies may skew results, bacterial rRNA features were removed from the feature count table for downstream analysis. To analyze sample variability, bacterial non-rRNA feature count results were normalized for sequencing depth and RNA composition using the DESeq2’s median of ratios method, followed by a regularized log transformation and principle component analysis^68,75^ (v1.36.0; blind = TRUE; ntop = 500; R version 4.2.0; fig. S3). DESeq2 was further used to identify differentially expressed (DE) genes by comparing samples of bacteria growing with algae (co-culture day 04, 06 and 09; figs. S1 and S3) to samples of bacteria growing exponentially in pure cultures (glucose 1; figs. S1 and S3). The apglm method was used to shrink log2 fold changes and account for overweighing of features with low counts and high variability^76^. Shrunken log2 fold changes and adjusted *p*-values are given in the data S1. To compare absolute transcript abundances between genes and samples, bacterial feature counts were converted to transcripts per kilobase million (TPM); a normalization method that accounts for gene length and sequencing depth (data S1). To identify the 20 highest transcribed metabolic genes, the *P. inhibens* feature table (data S1) was filtered for genes that are part of the KEGG metabolic pathways annotation map^32^ (pga01100; https://www.genome.jp/pathway/pga01100). The resulting 840 KEGG metabolic genes were filtered for the 20 genes with the highest mean transcript abundances in samples with algae (co-culture day 04, 06 and 09).

To identify methyl group metabolism genes in *P. inhibens*, publicly available functional annotations were added to the feature table (data S1). These annotations include automatically generated NCBI RefSeq annotations^61^, partially curated GenBank submitter annotations^60^, KEGG annotations^32^, as well as links to BioCyc^62^ and the unifying UniProt Archive (UniParc^77^; links redirect to e.g. pre-computed domain annotations at the InterPro database^78^). Features of *P. inhibens* were then filtered for relevant genes, based on differential expression patterns (adjusted *p*-value: ≤ 0.05; log2 fold change: ≤-0.585 and ≥ 0.585; data S1). The DE-filtered genes were manually curated by sourcing the public annotations, which identified methyl group metabolism genes that are involved in transporting and catabolizing methylated compounds, in harvesting, dissimilating or assimilating one-carbon groups, or in synthesizing tetrahydrofolate (THF). Also genes that are involved in the ethylmalonyl-CoA pathway were included, which replenishes the TCA cycle by capturing CO_2_ during growth on the methylated compound DMSP^79^. Complementary to this approach, methyl group metabolism genes were identified by revising literature on one-carbon metabolism^80,81^, and searching for homologues genes in *P. inhibens* using nucleotide and protein BLAST^82^. In the latter approach, genes were considered as relevant if they exhibited ≥ 50 mean TPM in at least one of the sampling points. The resulting list, which includes 83 curated *P. inhibens* methyl group metabolism genes, is given in table S3. For clarification, genes were excluded from the list that did not meet the above-described cut-off filters, such as a putative DMSP cleavage enzyme (*dddP*; PGA1_RS09315) that was poorly transcribed in all samples (< 50 TPM), and non-regulated methyltransferases.

### Growth response in bacteria to supplements

To test the effect of methylated compounds on the growth of *P. inhibens* wild-type and KO mutant with deleted *dmdA* gene (Fig. 3C), bacteria were streaked from glycerol stocks on ½ YTSS medium agar plates and incubated at 30 °C for 48 hours. After incubation, a single colony was used to inoculate a pre-culture with 10 mL ASW_b_ medium (1 mM glucose, 100 mL Erlenmeyer flasks). The pre-culture was cultivated at 30 °C with 130 rpm shaking for 48 hours. After 48 hours, the pre-culture was in the stationary phase and exhibited OD_600_ ≈ 0.1. The stationary phase pre-culture was used to inoculate the main cultures (1 mL ASW_b_ medium, 1 mM glucose, start OD_600_ = 0.00001 [based on OD_600_ in pre-culture and applied dilution factor]), which contained supplements as indicated (mainly 2 µM of a methylated compound; listed in Key Resources Table). Supplements were prepared as 50 mM stock solutions in Milli-Q water and sterilized by filtration with syringe filters (Millex-GV, 0.22 µm, PVDF, Merck, Darmstadt, Germany). Each inoculated main culture was divided into three or four different wells (biological replicates) of a 96-well microtiter plate (150 µl main culture per well) and overlaid with 50 µl hexadecane to prevent evaporation^83^. Cultivation of the main cultures was conducted at 30 °C in an Infinite 200 Pro M Plex plate reader (Tecan Group Ltd., Männedorf, Switzerland) with alternating cycles of 4 min shaking and 16 min incubation. Optical density measurements were conducted at 600 nm following the shaking step.

The same growth protocol was used to analyze growth responses when supplemented with 2 µM of the methylated compounds DMSP or betaine, or water as control in the marine bacteria *Vibrio* sp. DT22, *Labrenzia* sp. DT40, *Phaeobacter* sp. DT104, *Pseudoalteromonas* sp. DT35, *Labrenzia* sp. DT112, *Vibrio* sp. DT18, and *Marinomonas* sp. DT74 (fig. S16). A slightly modified protocol was used for *S. pontiacus* iR4 marine bacteria, in which ASW_b_ medium was replaced by filtered seawater medium (fig. S16). To study the responses to 2 µM DMSP or betaine, the plant-associated bacteria *B. subtilis* 168, *Pseudomonas koreensis* PK, *Pseudomonas aeruginosa* PAO1, *Ewingella americana* EA, *Lelliottia amnigena* LA, and *Enterobacter aerogenes* Ea, were grown in M9 minimal media supplemented with 20 mM glucose (fig. S17).

### Growth data analysis

For data analysis, microtiter plate absorption values were converted into OD_600_ values by multiplying measured values with a factor of 3,8603. This factor was determined by comparing the absorption (at 600 nm) of a culture sample in a spectrophotometer using a 1 cm cuvette (Ultrospec 2100 Pro, Biochrom, Cambridge, UK) with the absorption in the microtiter plate reader (150 µl culture + 50 µl hexadecane per well). The background absorption, which is the mean absorption during the non-growth phase with stable OD_600_ values (≈ 2-8 hours), was subtracted from each individual well. Results were plotted in R (v4.2.0) using ggplot2 (v3.4.1). For visual aid, smoothing lines were added that average the replicates for each condition (geom_smooth; method: loess). Doubling times and lag times were calculated for each replicate using the Growthcurver package^71^ (v0.3.1). For this package, background subtracted OD_600_ values were used as an input that span the time from the lowest to the highest measured absorptions. Growthcurver was used to fit the OD_600_ values into a logistic equation that describes the population size *N*_*t*_ (OD_600_) at time *t*. The output of Growthcurver includes the modelled growth rate *r*, the carrying capacity *K*, and the initial population size *N*_0_.

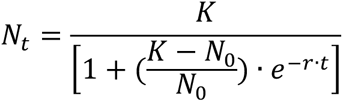

To calculate lag times, we formulated a definition in which lag times equal the time *t* at which *N*_*t*_ (OD_600_) reaches an absorption of 0.01. Using this definition, the logistic equation of Growthcurver was rearranged to *t* and solved with *N*_*t*_ = 0.01:

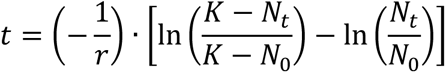

To calculate △lag times, we first determined the mean lag time for each condition. Then, the mean lag times of supplemented treatment conditions were subtracted from the mean lag time of the control condition. Thus, the resulting △lag times are a measure for the effect of supplements on the length of the lag phase. To determine the standard deviation of the estimated difference between means *σ*_*M*1-*M*2_, a formula was used that incorporates the standard deviation for the lag time of the treatment condition (*σ*_1_), the control condition (*σ*_2_), and the number of replicates *n* per condition:

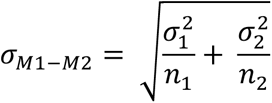

Doubling times were calculated by solving the logistic equation of Growthcurver with *N*_*t*_ = 2*N*_0_. The R package multcomp^72^ (v1.4-23) was used to identify significant differences between lag times and doubling times. Specifically, an ANOVA model (aov function) was fitted to the two growth parameters, followed by a Dunnett’s test (glht function).

### Knock-out mutants with deleted dmdA

Primers and plasmids used to generate *P. inhibens* knock-out (KO) mutants are listed in the Key Resources Table. PCR amplification and restriction-free cloning^84^ were performed using the Phusion High Fidelity DNA polymerase (Thermo Fisher Scientific, Waltham, MA, USA). PCR products were purified with the NucleoSpin Gel and PCR Clean-up kit (MACHEREY-NAGEL, Düren, Germany). Plasmids were purified with the QIAprep Spin Miniprep Kit (QIAGEN, Hilden, Germany).

To generate KO mutants for the *dmdA* gene (accession: PGA1_RS19020; gene 25) we first PCR-amplified ∼1,000 bp upstream and downstream regions of the respective loci (primers 1-4). A gentamycin resistance gene was amplified from the plasmid pBBR1MCS-5 (primers 7-8). The PCR products (upstream region + gentamycin resistance + downstream region) were then assembled and cloned into the pCR™II-TOPO™ vector (Invitrogen, Thermo Fisher Scientific, Waltham, MA, USA) using restriction-free cloning^84^. This resulted in the KO plasmid pDN12 (*dmdA*). For transformation, 10 µg of KO plasmid were added to 300 µl of competent *P. inhibens* cells (prepared by the method of Dower *et al*.^85^), and electroporated with a pulse of 2.5 kV (MicroPulser, Bio-Rad Laboratories, Hercules, CA, USA). Electroporated cells were recovered in 2 mL ½ YTSS medium for 12 hours at 30 °C and transferred to selective ½ YTSS medium agar plates with 30 µg/ml gentamycin. To screen for transformants, single colonies were used as template for PCR reactions (primers 5-6 and 9-10). Gene knock-outs were additionally verified in single-cell clones by Sanger sequencing.

### Stoichiometric calculations

To estimate the amount of methyl group requirements per *P. inhibens* cell during growth on glucose, we identified metabolites that are built from one-carbon (C1) groups^86,87^, and are thus C1 sinks. The focus was on C1 sinks that are used as building blocks to construct macromolecules such as DNA, RNA and proteins, and for which quantitative data are available in *E. coli*^39^. We identified seven C1 sinks in *P. inhibens* (ATP, GTP, dATP, dGTP, dTTP, histidine and methionine; table S5). Not included were serine, phosphatidylcholine, spermidine, macromolecule methylations and low abundant metabolites. Serine is an important C1 group donor in cells (Fig. 1; gene 67), however, serine synthesis from glucose does not consume C1 groups. Phosphatidylcholine has a choline backbone with three *N*-methyl groups and is produced by many bacteria^88^. It was reported that some *Phaeobacter* species produce phosphatidylcholine^89^, however, only minor amounts of this lipid were detected in *P. inhibens* DSM 17395 during growth with glucose^90^. The metabolite spermidine was excluded due to uncertainties about methyl group recycling. The bacterium *P. inhibens* possesses all genes required to synthesize spermidine by first decarboxylating *S*-adenosylmethionine (SAM) to *S*-adenosyl 3-(methylsulfanyl)propylamine (dcSAM), and then condensing dcSAM with putrescine to spermidine (Fig. 4A). Importantly, the *S*-methyl group of dcSAM is not directly incorporated into spermidine but remains attached to a side product of spermidine synthesis, which is *S*-methyl-5’-thioadenosine (MTA; Fig. 4A). While the *S*-methyl group of MTA is recycled in other bacteria via the methionine salvage pathway, this pathway appears to be incomplete in *P. inhibens*. Thus, the incomplete methionine salvage pathway may result in the loss of one methyl group per synthesized spermidine molecule. However, considering that a yet unknown methionine salvage pathway may exist in *P. inhibens*^91^, spermidine was not included as a C1 sink (table S5). Methylated forms of the macromolecules DNA^92^, RNA^93^ and proteins^94^ were also not included because to the best of our knowledge, no robust data exist on the quantity of macromolecule methylations in the cell. Lastly, not included were low-abundant C1 sinks such as pantothenate (required for CoA synthesis) or NAD/NADP, which represent negligible fractions of the bacterial biomass^39^. After identifying C1 sinks in *P. inhibens*, we gathered data on the amount of the respective sinks present in *E. coli* (table S5). The values represent *E. coli* cells grown at 37 °C in aerobic glucose minimal medium and at a doubling time of 40 min^39^. A single *E. coli* cell has a dry weight of 280 fg under these conditions, which we used to calculate the amount of C1 sinks present per cell (amol building block/cell; table S5). We then multiplied the amount of C1 sinks per cell with the net amount of C1 groups that are incorporated into each of the sinks (amol C1 group/cell; table S5). By summing up the results for the individual C1 sinks, we estimated that a single *E. coli* cell requires 190.1 amol C1 groups to synthesize its building blocks. To calculate the C1 group requirement per *P. inhibens* cell, we assumed that *P. inhibens* and *E. coli* produce the same amounts of the respective C1 sinks. This assumption is corroborated by the fact that *P. inhibens* is, like *E. coli*, a copiotrophic bacterium^95^ with a single cell length of 1-2 µm^89^. To estimate ATP costs, we retrieved the growth-associated ATP maintenance costs (GAM) from the metabolic model of *P. inhibens*, which is given as 85 mmol ATP per gram dry weight^40^. Based on a dry weight of 280 fg per cell^39^, this equals 23,800 amol ATP per *P. inhibens* cell.

### Lag phase RNA-sequencing

To analyze transcriptional changes induced by DMSP during the lag phase of *P. inhibens*, a glycerol stock of these bacteria was streaked on a ½ YTSS medium agar plate and incubated at 30 °C for 48 hours. After incubation, a single colony was used to inoculate four pre-culture flasks, each containing 100 ml ASW_b_ medium with 1 mM glucose. The four pre-cultures were incubated for 48 hours (30 °C, 130 rpm shaking), resulting in stationary phase bacteria (OD_600_ ≈ 0.1). The stationary pre-cultures were then pooled and washed by centrifugation (5 min, 3,220 g and 4 °C) using a 5810 R swing-out centrifuge (Eppendorf, Hamburg, Germany). After centrifugation, the supernatant was removed by vacuum suction, and pellets were re-suspended in ASW_b_ medium (without glucose). A second washing step was performed, after which pellets were resuspended in 30 mL ASW_b_ medium (without glucose) to concentrate the cells. The concentrated cells (OD_600_ = 0.82) were used to inoculate six main culture flasks with DMSP (50 µM) and six flasks without DMSP. Each flask contained 60 mL ASW_b_ medium (1 mM glucose, pre-warmed at 30 °C, start OD_600_ = 0.01). The 12 inoculated main culture flasks were incubated at 30 °C with 130 rpm shaking. Six flasks (three flasks with 50 µM DMSP and three flasks without DMSP) were harvested 15 min after inoculation for RNA extraction. The other six flasks were harvested 40 min after inoculation. Separate cultivation experiments were conducted to confirm that DMSP stimulates growth under the applied conditions (fig. S8). The protocol used for cell harvest and RNA extraction is described in the section *Co-culture RNA-sequencing: RNA extraction and library preparation* with the adaptation that cells were centrifuged at 4 °C. Results of the RNA extraction are shown in fig. S9. All 12 extracted RNA samples were subjected to RNA library preparation as described in the same section, but with minor modifications. RNA fragmentation was conducted with 300 ng RNA per sample, and rRNA depletion was performed after the pooling step, using 100 pmol Pan-Bacteria probes. Of note, the 3’-ligated cDNA was amplified with 12 PCR cycles (see fig. S21A for PCR cycle optimization). The resulting lag phase RNA-sequencing library (fig. S21B) was sequenced on a NextSeq 500 instrument with a 150 cycles Mid Output Kit (Illumina, San Diego, CA, USA) in paired-end mode (Read 1: 89 bp, Read 2: 79 bp, Index 1: 0 bp, Index 2: 0 bp; table S6 and S7).

The pipeline described in the section *Co-culture RNA-sequencing: Data analysis* was used to analyze lag phase RNA-sequencing data. Quality filtered and trimmed sequencing reads (deposited under the BioProject accession: PRJNA977030) were mapped to a reference fasta file that contained only the genome of *P. inhibens* DSM 17395 (accession: GCF_000154765.2). After generating the *P. inhibens* non-rRNA feature count table, the data was used as input with the DESeq2 PCA plot function to confirm close clustering of replicates (fig. S10). DESeq2 was also used to compare gene expression in DMSP-supplemented samples with control samples 15 min and 40 min after inoculation (fig. S11). The DE results were included in the TPM normalized non-rRNA feature count table (data S2).

### DMSP demethylation from cell crude extracts

Enzymatic DMSP demethylation was measured in cell crude extracts using a method adapted from published protocols^96,97^. A glucose-grown, stationary phase pre-culture of *P. inhibens* bacteria was used to inoculate eight main culture flasks (100 ml ASW_b_ medium, 5.5 mM glucose). The main cultures were then cultivated for four days (30 °C, 130 rpm) until reaching stationary phase (OD_600_ ≈ 1.0). Four stationary phase main culture flasks were supplemented with 2 µM DMSP (100 µl from a sterile 2 mM DMSP stock solution, dissolved in Milli-Q water) while the other four flasks were kept as non-supplemented controls (adding 100 µl Milli-Q water). Cells were harvested 1 hour and 2 hours after supplementation by centrifugation (5 min, 3,220 g, 4 °C), and pellets were resuspended in 1 mL Solution A (100 mM Hepes-KOH pH 7.5, 1 mM DTT; all steps were conducted on ice, using pre-cooled solutions at 4 °C). Resuspended cells were then washed twice with Solution A by centrifugation (5 min, 10,000 g, 4 °C). To disintegrate cell membranes, resuspended cells were transferred into 2 mL screw-capped tubes filled with 600 mg of 100 µm acid-washed Low Binding Silica Beads (SPEX SamplePrep, Metuchen, NJ, USA), plunged into liquid nitrogen, thawed, and disrupted by 5 min bead beating at 30 s^−1^ in a mixer mill MM 400 (Retsch, Haan, Germany). Disrupted cells were centrifuged (5 min, 15,000 g, 4 °C) and the supernatant transferred into a new tube, resulting in the cell crude extract. Protein concentrations were measured using the Protein Assay Dye Reagent Concentrate (Bio-Rad Laboratories, Hercules, CA, USA), and diluted to 0.7 mg/mL with Solution A. Enzymatic reactions were performed by mixing 0.5 mL of 0.7 mg/mL diluted protein (cell crude extract) with 1 mL of Solution B (20 mM Hepes-KOH pH 7.5, 3 mM homocysteine, 300 µM DMSP, 2 mM DTT), resulting in 0.35 mg protein per 1.5 mL reaction mix. The reaction mixes were incubated for 1 hour at 30 °C. After incubation, 500 µl of the enzymatic reactions were transferred into fresh tubes, plunged in liquid nitrogen, and lyophilized for subsequent metabolite extraction and LC-MS analysis, as described in the section *Lag phase LC-MS analysis with [13C-methyl]DMSP* (but in negative ionization mode). The identification of 3-MMPA was confirmed by using an authentic standard.

### Lag phase LC-MS analysis with [^13^C-methyl]DMSP

To analyze DMSP methyl group assimilation during the lag phase, [^13^C-methyl]DMSP, which contained two ^13^C-labeled methyl groups, was synthesized in the Organic Synthesis Unit, Chemical Research Support, Weizmann Institute of Science, Israel. Labeled [^13^C-methyl]DMSP was prepared according to Dickschat *et al*.^98^. The synthesis was carried out by the acid-catalyzed addition of [^13^C_2_]dimethyl sulfide to acrylic acid, using the experimental conditions described in the literature for the deuterium labelled compound. The ^13^C-labelling was confirmed by ^1^H NMR, based on the appearance of the methyl signals as a doublet centered at δ = 3 ppm, the typical ^1^H-^13^C J coupling constant of 145 Hz and the mass spectrum showing the molecular ion at *m/z* 137. The NMR and the mass spectra did not show a signal for unlabeled DMSP.

Growth conditions for experiments with [^13^C-methyl]DMSP were comparable to those applied for lag phase RNA-sequencing. Briefly, pre-cultures of *P. inhibens* were cultivated for 48 h (100 ml ASW_b_ medium, 1 mM glucose) and washed twice by centrifugation. A concentrated pre-culture pool of stationary phase bacteria was used to inoculate eight main culture flasks (60 ml ASW_b_ medium, 1 mM glucose; start OD_600_ = 0.01)—four flasks contained 50 µM [^13^C-methyl]DMSP and four flasks contained 50 µM non-labeled DMSP. The eight main culture flasks were harvested two hours after inoculation by centrifugation (10 min, 7.000 rpm, 4° C; see fig. S8 for growth curves), and cell pellets were subsequently snap-frozen in liquid nitrogen. The selection of the two-hour time point was informed by the observation of higher DMSP demethylation activity in crude extracts of stationary phase bacteria supplemented with 2 µM DMSP (Fig. 4B).

Extraction and analysis of polar metabolites were performed as previously described in Malitsky *et al*.^99^ and Zheng *et al*. with some modifications. The bacterial pellets were extracted with 1 mL of a pre-cooled (-20°C) homogenous methanol:methyl-tert-butyl-ether (MTBE) 1:3 (v/v) mixture. The tubes were vortexed and sonicated for 30 min in an ice-cold sonication bath (taken for a brief vortex every 10 min). Then, 0.5 mL of a UPLC-grade water:methanol (3:1, v/v) solution with 1:500 diluted ^13^C-and ^15^N-labeled amino acids standard mix (Sigma-Aldrich, Merck, Darmstadt, Germany) was added to the tubes, followed by vortex and centrifugation. The upper organic phase was discarded. The polar phase was re-extracted with 0.5 mL of MTBE. The lower polar phase was then dried under a gentle stream of nitrogen and kept at-80°C for metabolite analysis. Dry polar samples were re-suspended in 80 µL Methanol:DDW (50:50) and centrifuged twice to remove the debris. The resuspended samples (40 µL) were transferred to HPLC vials for injection.

LC-MS metabolic profiling was done as described by Zheng *et al*. with minor modifications. Briefly, analysis was performed using an Acquity I class UPLC System combined with a Q Exactive Plus Orbitrap™ mass spectrometer (Thermo Fisher Scientific, Waltham, USA), operated in a positive and negative ionization modes. The LC separation was done using a SeQuant Zic-pHilic column (150 mm × 2.1 mm) with a SeQuant guard column (20 mm × 2.1 mm; Merck, Darmstadt, Germany). The mobile phase B contained acetonitrile, and the mobile phase A contained 20 mM ammonium carbonate with 0.1% ammonia hydroxide in water:acetonitrile (80:20, v/v). The flow rate was kept at 200 μL/min and the gradient was as follows: 0-2 min 75% of B, 14 min 25% of B, 18 min 25% of B, 19 min 75% of B, for 4 min, 23 min 75% of B. Data processing was done using the Compound Discoverer software (v3.3; Thermo Fisher Scientific, Waltham, USA), when detected compounds were identified by accurate mass, retention time, isotope pattern, and fragments (data S4). The software reports the fractional label incorporation (exchange rate) after natural abundance correction for each compound. LC-MS data were manually searched for metabolites involved in methyl group metabolism. This included DMSP, 3-*S*-methylmercaptopropionate (3-MMPA), *S*-adenosylmethionine (SAM), *S*-adenosyl 3-(methylsulfanyl)propylamine (dcSAM), *S*-methyl-5’-thioadenosine (MTA), methionine, serine, glycine, 5-methyltetrahydrofolate, 5,10-methylenetetrahydrofolate, 5,10-methenyltetrahydrofolate, *N*^10^-formyltetrahydrofolate, thymidine-5’-phosphate (dTMP), *S*-methyl-5-thio-D-ribofuranose, *S*-methyl-5-thio-α-D-ribose 1-phosphate (1-PMTR), *S*-Methyl-5-thio-D-ribulose 1-phosphate and ATP. The manual search confirmed the presence of labeled DMSP, SAM and MTA (data S4), which were identified by the MS2 fragmentation patterns.

### Bioinformatic analysis of methyl group-related metabolism in algal-associated and plant-associated bacteria

Orthology clusters were employed to assess the presence or absence of methyl group metabolism genes that were identified in *P. inhibenns* (table S3) across the genomes of algal-associated and plant-associated bacteria (Table 1, figures S1 and S17). The analysis was conducted using OrthoFinder^101^, which utilizes MCL^102^ and FastME^103^ algorithms with default settings. Diamond^104^ blast results from an all-against-all search served as input data. Reference genomes were utilized in cases where a specific genome was unavailable. The used genomes and their accession numbers are detailed in table S8.

## Acknowledgments

### General

We appreciate the technical guidance of Dr. Ester Feldmesser, Dr. Bareket Dassa, Dr. Shifra Ben-Dor and Dr. Hadas Keren-Shaul in RNA-sequencing, are thankful for the help of Dr. Ron Rotkopf with statistical analysis, are grateful for the excellent bioinformatic support of Amir Szitenberg, and acknowledge the contribution of Dr. Maxim Itkin with LC-MS analysis (Life Sciences Core Facilities, Weizmann Institute of Science, Israel). We thank Dr. Roi Avraham and Dr. Gili Rosenberg (Weizmann Institute of Science, Israel) for sharing their expertise in dual RNA-sequencing. The compound gonyol was synthesized and kindly provided by Dr. Kathleen Thume and Prof. Georg Pohnert (Friedrich Schiller University, Jena, Germany). We are grateful for inspiring discussions with Dr. Torsten Schubert and Jonathan Hammer (Friedrich Schiller University, Jena, Germany) about cobalamin-dependent one-carbon metabolism, and highly appreciate the stimulating feedback we received from Dr. Elad Noor (Weizmann Institute of Science, Israel), Prof. Uwe Sauer (ETH Zürich, Switzerland), Dr. Anat Bren (Weizmann Institute of Science, Israel) and Dr. Johannes Zimmermann (Christian-Albrechts-University Kiel, Germany) about bacterial metabolism and the lag phase. We are especially thankful for the conceptual guidance and encouragement from the late Prof. Dan Tawfik (Weizmann Institute of Science, Israel) during the early phase of the study.

### Funding

M.S. received a Dean of Faculty Fellowship, a Sir Charles Clore Fellowship (Clore Israel Foundation) and a Senior Postdoc Fellowship. D.A.N.B received the Armando and Maria Jinich Fellowship. The study was funded by the Minerva Foundation with funding from the German Federal Ministry for Education and Research, the Israel Science Foundation (ISF 947/18), the European Research Council (ERC StG 101075514) and the de Botton center for marine sciences, granted to E.S.

### Author contributions

M.S., D.A.N.B. and E.S. designed the study. M.S., D.A.N.B., S.M., V.F. and L.Y. performed and analyzed experiments. J.R. contributed to the design of the research and helped supervise the project. M.S., D.A.N.B. and E.S. wrote the manuscript. All authors discussed the results and contributed to the final manuscript.

### Competing interests

The authors published a patent. Other than that, the authors are not aware of any affiliations, memberships, funding, or financial holdings that might be perceived as affecting the objectivity of this research.

### Data and materials availability

All data are available in the main text or the supplementary materials. Further information and requests for resources and reagents should be directed to and will be fulfilled by the lead contact, Einat Segev. Plasmids generated in this study, as well as bacteria that are not deposited in public strain collections, will be made available on reasonable request. RNA-sequencing data have been deposited at the NCBI Sequence Read Archive (SRA) and are publicly available as of the date of publication. Accession numbers are listed in the Key Resources Table. Raw metabolomics data have been deposited to the EMBL-EBI MetaboLights database with the identifier MTBLS9742 and can be accessed here https://www.ebi.ac.uk/metabolights/MTBLS9742.

## Supplementary Materials

Figs. S1 to S21

Tables S1 to S8

Data S1 to S4

References for Supplementary Materials

