## Supplemental Information for "Reducing the Bacterial Lag Phase Through Methylated Compounds: Insights from Algal-Bacterial Interactions"

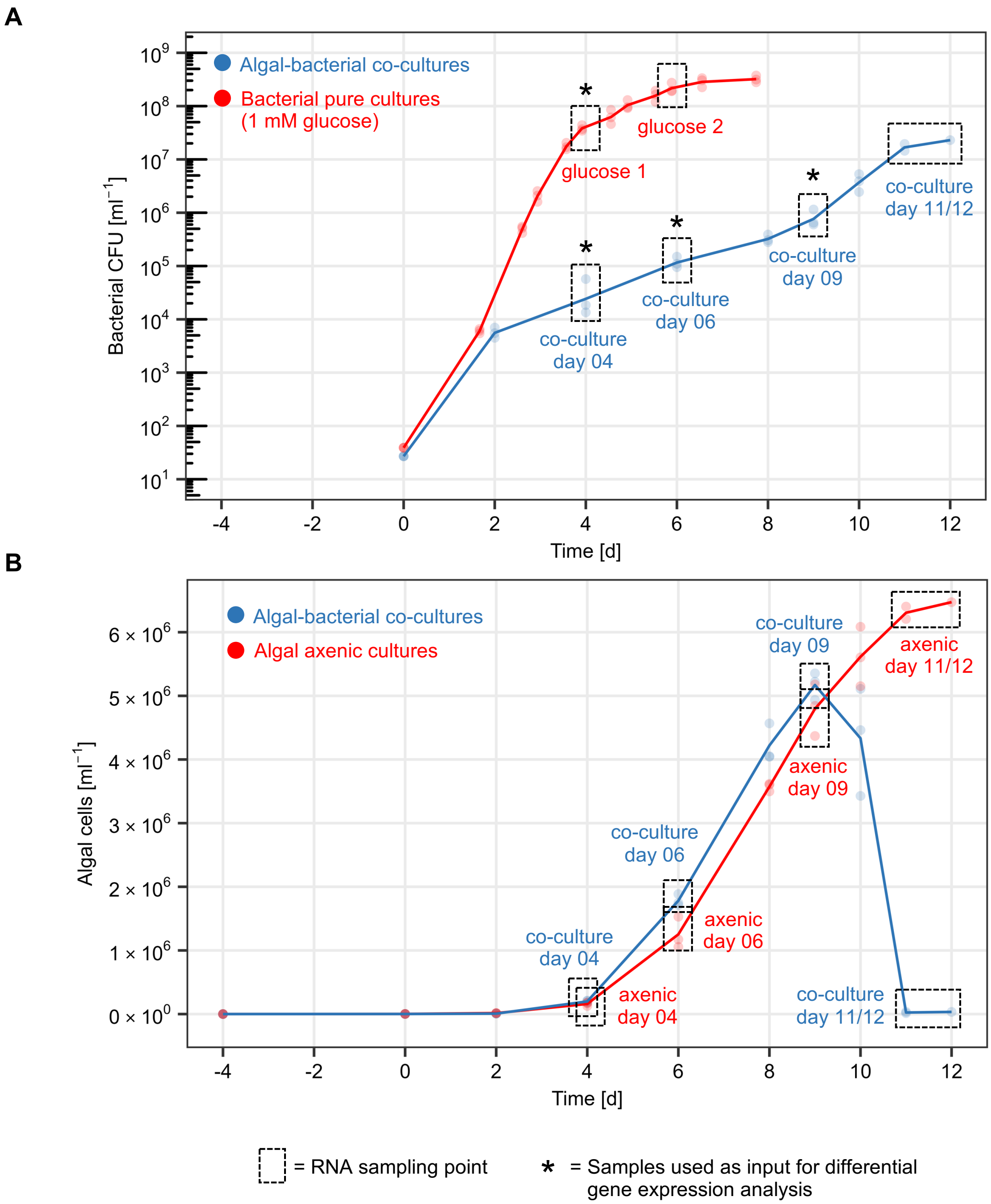


**Fig. S1**: RNA sampling points of microbial cultures. (**A**) RNA was sampled from the late exponential and stationary phases of *P. inhibens* bacteria grown in pure cultures with glucose (red; samples designated glucose 1 and 2, respectively), as well as from the different interaction phases of *P. inhibens* in co-cultures with *E. huxleyi* algae (blue; samples designated co-cultures day 04 to day 11/12). All RNA samples (dashed box) were subjected to co-culture RNA-sequencing (see *Method Details*). Since exponentially growing bacteria in co-cultures are under constant nutrient limitation, they were compared with the late-exponential growth phase in pure cultures, in which nutrients become limiting. As mentioned in the text, the limitations of such a comparison should be acknowledged, yet this analysis can provide valuable observations and serve to generate hypotheses for further exploration. Samples marked with asterisk were included in the differential gene expression analysis to compare exponentially growing bacterial pure cultures (glucose 1) with bacteria in co-cultures with algae (co-culture day 04, 06 and 09). Three flasks were sampled per time point and condition (biological triplicate), and each dot represents the bacterial CFU number (determined as technical duplicate). (**B**) Bacteria stimulate algal growth during the early interaction phase and promote algal death during the late interaction phase (blue: algal growth in co-cultures with bacteria, corresponding to panel A; red: algal growth in axenic cultures, i.e. without bacteria). RNA samples of algae without bacteria (axenic day 04 – 11/12) were not analyzed for differential gene expression in this study. Three flasks were sampled per time point and condition. Each dot represents the mean of algal flow cytometry cell counts, determined as technical triplicate for every sampled flask. The lines in (**A**) and (**B**) depict the mean of the plotted dots for each condition.

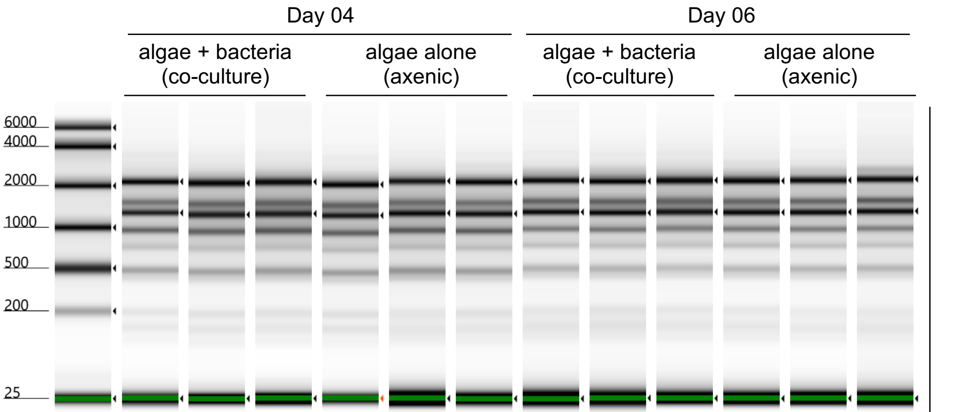


| Sample ID \| replicate | co-culture day 04 \| rep 1 | co-culture day 04 \| rep 2 | co-culture day 04 \| rep 3 | axenic day 04 \| rep 1 | axenic day 04 \| rep 2 | axenic day 04 \| rep 3 | co-culture day 06 \| rep 1 | co-culture day 06 \| rep 2 | co-culture day 06 \| rep 3 | axenic day 06 \| rep 1 | axenic day 06 \| rep 2 | axenic day 06 \| rep 3 |
|---|---|---|---|---|---|---|---|---|---|---|---|---|
| Qubit [ng/ul] | 146.4 | 116 | 105.4 | 155.8 | 68 | 108.8 | 369 | 427 | 441 | 418 | 342 | 342 |
| Tapestation [ng/ul] | 105.4 | 108.4 | 94.6 | 146.8 | 61 | 93.4 | 343 | 388 | 407 | 385 | 308 | 333 |
| Tapestation (RIN) | 6.7 | 6.3 | 6 | 6.4 | 6.4 | 6.6 | 7.2 | 7 | 6.9 | 7.2 | 7.3 | 7.3 |

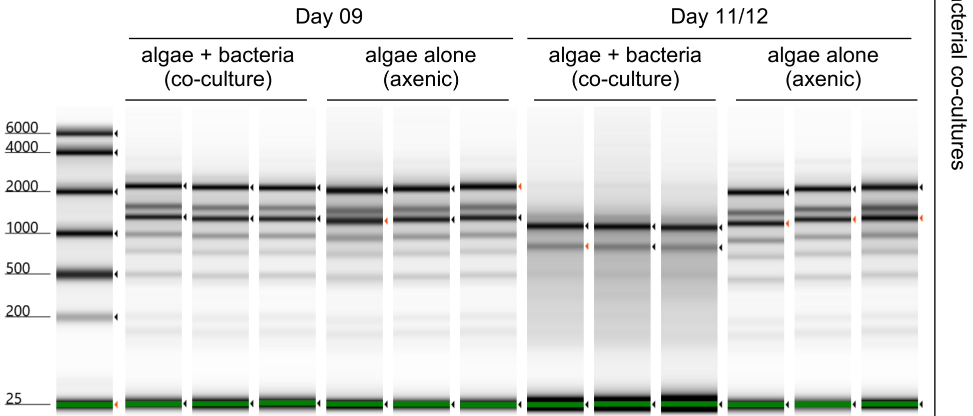


| Sample ID \| replicate | co-culture day 09 \| rep 1 | co-culture day 09 \| rep 2 | co-culture day 09 \| rep 3 | axenic day 09 \| rep 1 | axenic day 09 \| rep 2 | axenic day 09 \| rep 3 | co-culture day 11/12 \| rep 1 | co-culture day 11/12 \| rep 2 | co-culture day 11/12 \| rep 3 | axenic day 11/12 \| rep 1 | axenic day 11/12 \| rep 1 | axenic day 11/12 \| rep 1 |
|---|---|---|---|---|---|---|---|---|---|---|---|---|
| Qubit [ng/ul] | 756 | 622 | 608 | 664 | 688 | 633 | 103.5 | 99.7 | 70.8 | 684 | 775 | 667 |
| Tapestation [ng/ul] | 595 | 460 | 462 | 563 | 600 | 570 | 57.8 | 61.8 | 40.6 | 506 | 592 | 520 |
| Tapestation (RIN) | 7.6 | 7.6 | 7.8 | 6.9 | 7.5 | 7.5 | 4.6 | 4.4 | 4.2 | 7.5 | 7.6 | 7.1 |

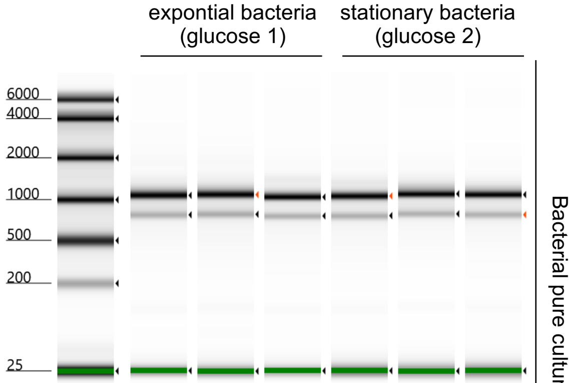


| Sample ID \| replicate | glucose 1 \| rep 1 | glucose 1 \| rep 2 | glucose 1 \| rep 3 | glucose 2 \| rep 1 | glucose 2 \| rep 2 | glucose 2 \| rep 3 |
|---|---|---|---|---|---|---|
| Qubit [ng/ul] | 334 | 274 | 360 | 469 | 511 | 427 |
| Tapestation [ng/ul] | 232 | 214 | 256 | 400 | 427 | 362 |
| Tapestation (RIN) | 8.6 | 8.4 | 8.6 | 8.5 | 8.8 | 8.7 |

**Fig. S2**: Quality control and quantification of RNA samples extracted for co-culture RNA-sequencing analysis. Tapestation intensities were scaled to samples. Arrows indicate ribosomal subunits. RIN—RNA integrity numbers as determined by the Tapestation software.

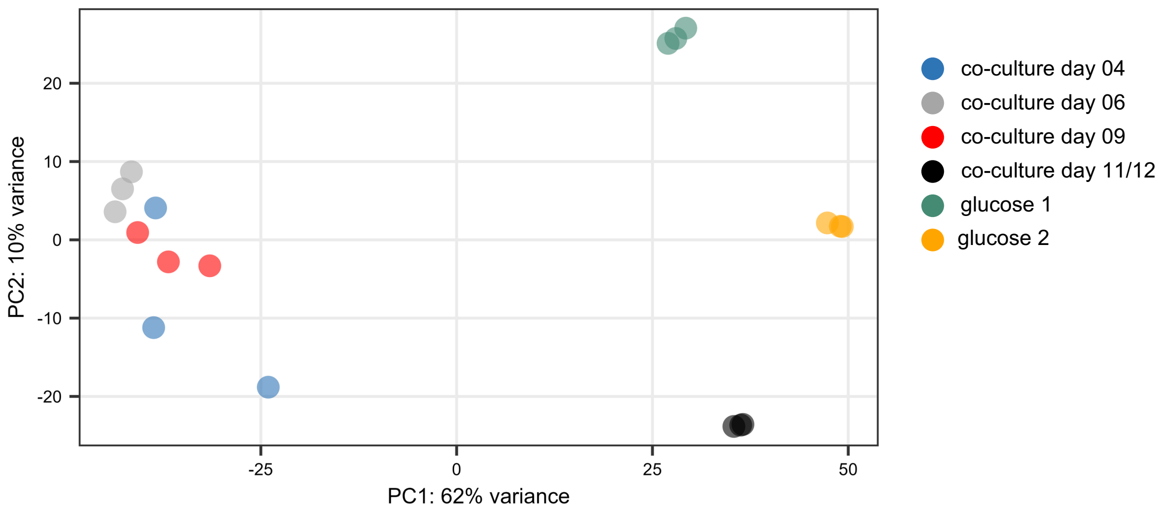


**Fig. S3**: Characterization of bacterial gene transcription profiles in the presence and absence of algae. Bacterial transcription profiles were similar in the presence of live algae (co-culture day 04, 06 and 09), but shifted when algal death was evident (co-culture day 11/12), or in the absence of algae (glucose 1 and 2). Each condition was analyzed as biological triplicate. Based on the close clustering of co-culture samples with live algae (co-culture day 04, 06 and 09), these nine samples were treated as replicates and compared to samples of exponentially growing bacteria on glucose (glucose 1) for differential gene expression analysis. The PCA plot was generated using the DESeq2::plotPCA function with default options (ntop=500; top 500 variable genes) and DESeq2::rlog transformed read counts as input.

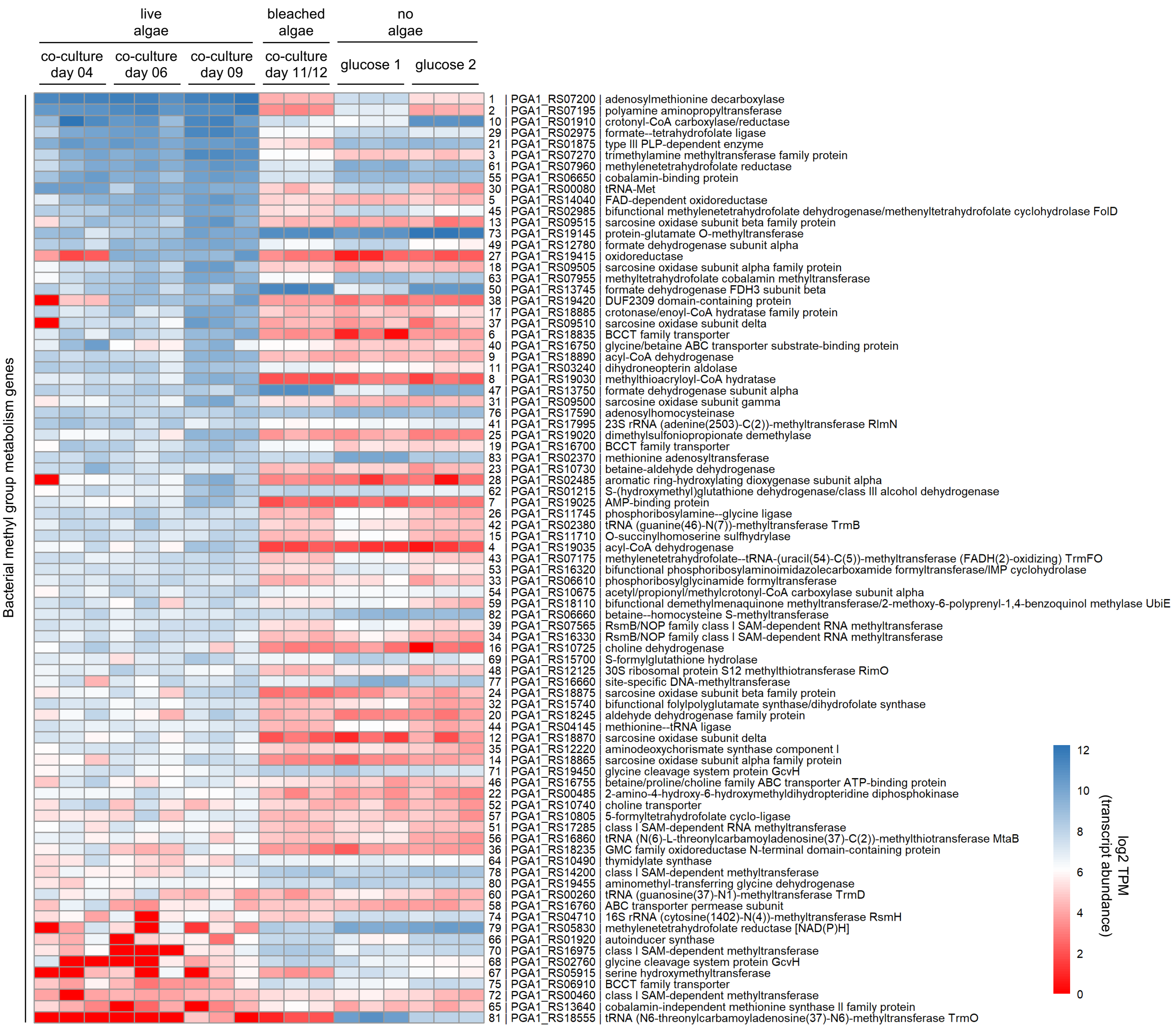


**Fig. S4**: Transcript abundances of 83 manually curated bacterial methyl group metabolism genes in the presence and absence of algae. Transcript abundances were similar in co-cultures with live algae (co-culture day 04, 06 and 09), but shifted when algal death was evident (co-culture day 11/12), or in the absence of algae (glucose 1 and 2). Rows were sorted from highest to lowest mean transcript abundances in co-culture samples with live algae (co-culture day 04, 06 and 09). Transcript abundances are presented as log2 TPM (transcripts per kilobase million) normalized paired-end read counts. Row names include gene numbers (as appears throughout the current study), followed by the RefSeq gene locus tag accession and the RefSeq functional annotation.

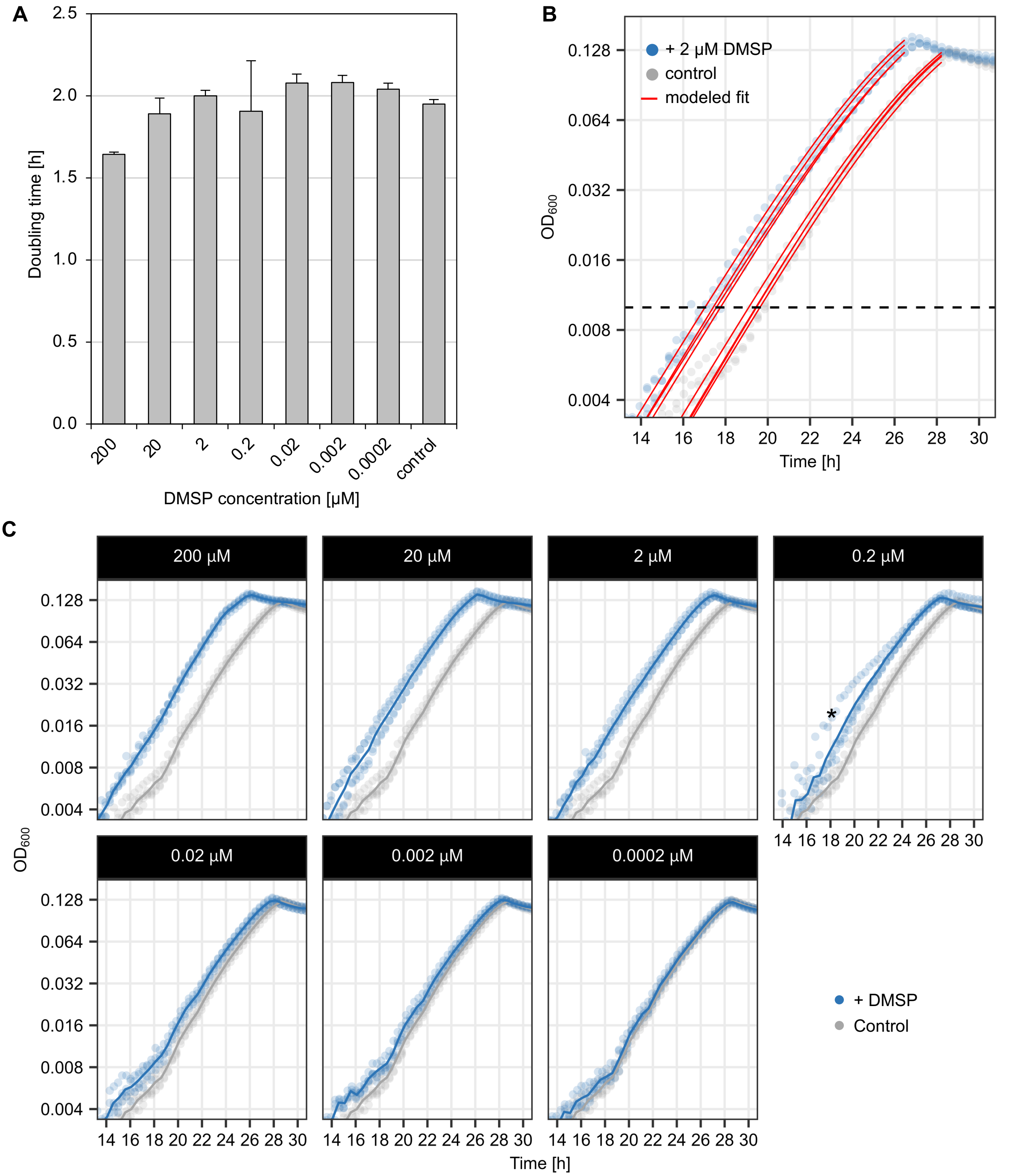


**Fig. S5:** Bacterial growth dynamics in response to DMSP. (**A**) Doubling times of bacterial pure cultures supplemented with different DMSP concentrations, calculated from the modeled fit. Error bars indicate standard deviations for four biological replicates. (**B**) Lag times were determined for individual replicates by fitting an exponential growth function (red lines) and determining the time at which each replicate reached $OD_{600}$ 0.01 (dashed line). (**C**) Growth curves of bacterial cultures supplemented with different DMSP concentrations (indicated above the panels). The $OD_{600}$ measurements were used to calculate lag times (Fig. 2C) and doubling times (panel **B**). The asterisk marks an outlier replicate that was not included in the calculations. Dots represent measurements for individual cultures and lines depict a smoothed average of four biological replicates.

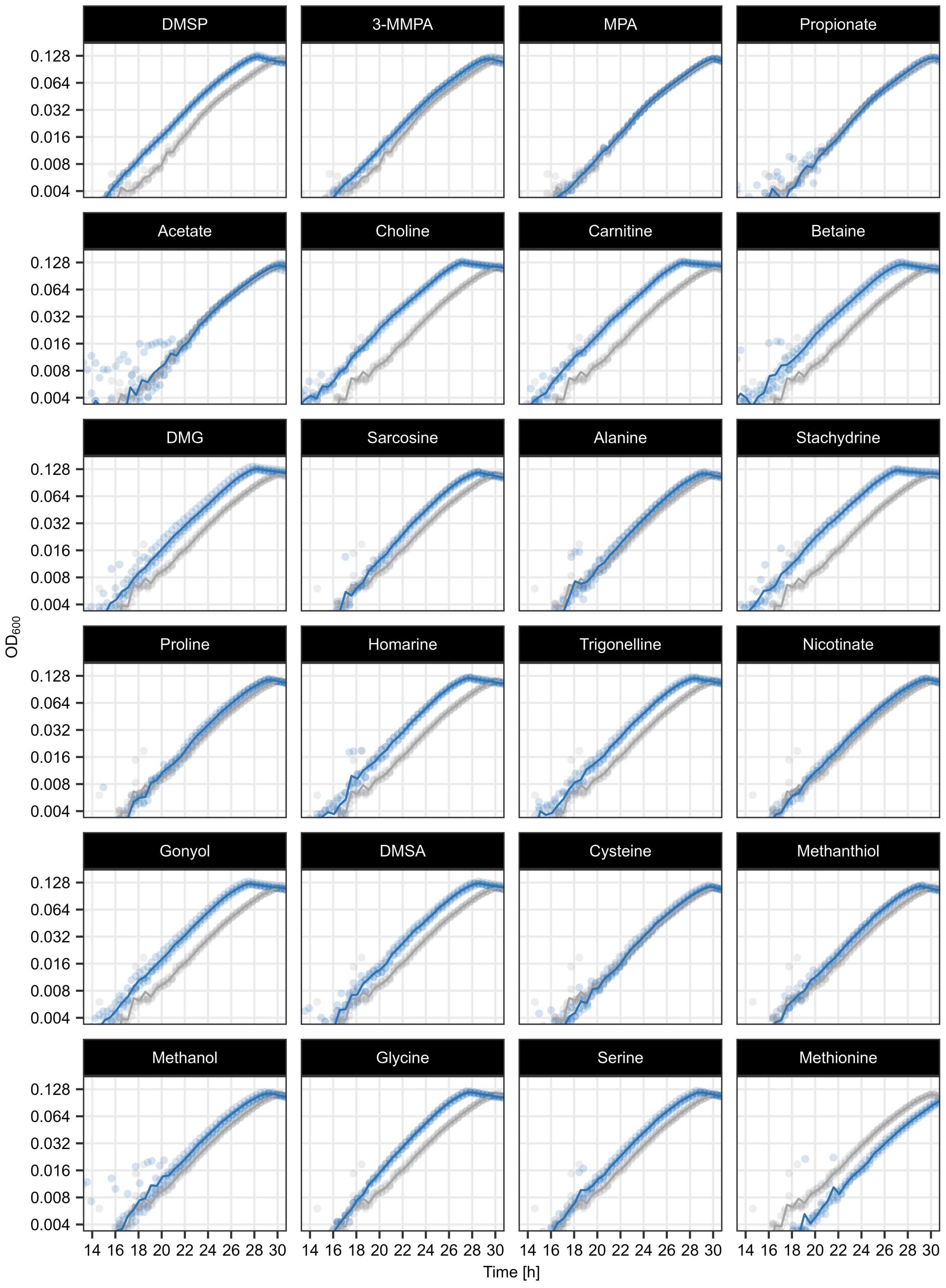
DMSP
3-MMPA
MPA
Propionate
Acetate
Choline
Carnitine
Betaine
DMG
Sarcosine
Alanine
Stachydrine
Proline
Homarine
Trigonelline
Nicotinate
Gonyol
DMSA
Cysteine
Methanthiol
Methanol
Glycine
Serine
Methionine
0.128
0.064
0.032
0.016
0.008
0.004
$OD_{600}$
14 16 18 20 22 24 26 28 30
Time [h]
+ 2 µM supplement
Control

**Fig. S6**

**Fig. S6:** Bacterial growth dynamics in response to different supplements. Growth curves of supplemented cultures (blue) were compared to growth curves of control cultures (grey). Experiments were conducted with four biological replicates. Growth dynamics in response to DMSP analogues (DMSP, 3-MMPA, MPA, propionate, acetate) and other methylated compounds (choline, carnitine, betaine, DMG, sarcosine, alanine, stachydrine, proline, homarine, trigonelline, nicotinate, gonyol, DMSA, cysteine, methanthiol, methanol, glycine, serine, methionine) were measured in artificial seawater medium ($ASW_b$) with 1 mM glucose and 2 μM of the respective supplement. The $OD_{600}$ measurements were used to calculate lag times (Fig. 3A-B) and doubling times (Fig. S7). Dots represent measurements for individual cultures and lines depict a smoothed average of four biological replicates.

**A**

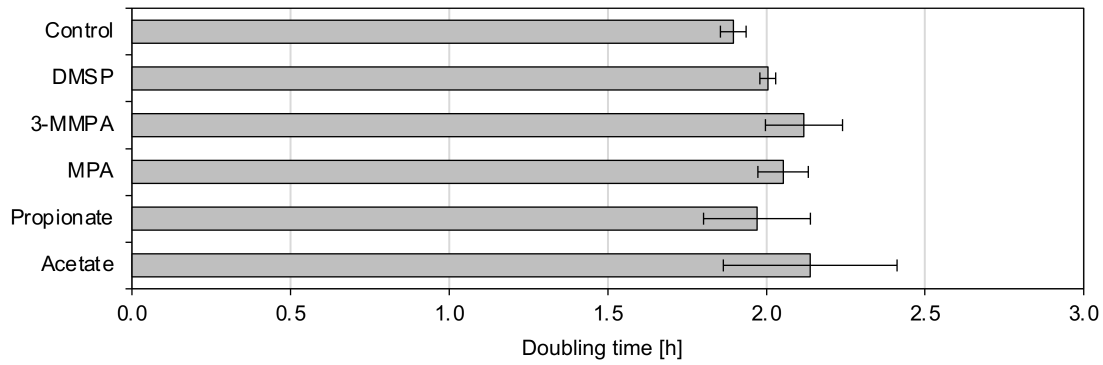


**B**

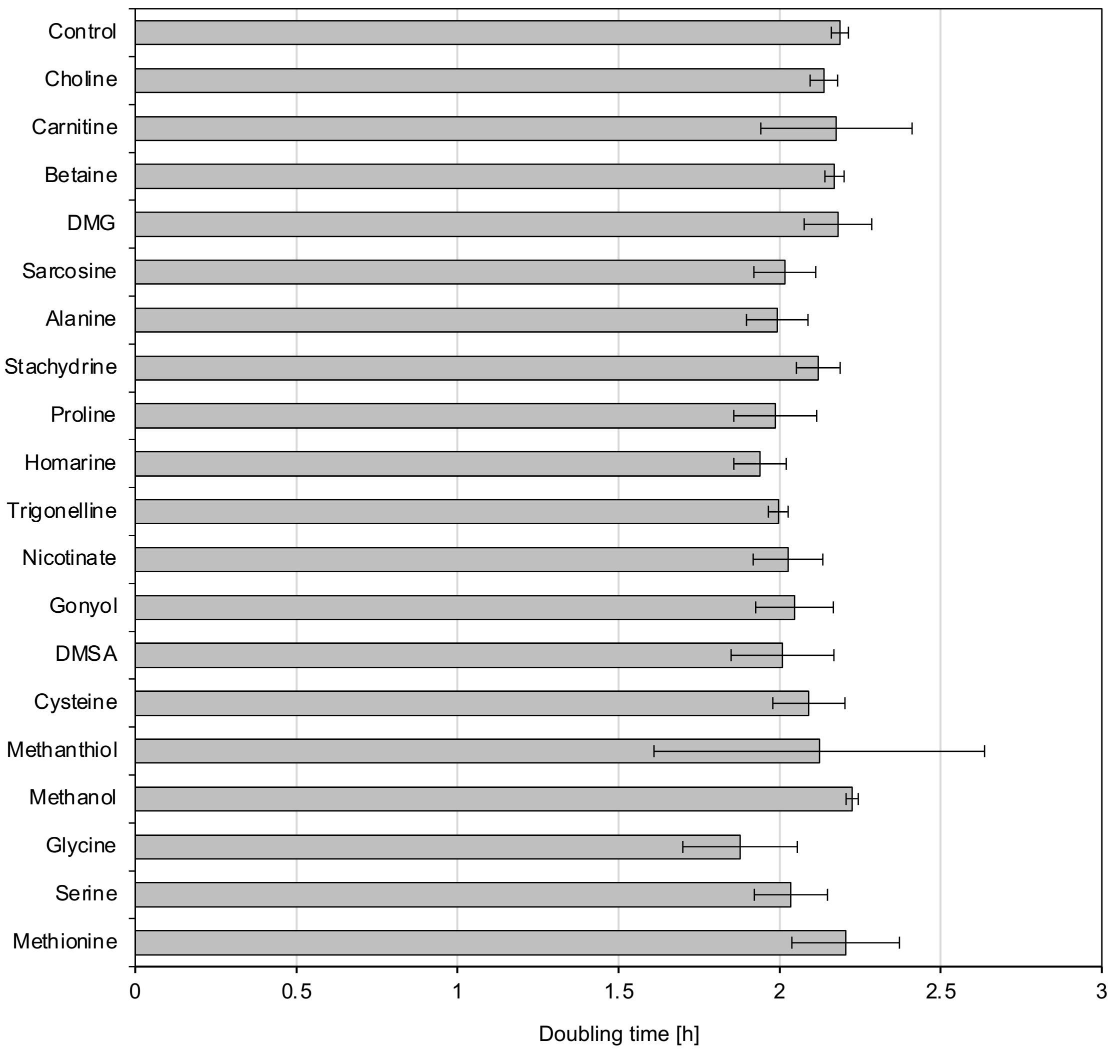


**Fig. S7**: Doubling times of *P. inhibens* bacteria grown in the presence of different supplements. Bacterial growth dynamics were monitored in response to (**A**) analogues of DMSP and (**B**) other supplements, mainly methylated compounds. Cultivations were conducted with four biological replicates in artificial seawater medium ($ASW_b$) with 1 mM glucose and 2 µM of the respective supplement. Error bars indicate standard deviations of the mean. Differences between control cultures and supplemented treatment cultures were not significant using an ANOVA test.

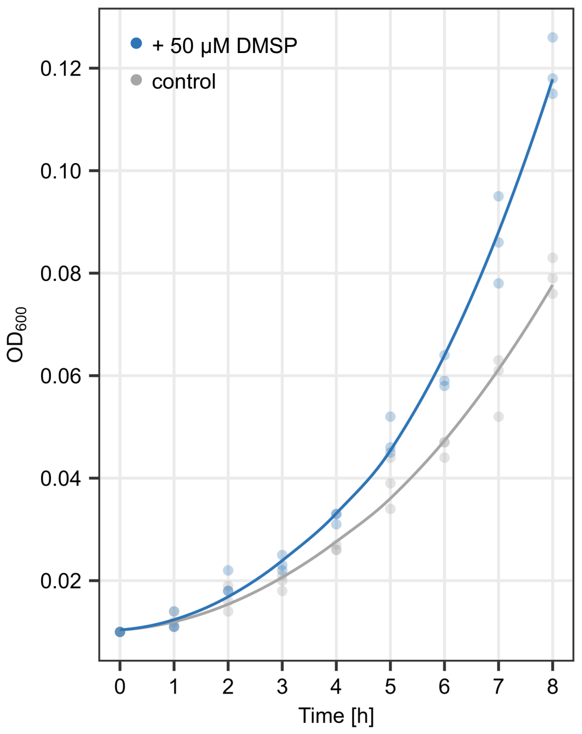


**Fig. S8:** Growth dynamics of *P. inhibens* bacteria in large volume cultures initiated with a denser inoculum and more DMSP. Cultures were inoculated with $OD_{600}$ = OD 0.01 in 60 mL $ASW_b$ medium (1 mM glucose) and supplemented with 50 µM DMSP. A comparatively high starting OD together with more DMSP yielded sufficient biomass for lag phase RNA-sequencing and [$^{13}$C-methyl] DMSP LC-MS analysis. Dots represent measurements for individual cultures and lines depict a smoothed average of three biological replicates.

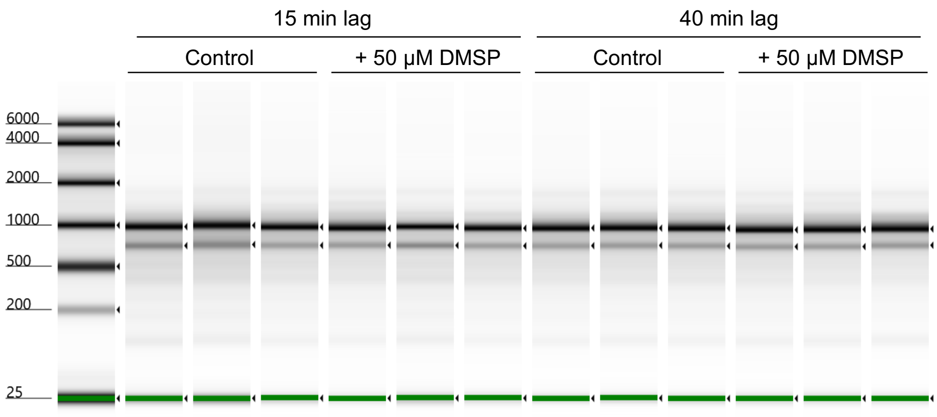


| Sample ID \| replicate | Control (15 min lag) \| rep 1 | Control (15 min lag) \| rep 1 | Control (15 min lag) \| rep 1 | DMSP (15 min lag) \| rep 1 | DMSP (15 min lag) \| rep 2 | DMSP (15 min lag) \| rep 3 | Control (40 min lag) \| rep 1 | Control (40 min lag) \| rep 2 | Control (40 min lag) \| rep 3 | DMSP (40 min lag) \| rep 1 | DMSP (40 min lag) \| rep 2 | DMSP (40 min lag) \| rep 3 |
|---|---|---|---|---|---|---|---|---|---|---|---|---|
| Qubit [ng/ul] | 245 | 195 | 346 | 444 | 240 | 372 | 361 | 337 | 335 | 434 | 460 | 478 |
| Tapestation [ng/ul] | 171 | 155 | 272 | 412 | 199 | 281 | 335 | 297 | 310 | 309 | 303 | 353 |
| Tapestation (RIN) | 6.1 | 5.9 | 6.0 | 6.3 | 6.5 | 6.1 | 6.1 | 6.2 | 6.3 | 6.4 | 6.6 | 6.8 |

**Fig. S9**: Quality control and quantification of RNA samples extracted from *P. inhibens* bacterial pure cultures during the lag phase. Tapestation intensities were scaled to samples. Arrows indicate ribosomal subunits. RIN—RNA integrity numbers as determined by the Tapestation software.

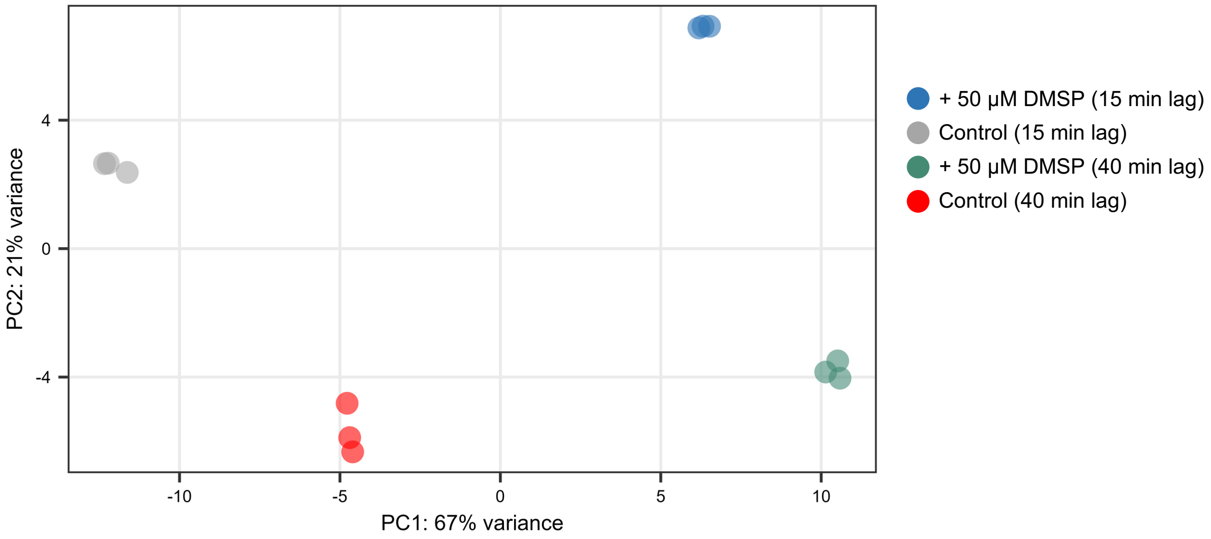


**Fig. S10:** Characterization of bacterial gene transcription profiles during the lag phase in cultures supplemented with DMSP compared to control cultures. Exposure to 50 µM DMSP altered gene transcription profiles during the lag phase, as evident by a pronounced separation of supplemented cultures and control cultures on the PC1 axis. RNA was sampled 15 min and 40 min after transfer of stationary phase bacteria into fresh medium containing 1 mM glucose. Cultures were initiated with a density of $OD_{600}$ = 0.01 (Fig. S8). The PCA plot was generated using the DESeq2::plotPCA function with default options (ntop=500; top 500 variable genes) and DESeq2::rlog transformed read counts as input.

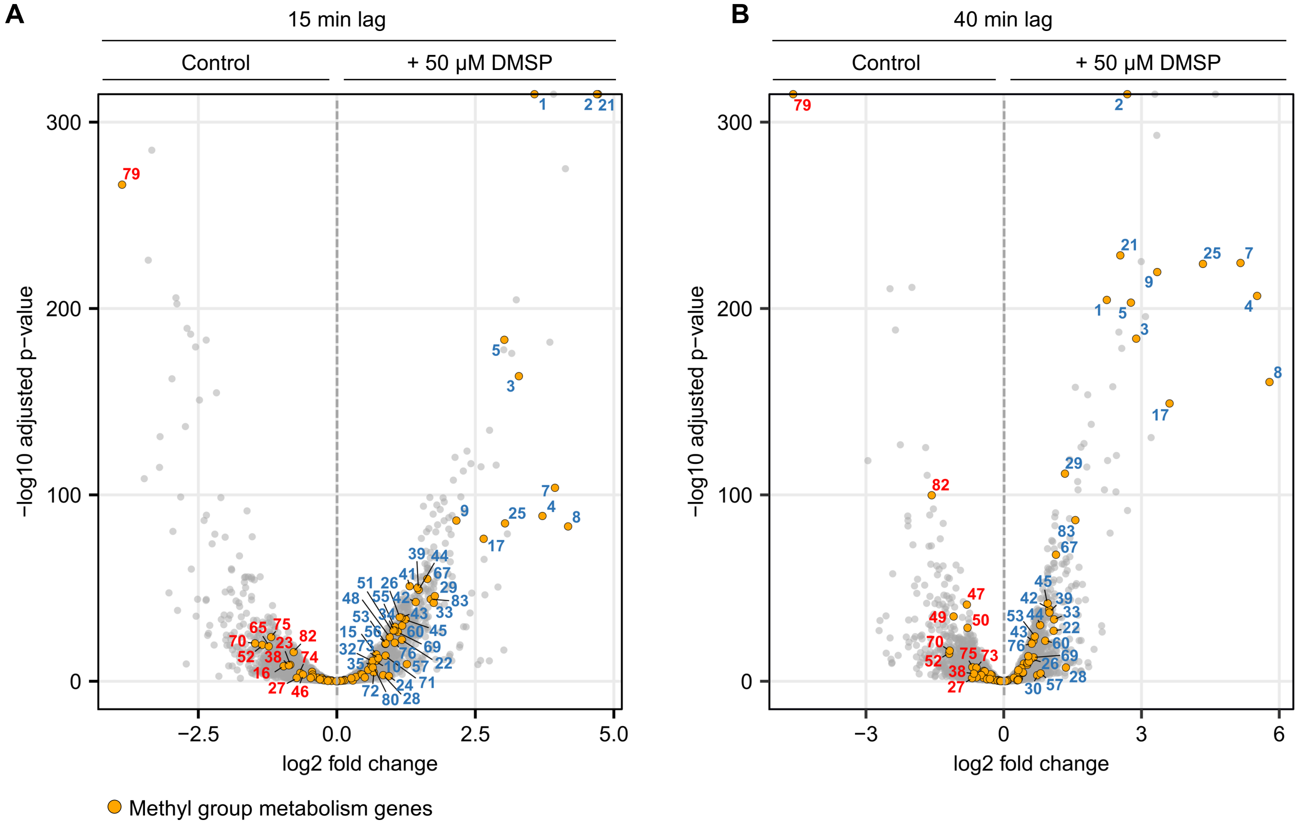


**Fig. S11:** Methyl group metabolism genes were upregulated in DMSP-supplemented lag phase bacteria compared to control cultures. The transcriptional response was measured (**A**) 15 min and (**B**) 40 min after initiation of the cultures. Orange dots depict 83 methyl group metabolism genes of interest (see table S3 for gene accessions and functional annotations). Gene numbers in blue and red indicate genes that were significantly upregulated or downregulated in the presence of DMSP compared to control cultures, respectively (thresholds for coloring: adjusted *p*-value < 0.05; log2 fold change < -0.585 and > +0.585). Overall, 43 and 29 methyl group metabolism genes were upregulated, while 12 and 12 genes were downregulated after 15 min and 40 min, respectively. The most upregulated genes were involved in DMSP catabolism (gene 4, 7, 8, 9, 17, 25) and spermidine synthesis (gene 1, 2 and 21), while the most downregulated gene was associated with the oxidation of methyl groups (gene 79).

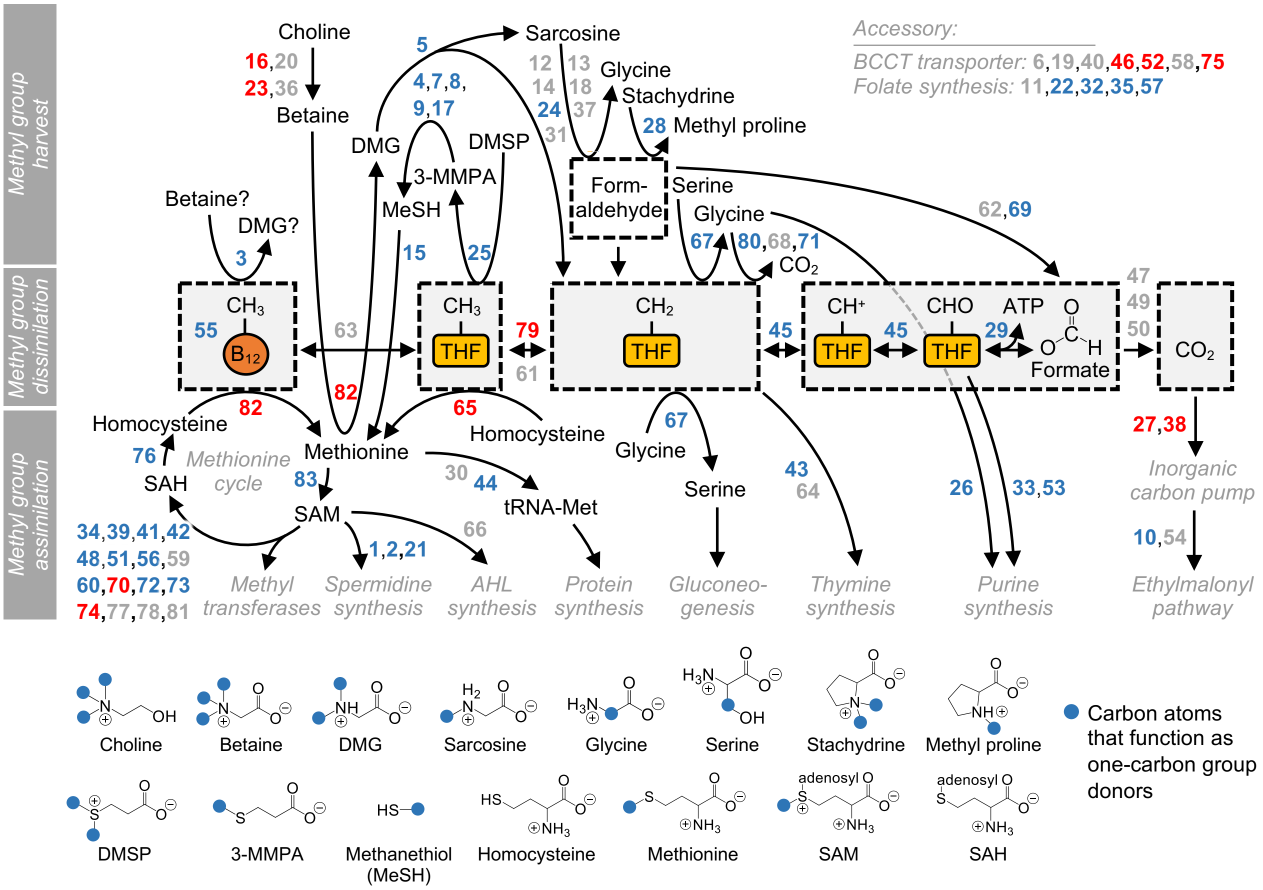


**Fig. S12**: Transcriptional response of *P. inhibens* bacteria towards DMSP during the lag phase. The presented data were generated by harvesting DMSP-supplemented and non-supplemented cultures 15 min after initiation, using biological triplicates (based on data presented in figs. S8-S11, tables S3, S6-S7 and data S2). Numbers in blue, grey and red indicate genes that were upregulated (43 genes), unchanged (28 genes) or downregulated (12 genes) in cultures supplemented with DMSP compared to control cultures, respectively (thresholds for coloring: adjusted *p*-value < 0.05; log2 fold change < -0.585 and > +0.585).

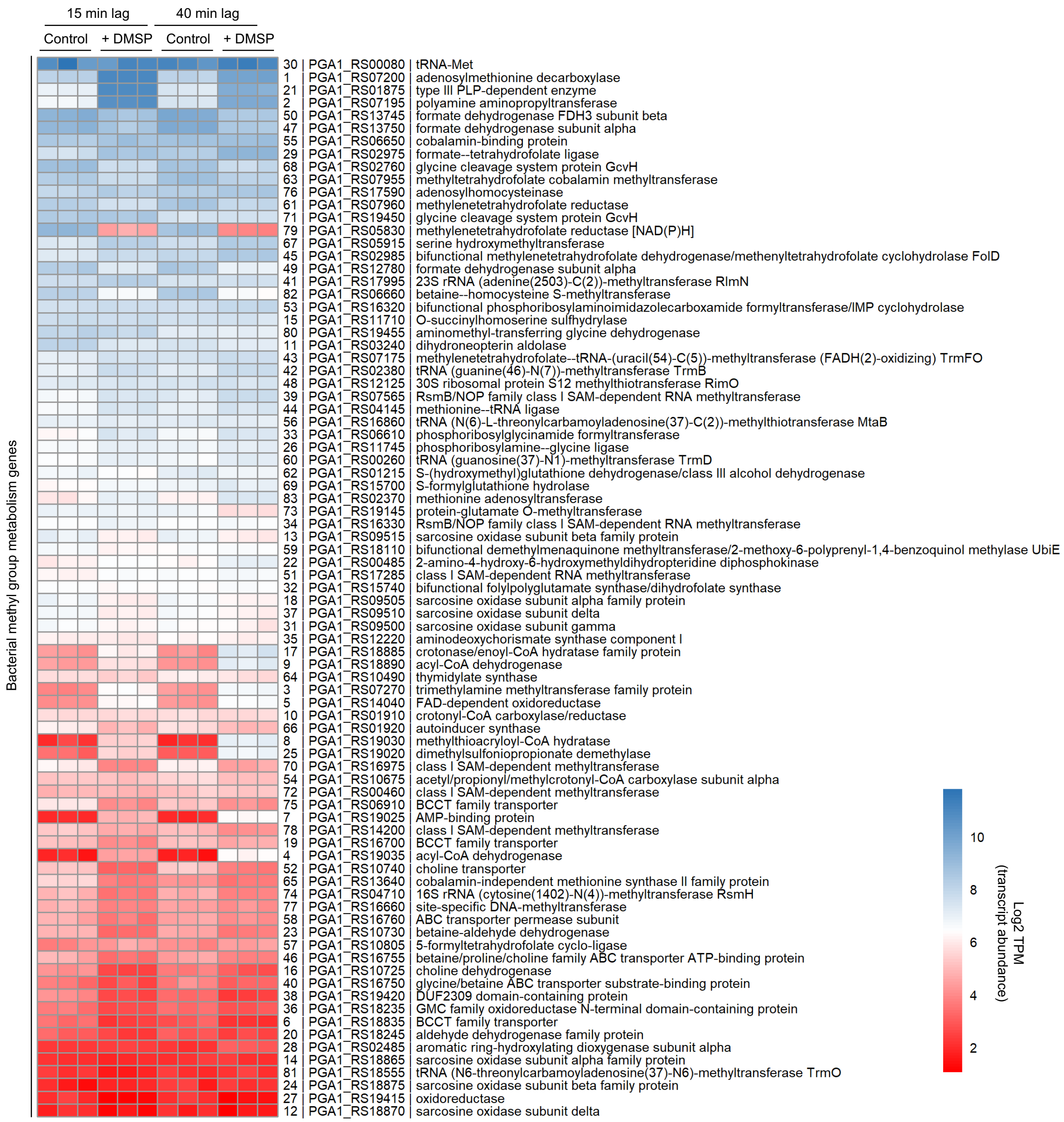


**Fig. S13**: Transcript abundances of methyl group metabolism genes during the lag phase of *P. inhibens* bacteria supplemented with 50 µM DMSP compared to control cultures. Rows were sorted from highest to lowest mean transcript abundances over all samples. Transcript abundances are presented as log2 TPM (transcripts per kilobase million) normalized paired- end read counts. Row names include gene numbers as appear throughout the current study, followed by the RefSeq gene locus tag accession and the RefSeq functional annotations. Based on data presented in figs. S8-S12, tables S3, S6-S7 and data S2.

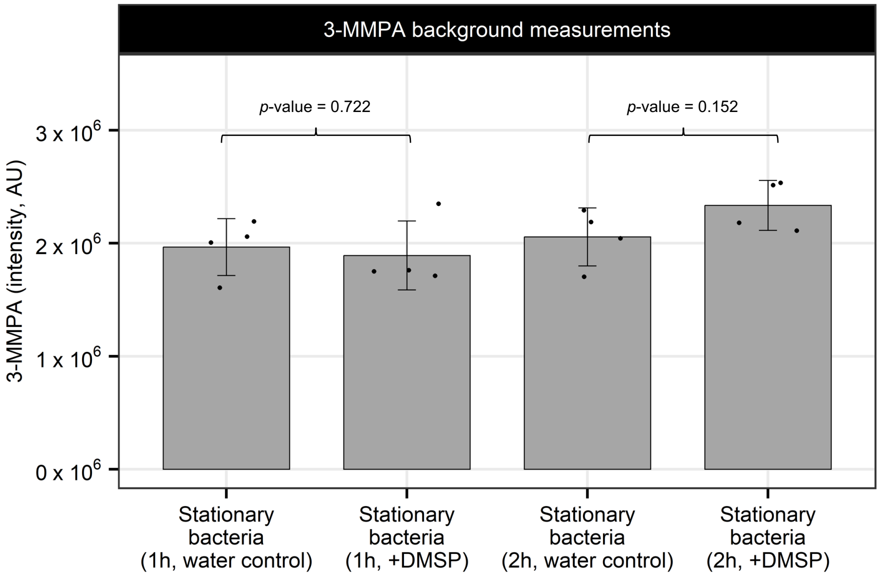


**Fig. S14:** Measurements of 3-MMPA in stationary phase bacteria, prior to generating crude extracts for enzymatic reactions. Samples of stationary bacteria were taken after incubating bacteria for either 1 or 2 hours with 2 µM DMSP (or water as control). For comparison with Fig. 4C, results were normalized to protein concentrations. Measured 3-MMPA levels were similar across all samples. Bars represent the average of four biological replicates with error bars showing the standard deviation. The Student's t-Test was used to calculate *p*-values.

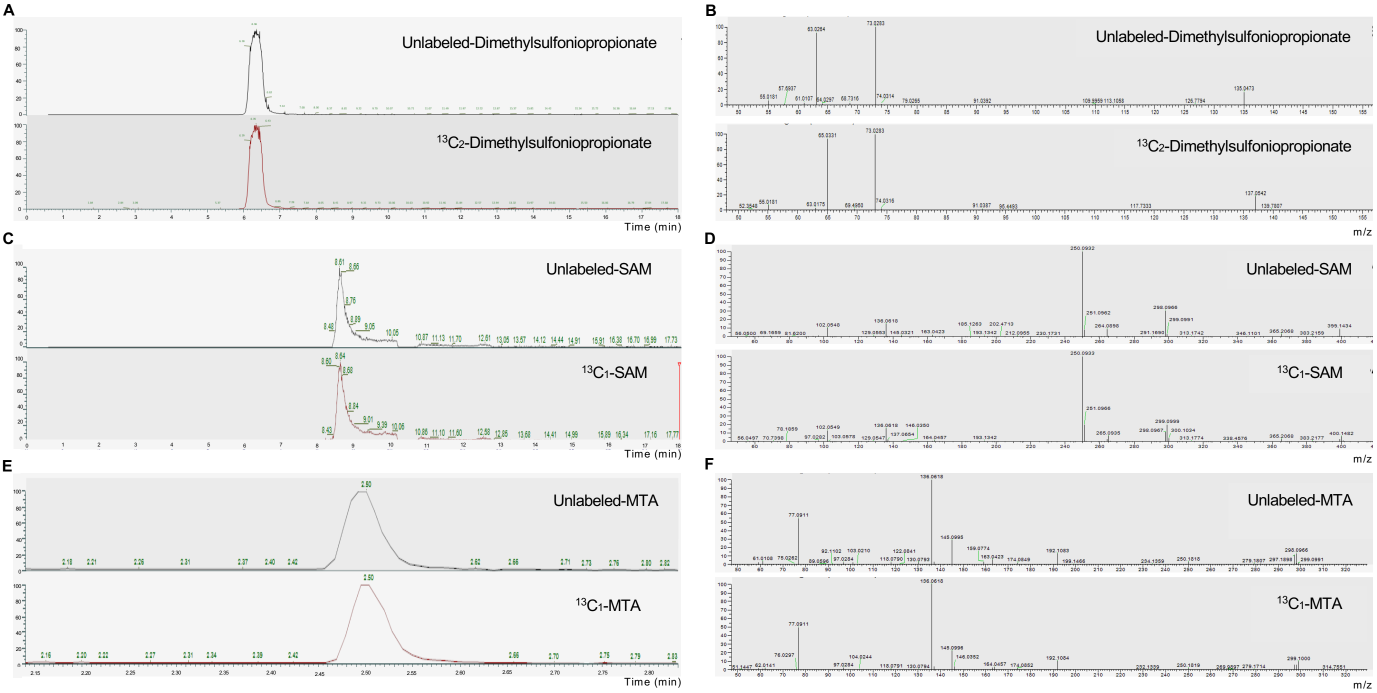


**Fig. S15:** Representative LC-MS/MS chromatograms and MS/MS spectra of detected unlabeled and labeled compounds. (**A-B**) Dimethylsulfoniopropionate (DMSP) and $^{13}C_2$-DMSP. (**C-D**) unlabeled-*S*-Adenosylmethionine (SAM) and $^{13}C_1$-SAM. (**E-F**) unlabeled-5'-*S*-Methyl-5'-thioadenosine (MTA) and $^{13}C_1$-MTA. The observed mass shifts are associated with the $^{13}C$ isotope.

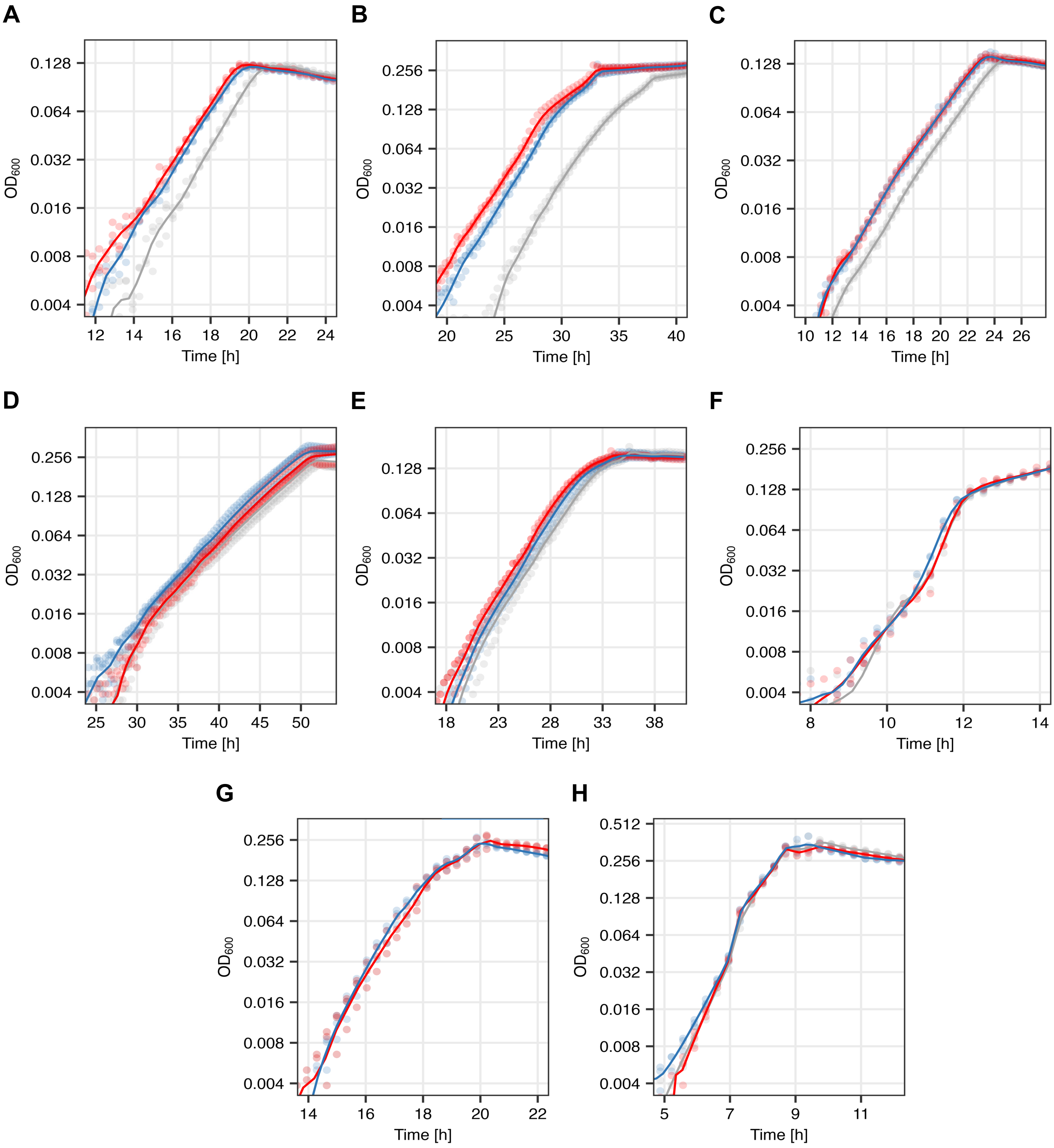


**Fig. S16:** Marine bacteria (**A**) *Sulfitobacter pontiacus* iR4, (**B**) *Labrenzia* sp. DT40, (**C**) *Vibrio* sp. DT22, (**D**) *Phaeobacter* sp. DT104, (**E**) *Pseudoalteromonas* sp. DT35, (**F**) *Marinomonas* sp. DT74, (**G**) *Labrenzia* sp. DT112 and (**H**) *Vibrio* sp. DT18, exhibit various lag reduction capabilities in response to 2 µM DMSP (blue) or betaine (red) when compared to control cultures (grey). Optical Density ($OD_{600}$) measurements are presented in logarithmic scale. Dots represent measurements for individual cultures and lines depict a smoothed average of three replicates.

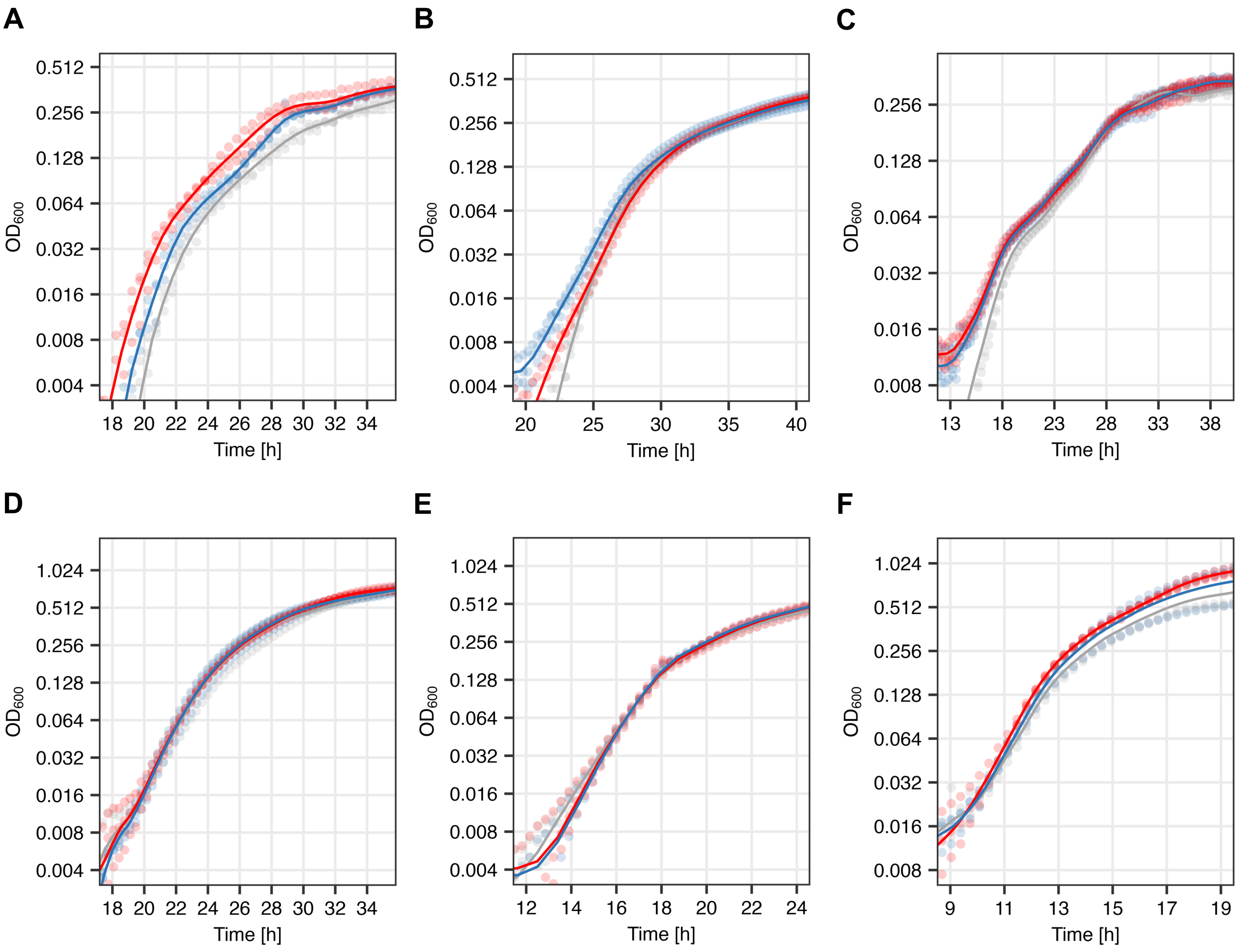


**Fig. S17:** Plant-associated bacteria (**A**) *Bacillus subtilis* 168, (**B**) *Pseudomonas koreensis* PK, (**C**) *Pseudomonas aeruginosa* PAO1, (**D**) *Ewingella americana* EA, (**E**) *Lelliottia amnigena* LA, and (**F**) *Enterobacter aerogenes* Ea, exhibit various lag reduction capabilities in response to 2 μM DMSP (blue) or 2 μM betaine (red) when compared to control cultures (grey). Optical Density ($OD_{600}$) measurements are presented in logarithmic scale. Dots represent measurements for individual cultures and lines depict a smoothed average of three replicates.

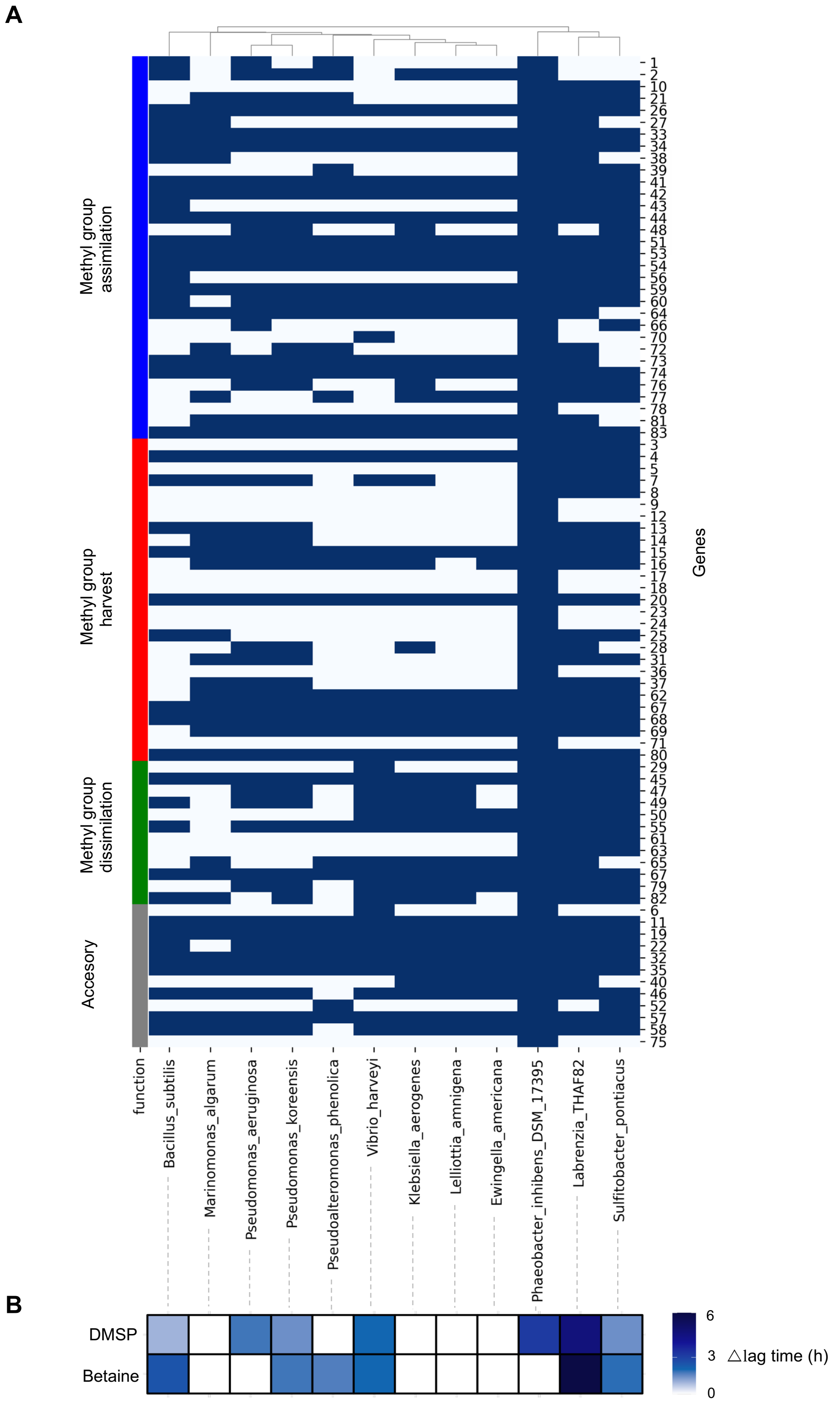


**Fig. S18:** Methyl group-related metabolism in algal-associated and plant-associated bacteria. (**A**) Orthology clusters indicate the presence (blue) or absence (white) of methyl group metabolism genes that were identified in *P. inhibens* (genes numbers correspond to table S3) among the genomes of the tested algal-associated and plant-associated bacteria (Table 1, figs. S16 and S17), as determined by OrthoFinder[1]. OrthoFinder employs MCL[2] and FastME[3] to delineate orthogroups, with default settings retained. Diamond[4] blast results from an all-against-all search were utilized as input. In cases where a genome was unavailable, a reference genome was used (see table S8 for list of genomes with NCBI accession numbers). (**B**) The heatmap shows the diverse lag phase reduction capabilities (in hours) in response to DMSP and betaine among the tested bacteria (Table 1, figs. S16 and S17).

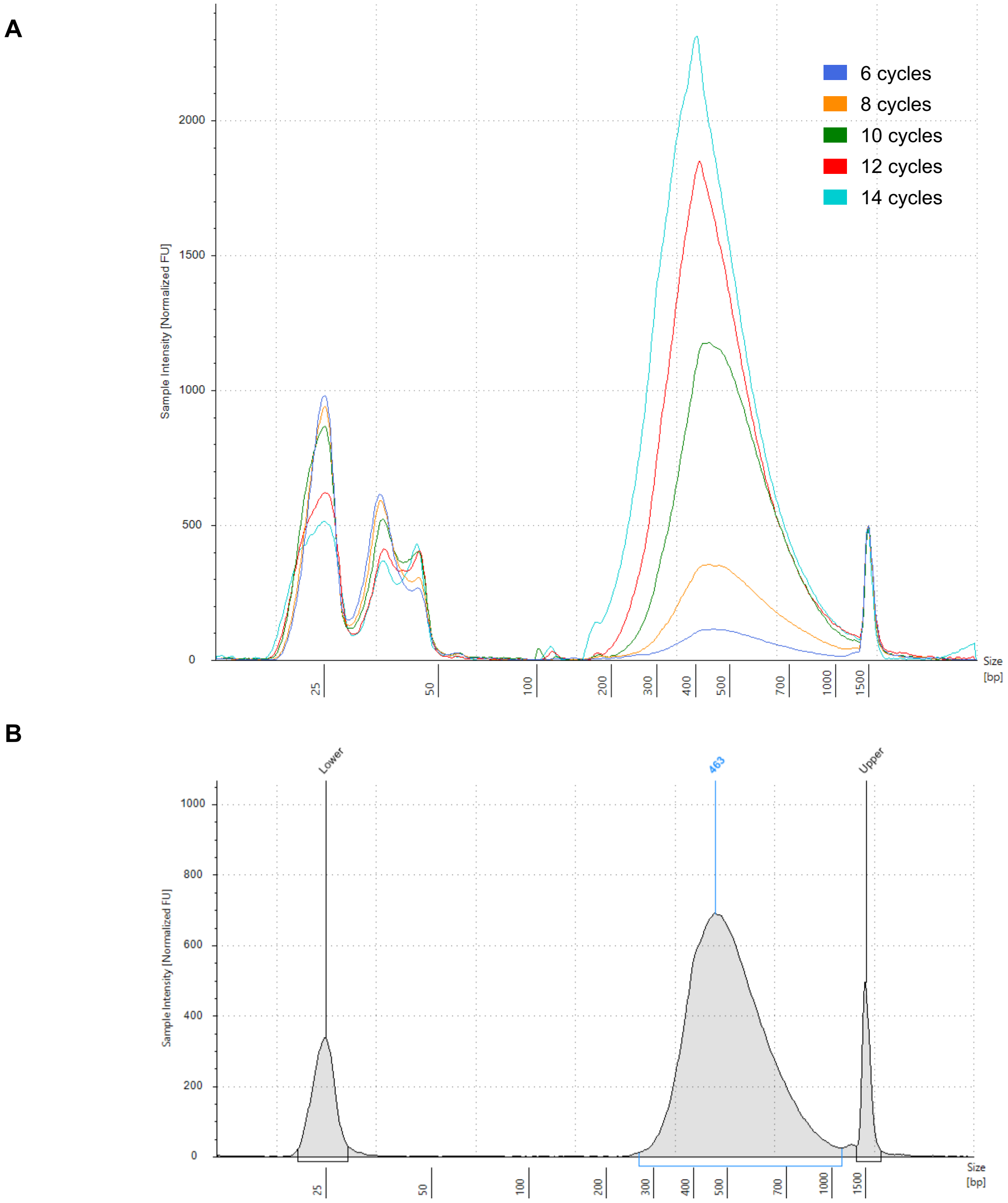


**Fig. S19:** Quality control of calibration RNA-sequencing library. (**A**) Overlay of electropherograms generated for libraries amplified with different PCR cycles. (**B**) Electropherogram of purified RNA-sequencing library. Data were generated with the High Sensitivity D1000 ScreenTape assay (Agilent Technologies, Santa Clara, CA, USA). The purified library exhibited a concentration of 2.97 ng/µl, based on Qubit 1X dsDNA High Sensitivity assay (Thermo Fisher Scientific, Waltham, MA, US

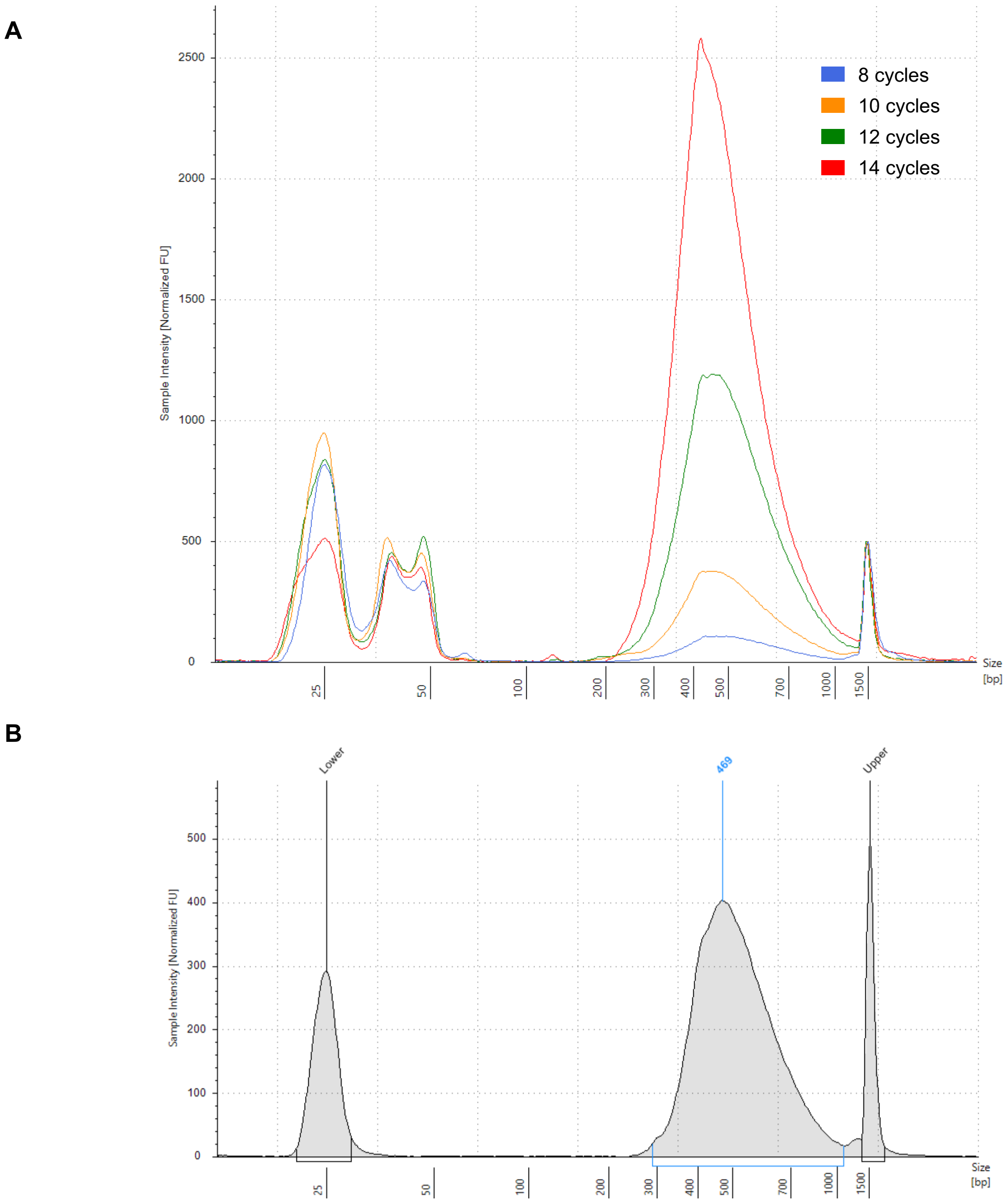


**Fig. S20:** Quality control of co-culture RNA-sequencing library. (**A**) Overlay of electropherograms generated for libraries amplified with different PCR cycles. (**B**) Electropherogram of purified co-culture RNA-sequencing library. Data were generated with the High Sensitivity D1000 ScreenTape assay (Agilent Technologies, Santa Clara, CA, USA). The purified library exhibited a concentration of 1.95 ng/µl, based on Qubit 1X dsDNA High Sensitivity assay (Thermo Fisher Scientific, Waltham, MA, USA).

**A**

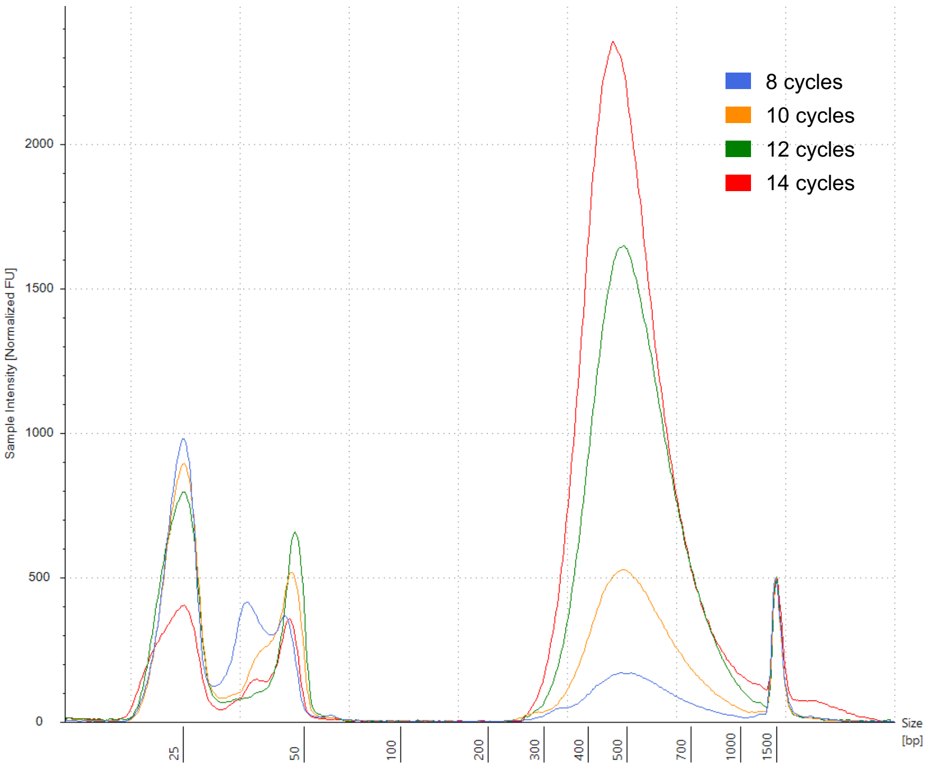


**B**

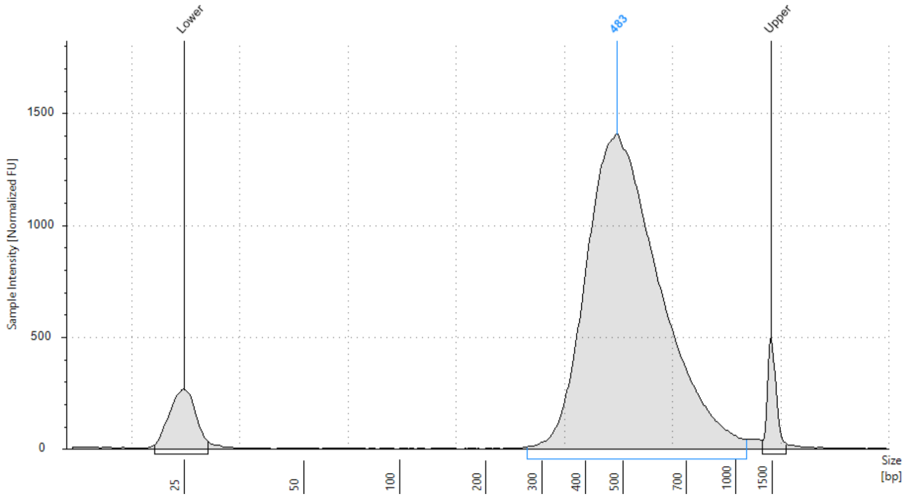


**Fig. S21:** Quality control of lag phase RNA-sequencing library. (**A**) Overlay of electropherograms generated for libraries amplified with different PCR cycles. (**B**) Electropherogram of purified lag phase RNA-sequencing library. Data were generated with the High Sensitivity D1000 ScreenTape assay (Agilent Technologies, Santa Clara, CA, USA). The purified library exhibited a concentration of 6.87 ng/µl, based on Qubit 1X dsDNA High Sensitivity assay (Thermo Fisher Scientific, Waltham, MA, USA).

**Table S1:** Sequencing yields of co-culture RNA-sequencing run.

| | Cycles | Yield (Gbp) | % > Q30 | Reads (M) | Reads passing filter (M) |
|---|---|---|---|---|---|
| Total | 118 | 449.52 | 92.47 | 11,522 | 7,750 |
| Read 1 | 64 | 244.139 | 91.60 | 5,761 | 3,875 |
| Read 2 | 54 | 205.381 | 93.51 | 5,761 | 3,875 |

**Table S2:** Algal and bacterial read counts per sample of co-culture RNA-sequencing run.

| Sample ID \| replicate | Paired-end reads (after demultiplexing) | Paired-end reads (after cutadapt) | *Emiliania huxleyi* feature counts | | *Phaeobacter inhibens* feature counts | |
|---|---|---|---|---|---|---|
| | | | total | non-rRNA | total | non-rRNA |
| **Samples with bacteria** | | | | | | |
| co-culture day 04 \| rep 1 | 394,493,942 | 393,728,523 | 359,778,099 | 356,869,622 | 392,785 | 389,312 |
| co-culture day 04 \| rep 2 | 82,196,551 | 82,043,741 | 75,153,364 | 74,496,050 | 386,857 | 384,464 |
| co-culture day 04 \| rep 3 | 54,291,481 | 54,204,536 | 49,610,317 | 49,154,523 | 378,032 | 375,186 |
| co-culture day 06 \| rep 1 | 345,539,973 | 344,369,218 | 312,349,003 | 310,170,228 | 211,954 | 206,050 |
| co-culture day 06 \| rep 2 | 356,221,349 | 355,409,747 | 322,917,441 | 321,222,154 | 207,351 | 204,656 |
| co-culture day 06 \| rep 3 | 441,433,612 | 440,437,425 | 402,400,748 | 398,384,445 | 227,898 | 221,668 |
| co-culture day 09 \| rep 1 | 470,080,858 | 468,929,331 | 424,468,336 | 420,392,969 | 503,047 | 500,235 |
| co-culture day 09 \| rep 2 | 330,459,894 | 329,572,907 | 297,519,953 | 295,253,525 | 426,485 | 423,914 |
| co-culture day 09 \| rep 3 | 566,156,857 | 564,972,691 | 512,310,003 | 508,633,530 | 430,678 | 427,880 |
| co-culture day 11/12 \| rep 1 | 46,392,253 | 45,102,945 | 22,016,187 | 20,011,612 | 14,968,563 | 14,592,173 |
| co-culture day 11/12 \| rep 2 | 39,800,421 | 38,213,653 | 19,992,702 | 18,072,191 | 10,833,700 | 10,613,582 |
| co-culture day 11/12 \| rep 3 | 43,723,899 | 41,714,542 | 22,274,938 | 20,246,580 | 11,197,490 | 10,953,892 |
| glucose 1 \| rep 1 | 4,023,097 | 4,022,839 | 15,974 | 15,963 | 3,785,341 | 3,531,816 |
| glucose 1 \| rep 2 | 4,161,707 | 4,161,252 | 12,306 | 12,227 | 3,972,265 | 3,794,756 |
| glucose 1 \| rep 3 | 4,355,903 | 4,355,619 | 4,968 | 4,926 | 4,168,592 | 3,946,424 |
| glucose 2 \| rep 1 | 4,464,358 | 4,463,708 | 1,899 | 1,794 | 4,090,721 | 3,753,760 |
| glucose 2 \| rep 2 | 4,571,322 | 4,569,155 | 7,835 | 7,719 | 4,242,487 | 3,723,849 |
| glucose 2 \| rep 3 | 4,471,401 | 4,470,558 | 40,114 | 39,608 | 4,127,911 | 3,408,995 |
| **Samples without bacteria** | | | | | | |
| axenic day 04 \| rep 1 | 35,230,673 | 35,165,646 | 32,384,125 | 32,121,703 | 1,113 | 910 |
| axenic day 04 \| rep 2 | 33,432,598 | 33,352,029 | 30,413,097 | 30,071,739 | 383 | 211 |
| axenic day 04 \| rep 3 | 31,872,791 | 31,808,747 | 29,301,614 | 29,038,128 | 326 | 143 |
| axenic day 06 \| rep 1 | 39,435,756 | 39,334,511 | 36,033,965 | 35,753,053 | 189 | 154 |
| axenic day 06 \| rep 2 | 38,039,758 | 37,919,471 | 34,508,267 | 34,263,463 | 846 | 325 |
| axenic day 06 \| rep 3 | 40,960,442 | 40,882,202 | 37,357,061 | 37,018,630 | 358 | 292 |
| axenic day 09 \| rep 1 | 44,861,993 | 44,756,241 | 40,695,380 | 39,867,427 | 428 | 199 |
| axenic day 09 \| rep 2 | 45,383,172 | 45,280,741 | 41,185,604 | 40,801,350 | 656 | 247 |
| axenic day 09 \| rep 3 | 49,497,859 | 49,349,804 | 44,802,627 | 44,382,768 | 1,789 | 534 |
| axenic day 11/12 \| rep 1 | 41,514,091 | 41,407,679 | 37,368,996 | 37,188,706 | 8,579 | 8,139 |
| axenic day 11/12 \| rep 2 | 45,675,704 | 45,534,841 | 41,097,200 | 40,869,669 | 585 | 459 |
| axenic day 11/12 \| rep 3 | 40,972,617 | 40,868,606 | 36,908,030 | 36,679,879 | 831 | 551 |

* Bacterial reads in algal cultures without bacteria (axenic algal cultures) are due to algal genes with high similarity to bacterial genes. These reads account for roughly 1,000 reads in average per sample, which represent 0.003% of the total reads. Importantly, these reads should not be interpreted as a contamination since potential contaminations were closely monitored (see Materials and Methods).

**Table S3:** Methyl group metabolism genes.

| Gene no. | RefSeq accession | Old accession | RefSeq product | GenBank product | Kegg product |
|---|---|---|---|---|---|
| 1 | PGA1_RS07200 | PGA1_c14480 | adenosylmethionine decarboxylase | S-adenosylmethionine decarboxylase SpeH | speD, AMD1; S-adenosylmethionine decarboxylase [EC:4.1.1.50] |
| 2 | PGA1_RS07195 | PGA1_c14470 | polyamine aminopropyltransferase | spermidine synthase SpeE | speE, SRM, SPE3; spermidine synthase [EC:2.5.1.16] |
| 3 | PGA1_RS07270 | PGA1_c14630 | trimethylamine methyltransferase family protein | trimethylamine methyltransferase MttB | mttB; trimethylamine---corrinoid protein Co-methyltransferase [EC:2.1.1.250] |
| 4 | PGA1_RS19035 | PGA1_262p01860 | acyl-CoA dehydrogenase | putative acyl-Co-A dehydrogenase | dmdC; 3-(methylsulfanyl)propanoyl-CoA dehydrogenase [EC:1.3.99.41] |
| 5 | PGA1_RS14040 | PGA1_c28230 | FAD-dependent oxidoreductase | dimethylglycine dehydrogenase DmgdH | DMGDH; dimethylglycine dehydrogenase [EC:1.5.8.4] |
| 6 | PGA1_RS18835 | PGA1_262p01440 | BCCT family transporter | BCCT transporter | opuD, betL; glycine betaine transporter |
| 7 | PGA1_RS19025 | PGA1_262p01840 | AMP-binding protein | ATP-dependent AMP binding enzyme | dmdB; 3-(methylthio)propionyl---CoA ligase [EC:6.2.1.44] |
| 8 | PGA1_RS19030 | PGA1_262p01850 | methylthioacryloyl-CoA hydratase | putative enoyl-CoA hydratase | dmdD; (methylthio)acryloyl-CoA hydratase [EC:4.2.1.155] |
| 9 | PGA1_RS18890 | PGA1_262p01550 | acyl-CoA dehydrogenase | acyl-CoA dehydrogenase | dmdC; 3-(methylsulfanyl)propanoyl-CoA dehydrogenase [EC:1.3.99.41] |
| 10 | PGA1_RS01910 | PGA1_c03870 | crotonyl-CoA carboxylase/reductase | crotonyl-CoA reductase | ccr; crotonyl-CoA carboxylase/reductase [EC:1.3.1.85] |
| 11 | PGA1_RS03240 | PGA1_c06530 | dihydroneopterin aldolase | dihydroneopterin aldolase-like protein | folB; 7,8-dihydroneopterin aldolase/epimerase/oxygenase [EC:4.1.2.25 5.1.99.8 1.13.11.81] |
| 12 | PGA1_RS18870 | PGA1_262p01510 | sarcosine oxidase subunit delta | sarcosine oxidase, delta subunit | mgdB; methylglutamate dehydrogenase subunit B [EC:1.5.99.5] |
| 13 | PGA1_RS09515 | PGA1_c19170 | sarcosine oxidase subunit beta family protein | sarcosine oxidase subunit beta | soxB; sarcosine oxidase, subunit beta [EC:1.5.3.24 1.5.3.1] |
| 14 | PGA1_RS18865 | PGA1_262p01500 | sarcosine oxidase subunit alpha family protein | sarcosine oxidase subunit alpha | mgdC; methylglutamate dehydrogenase subunit C [EC:1.5.99.5] |
| 15 | PGA1_RS11710 | PGA1_c23540 | O-succinylhomoserine sulfhydrylase | O-succinylhomoserine sulfhydrylase MetZ | metZ; O-succinylhomoserine sulfhydrylase [EC:2.5.1.-] |
| 16 | PGA1_RS10725 | PGA1_c21660 | choline dehydrogenase | choline dehydrogenase BetA | betA, CHDH; choline dehydrogenase [EC:1.1.99.1] |
| 17 | PGA1_RS18885 | PGA1_262p01540 | crotonase/enoyl-CoA hydratase family protein | putative enoyl-CoA hydratase / isomerase | dmdD; (methylthio)acryloyl-CoA hydratase [EC:4.2.1.155] |
| 18 | PGA1_RS09505 | PGA1_c19150 | sarcosine oxidase subunit alpha family protein | sarcosine oxidase subunit alpha | soxA; sarcosine oxidase, subunit alpha [EC:1.5.3.24 1.5.3.1] |
| 19 | PGA1_RS16700 | PGA1_c33650 | BCCT family transporter | putative transporter, BCCT family | |
| 20 | PGA1_RS18245 | PGA1_262p00230 | aldehyde dehydrogenase family protein | betaine aldehyde dehydrogenase BetB | ALDH; aldehyde dehydrogenase (NAD+) [EC:1.2.1.3] |
| 21 | PGA1_RS01875 | PGA1_c03800 | type III PLP-dependent enzyme | putative lysine/ornithine decarboxylase | E4.1.1.17, ODC1, speC, speF; ornithine decarboxylase [EC:4.1.1.17] |
| 22 | PGA1_RS00485 | PGA1_c00990 | 2-amino-4-hydroxy-6-hydroxymethyldihydropteridine diphosphokinase | 2-amino-4-hydroxy-6-hydroxymethyldihydropteridine pyrophosphokinase FolK | folK; 2-amino-4-hydroxy-6-hydroxymethyldihydropteridine diphosphokinase [EC:2.7.6.3] |
| 23 | PGA1_RS10730 | PGA1_c21670 | betaine-aldehyde dehydrogenase | betaine aldehyde dehydrogenase BetB | betB, gbsA; betaine-aldehyde dehydrogenase [EC:1.2.1.8] |
| 24 | PGA1_RS18875 | PGA1_262p01520 | sarcosine oxidase subunit beta family protein | sarcosine oxidase 3 subunit beta | mgdA; methylglutamate dehydrogenase subunit A [EC:1.5.99.5] |
| 25 | PGA1_RS19020 | PGA1_262p01830 | dimethylsulfoniopropionate demethylase | putative aminomethyltransferase | dmdA; dimethylsulfoniopropionate demethylase [EC:2.1.1.269] |
| 26 | PGA1_RS11745 | PGA1_c23610 | phosphoribosylamine--glycine ligase | phosphoribosylamine--glycine ligase PurD | purD; phosphoribosylamine---glycine ligase [EC:6.3.4.13] |
| 27 | PGA1_RS19415 | PGA1_78p00230 | oxidoreductase | putative NADH-ubiquinone oxidoreductase | ndhF; NAD(P)H-quinone oxidoreductase subunit 5 [EC:7.1.1.2] |
| 28 | PGA1_RS02485 | PGA1_c05010 | aromatic ring-hydroxylating dioxygenase subunit alpha | putative dioxygenase | stc2, hpbB; stachydrine N-demethylase [EC:1.14.13.247] |
| 29 | PGA1_RS02975 | PGA1_c05990 | formate--tetrahydrofolate ligase | formate--tetrahydrofolate ligase Fhs | fhs; formate--tetrahydrofolate ligase [EC:6.3.4.3] |
| 30 | PGA1_RS00080 | PGA1_c00160 | tRNA-Met | tRNA-Met | |
| 31 | PGA1_RS09500 | PGA1_c19140 | sarcosine oxidase subunit gamma | sarcosine oxidase-like protein | soxG; sarcosine oxidase, subunit gamma [EC:1.5.3.24 1.5.3.1] |
| 32 | PGA1_RS15740 | PGA1_c31680 | bifunctional folylpolyglutamate synthase/dihydrofolate synthase | putative bifunctional enzyme FolC | folC; dihydrofolate synthase / folylpolyglutamate synthase [EC:6.3.2.12 6.3.2.17] |
| 33 | PGA1_RS06610 | PGA1_c13270 | phosphoribosylglycinamide formyltransferase | phosphoribosylglycinamide formyltransferase PurN | purN; phosphoribosylglycinamide formyltransferase 1 [EC:2.1.2.2] |
| 34 | PGA1_RS16330 | PGA1_c32880 | RsmB/NOP family class I SAM-dependent RNA methyltransferase | putative ribosomal RNA small subunit methyltransferase B | rsmB, sun; 16S rRNA (cytosine967-C5)-methyltransferase [EC:2.1.1.176] |
| 35 | PGA1_RS12220 | PGA1_c24550 | aminodeoxychorismate synthase component I | putative para-aminobenzoate synthase component 1 | pabB; para-aminobenzoate synthetase component I [EC:2.6.1.85] |
| 36 | PGA1_RS18235 | PGA1_262p00210 | GMC family oxidoreductase N-terminal domain-containing protein | alcohol dehydrogenase AlkJ | betA, CHDH; choline dehydrogenase [EC:1.1.99.1] |
| 37 | PGA1_RS09510 | PGA1_c19160 | sarcosine oxidase subunit delta | sarcosine oxidase 2 subunit delta | soxD; sarcosine oxidase, subunit delta [EC:1.5.3.24 1.5.3.1] |
| 38 | PGA1_RS19420 | PGA1_78p00240 | DUF2309 domain-containing protein | hypothetical protein | K09822; uncharacterized protein |
| 39 | PGA1_RS07565 | PGA1_c15230 | RsmB/NOP family class I SAM-dependent RNA methyltransferase | rRNA methyltransferase-like protein | rsmB, sun; 16S rRNA (cytosine967-C5)-methyltransferase [EC:2.1.1.176] |
| 40 | PGA1_RS16750 | PGA1_c33750 | glycine/betaine ABC transporter substrate-binding protein | putative glycine betaine transport system substrate binding protein | proX; glycine betaine/proline transport system substrate-binding protein |
| 41 | PGA1_RS17995 | PGA1_c36240 | 23S rRNA (adenine(2503)-C(2))-methyltransferase RlmN | ribosomal RNA large subunit methyltransferase N | rlmN; 23S rRNA (adenine2503-C2)-methyltransferase [EC:2.1.1.192] |
| 42 | PGA1_RS02380 | PGA1_c04800 | tRNA (guanine(46)-N(7))-methyltransferase TrmB | tRNA (guanine-N(7)-)-methyltransferase TrmB | trmB, METTL1, TRM8; tRNA (guanine-N7-)-methyltransferase [EC:2.1.1.33] |
| 43 | PGA1_RS07175 | PGA1_c14430 | methylenetetrahydrofolate--tRNA-(uracil(54)-C(5))-methyltransferase (FADH(2)-oxidizing) TrmFO | methylenetetrahydrofolate-tRNA-(uracil-5-)-methyltransferase | trmFO, gid; methylenetetrahydrofolate--tRNA-(uracil-5-)-methyltransferase [EC:2.1.1.74] |
| 44 | PGA1_RS04145 | PGA1_c08340 | methionine--tRNA ligase | methionyl-tRNA synthetase MetG | MARS, metG; methionyl-tRNA synthetase [EC:6.1.1.10] |
| 45 | PGA1_RS02985 | PGA1_c06010 | bifunctional methylenetetrahydrofolate dehydrogenase/methenyltetrahydrofolate cyclohydrolase FolD | bifunctional protein FolD | folD; methylenetetrahydrofolate dehydrogenase (NADP+) / methenyltetrahydrofolate cyclohydrolase [EC:1.5.1.5 3.5.4.9] |
| 46 | PGA1_RS16755 | PGA1_c33760 | betaine/proline/choline family ABC transporter ATP-binding protein | putative glycine betaine transport ATP-binding protein | proV; glycine betaine/proline transport system ATP-binding protein [EC:7.6.2.9] |
| 47 | PGA1_RS13750 | PGA1_c27650 | formate dehydrogenase subunit alpha | putative formate dehydrogenase H | fdoG, fdhF, fdwA; formate dehydrogenase major subunit [EC:1.17.1.9] |

**Table S3** (continued)

| Gene no. | RefSeq accession | Old accession | RefSeq product | GenBank product | Kegg product |
|---|---|---|---|---|---|
| 48 | PGA1_RS12125 | PGA1_c24370 | 30S ribosomal protein S12 methylthiotransferase RimO | ribosomal protein S12 methylthiotransferase RimO | rimO; ribosomal protein S12 methylthiotransferase [EC:2.8.4.4] |
| 49 | PGA1_RS12780 | PGA1_c25740 | formate dehydrogenase subunit alpha | formate dehydrogenase H | fdoG, fdhF, fdwA; formate dehydrogenase major subunit [EC:1.17.1.9] |
| 50 | PGA1_RS13745 | PGA1_c27640 | formate dehydrogenase FDH3 subunit beta | formate dehydrogenase iron-sulfur subunit | fdoH, fdsB; formate dehydrogenase iron-sulfur subunit |
| 51 | PGA1_RS17285 | PGA1_c34810 | class I SAM-dependent RNA methyltransferase | putative RNA methyltransferase | rumA; 23S rRNA (uracil1939-C5)-methyltransferase [EC:2.1.1.190] |
| 52 | PGA1_RS10740 | PGA1_c21690 | choline transporter | transporter, BCCT family | opuD, betL; glycine betaine transporter |
| 53 | PGA1_RS16320 | PGA1_c32860 | bifunctional phosphoribosylaminoimidazolecarboxamide formyltransferase/IMP cyclohydrolase | bifunctional purine biosynthesis protein PurH | purH; phosphoribosylaminoimidazolecarboxamide formyltransferase / IMP cyclohydrolase [EC:2.1.2.3 3.5.4.10] |
| 54 | PGA1_RS10675 | PGA1_c21540 | acetyl/propionyl/methylcrotonyl-CoA carboxylase subunit alpha | propionyl-CoA carboxylase alpha chain | PCCA, pccA; propionyl-CoA carboxylase alpha chain [EC:6.4.1.3] |
| 55 | PGA1_RS06650 | PGA1_c13350 | cobalamin-binding protein | putative dimethylamine corrinoid protein | |
| 56 | PGA1_RS16860 | PGA1_c33960 | tRNA (N(6)-L-threonylcarbamoyladenosine(37)-C(2))-methylthiotransferase MtaB | MiaB-like tRNA modifying enzyme | mtaB; threonylcarbamoyladenosine tRNA methylthiotransferase MtaB [EC:2.8.4.5] |
| 57 | PGA1_RS10805 | PGA1_c21820 | 5-formyltetrahydrofolate cyclo-ligase | 5-formyltetrahydrofolate cycloligase-like protein | MTHFS; 5-formyltetrahydrofolate cyclo-ligase [EC:6.3.3.2] |
| 58 | PGA1_RS16760 | PGA1_c33770 | ABC transporter permease subunit | putative ABC transporter permease protein | proW; glycine betaine/proline transport system permease protein |
| 59 | PGA1_RS18110 | PGA1_c36480 | bifunctional demethylmenaquinone methyltransferase/2-methoxy-6-polyprenyl-1,4-benzoquinol methylase UbiE | ubiquinone/menaquinone biosynthesis methyltransferase UbiE | ubiE; demethylmenaquinone methyltransferase / 2-methoxy-6-polyprenyl-1,4-benzoquinol methylase [EC:2.1.1.163 2.1.1.201] |
| 60 | PGA1_RS00260 | PGA1_c00520 | tRNA (guanosine(37)-N1)-methyltransferase TrmD | tRNA (guanine-N(1)-)-methyltransferase TrmD | trmD; tRNA (guanine37-N1)-methyltransferase [EC:2.1.1.228] |
| 61 | PGA1_RS07960 | PGA1_c16050 | methylenetetrahydrofolate reductase | hypothetical protein | metF, MTHFR; methylenetetrahydrofolate reductase (NADH) [EC:1.5.1.54] |
| 62 | PGA1_RS01215 | PGA1_c02510 | S-(hydroxymethyl)glutathione dehydrogenase/class III alcohol dehydrogenase | S-(hydroxymethyl)glutathione dehydrogenase FrmA | frmA, ADH5, adhC; S-(hydroxymethyl)glutathione dehydrogenase / alcohol dehydrogenase [EC:1.1.1.284 1.1.1.1] |
| 63 | PGA1_RS07955 | PGA1_c16040 | methyltetrahydrofolate cobalamin methyltransferase | pterin binding domain-containing protein | metH, MTR; 5-methyltetrahydrofolate--homocysteine methyltransferase [EC:2.1.1.13] |
| 64 | PGA1_RS10490 | PGA1_c21180 | thymidylate synthase | thymidylate synthase ThyA | thyA, TYMS; thymidylate synthase [EC:2.1.1.45] |
| 65 | PGA1_RS13640 | PGA1_c27430 | cobalamin-independent methionine synthase II family protein | 2-hydroxypropyl-CoM lyase-like protein | metE; 5-methyltetrahydropteroyltriglutamate--homocysteine methyltransferase [EC:2.1.1.14] |
| 66 | PGA1_RS01920 | PGA1_c03890 | autoinducer synthase | putative acyl-homoserine-lactone synthase RaiI | raiI; acyl homoserine lactone synthase [EC:2.3.1.184] |
| 67 | PGA1_RS05915 | PGA1_c11870 | serine hydroxymethyltransferase | serine hydroxymethyltransferase GlyA | glyA, SHMT; glycine hydroxymethyltransferase [EC:2.1.2.1] |
| 68 | PGA1_RS02760 | PGA1_c05560 | glycine cleavage system protein GcvH | glycine cleavage system protein GcvH | gcvH, GCSH; glycine cleavage system H protein |
| 69 | PGA1_RS15700 | PGA1_c31600 | S-formylglutathione hydrolase | S-formylglutathione hydrolase FghA | frmB, ESD, fghA; S-formylglutathione hydrolase [EC:3.1.2.12] |
| 70 | PGA1_RS16975 | PGA1_c34190 | class I SAM-dependent methyltransferase | methyltransferase-like protein | |
| 71 | PGA1_RS19450 | PGA1_78p00300 | glycine cleavage system protein GcvH | glycine cleavage system protein GcvH | gcvH, GCSH; glycine cleavage system H protein |
| 72 | PGA1_RS00460 | PGA1_c00940 | class I SAM-dependent methyltransferase | putative methyltransferase | |
| 73 | PGA1_RS19145 | PGA1_262p02100 | protein-glutamate O-methyltransferase | methyltransferase, cheR type | cheR; chemotaxis protein methyltransferase CheR [EC:2.1.1.80] |
| 74 | PGA1_RS04710 | PGA1_c09490 | 16S rRNA (cytosine(1402)-N(4))-methyltransferase RsmH | S-adenosyl-L-methionine-dependent methyltransferase MraW | mraW, rsmH; 16S rRNA (cytosine1402-N4)-methyltransferase [EC:2.1.1.199] |
| 75 | PGA1_RS06910 | PGA1_c13880 | BCCT family transporter | putative transporter, BCCT family | TC.BCT; betaine/carnitine transporter, BCCT family |
| 76 | PGA1_RS17590 | PGA1_c35420 | adenosylhomocysteinase | adenosylhomocysteinase AhcY | E3.3.1.1, ahcY; adenosylhomocysteinase [EC:3.3.1.1] |
| 77 | PGA1_RS16660 | PGA1_c33570 | site-specific DNA-methyltransferase | modification methylase BabI | ccrM; modification methylase [EC:2.1.1.72] |
| 78 | PGA1_RS14200 | PGA1_c28550 | class I SAM-dependent methyltransferase | methyltransferase | evaC; methylation protein EvaC |
| 79 | PGA1_RS05830 | PGA1_c11700 | methylenetetrahydrofolate reductase [NAD(P)H] | 5,10-methylenetetrahydrofolate reductase MetF | metF, MTHFR; methylenetetrahydrofolate reductase (NADH) [EC:1.5.1.54] |
| 80 | PGA1_RS19455 | PGA1_78p00310 | aminomethyl-transferring glycine dehydrogenase | glycine dehydrogenase | GLDC, gcvP; glycine dehydrogenase [EC:1.4.4.2] |
| 81 | PGA1_RS18555 | PGA1_262p00860 | tRNA (N6-threonylcarbamoyladenosine(37)-N6)-methyltransferase TrmO | hypothetical protein | TRMO, trmO; tRNA (adenine37-N6)-methyltransferase [EC:2.1.1.-] |
| 82 | PGA1_RS06660 | PGA1_c13370 | betaine--homocysteine S-methyltransferase | putative homocysteine S-methyltransferase | metH, MTR; 5-methyltetrahydrofolate--homocysteine methyltransferase [EC:2.1.1.13] |
| 83 | PGA1_RS02370 | PGA1_c04780 | methionine adenosyltransferase | S-adenosylmethionine synthetase MetK | metK, MAT; S-adenosylmethionine synthetase [EC:2.5.1.6] |

**Table S4.** Comparison of lag phase shortening responses in *P. inhibens* bacteria induced by various compounds. Differences in lag phase shortening upon treatment with 2 µM of various compounds were reported in Figs. 3A and 3B, and the statistical significance of the difference between responses was calculated. The comparison of lag phase shortening induced by various methylated compounds within the same group ("Methylated compound groups" in the table) was assessed using Tukey's multiple comparisons test. In cases where there were only two compounds in a group, an unpaired t-test was conducted. Adjusted *p*-value thresholds: **** ≤ 0.0001, *** ≤ 0.001, ** ≤ 0.01, * ≤ 0.05, ns [not significant] > 0.05.

| Methylated compound groups | Compounds comparison | Significance | Adjusted *p*-value |
|---|---|---|---|
| DMSP (and analogues) | Acetate vs. Propionate | ns | 0.5294 |
| | Acetate vs. MPA | ns | 0.9998 |
| | Acetate vs. 3-MMPA | ** | 0.0019 |
| | Acetate vs. DMSP | **** | <0.0001 |
| | Propionate vs. MPA | ns | 0.4357 |
| | Propionate vs. 3-MMPA | *** | 0.0001 |
| | Propionate vs. DMSP | **** | <0.0001 |
| | MPA vs. 3-MMPA | ** | 0.0027 |
| | MPA vs. DMSP | **** | <0.0001 |
| | 3-MMPA vs. DMSP | ** | 0.0033 |
| *N*-methylated quaternary ammonium compounds (and analogues) | Betaine vs. Choline | ns | 0.2249 |
| | Betaine vs. DMG | ns | 0.6042 |
| | Betaine vs. Sarcosine | *** | 0.0003 |
| | Betaine vs. Alanine | **** | <0.0001 |
| | Betaine vs. Carnitine | ns | 0.9925 |
| | Choline vs. DMG | * | 0.0101 |
| | Choline vs. Sarcosine | **** | <0.0001 |
| | Choline vs. Alanine | **** | <0.0001 |
| | Choline vs. Carnitine | ns | 0.4946 |
| | DMG vs. Sarcosine | ** | 0.0079 |
| | DMG vs. Alanine | *** | 0.0002 |
| | DMG vs. Carnitine | ns | 0.2993 |
| | Sarcosine vs. Alanine | ns | 0.5489 |
| | Sarcosine vs. Carnitine | **** | <0.0001 |
| | Alanine vs. Carnitine | **** | <0.0001 |
| Stachydrine (and analogues) | Proline vs. Stachydrine | **** | <0.0001 |
| *N*-methylated aromatic compounds (and analogues) | Trigonelline vs. Homarine | ns | 0.7399 |
| | Trigonelline vs. Nicotinic acid | * | 0.0316 |
| | Homarine vs. Nicotinic acid | * | 0.01 |
| *S*-methylated sulfonium compounds (and analogues) | Gonyol vs. DMSA | * | 0.0299 |
| | Gonyol vs. Cysteine | *** | 0.0001 |
| | DMSA vs. Cysteine | ** | 0.0052 |
| One-carbon compounds | Methanol vs. Methanthiol | ns | 0.7912 |
| C1 group donating amino acids | Glycine vs. Serine | ns | 0.6225 |
| | Glycine vs. Methionine | **** | <0.0001 |
| | Serine vs. Methionine | **** | <0.0001 |

**Table S5:** Estimated methyl group requirements per *P. inhibens* bacterial cell. Values based on *E. coli* (Neidhardt *et al.,*[5])

| Building blocks that are C1 sinks | ATP (RNA) | GTP (RNA) | dATP (DNA) | dGTP (DNA) | dTTP (DNA) | Histidine[1] (proteins) | Methionine (proteins) |
|---|---|---|---|---|---|---|---|
| Class | Purine | Purine | Purine | Purine | Thymine | Amino acid | Amino acid |
| **Experimentally determined amount of building block in *E. coli* B/r**[3] | | | | | | | |
| [µmol building block/ g dried cells] | 165.0 | 203.0 | 24.7 | 25.4 | 24.7 | 90.0 | 146.0 |
| [amol building block/ cell] | 46.2 | 56.8 | 6.9 | 7.1 | 6.9 | 25.2 | 40.9 |
| **Incorporation of C1 groups during building block synthesis in *P. inhibens*** | | | | | | | |
| Amount of C1 groups incorporated per building block | 2 | 2 | 2 | 2 | 1 | 1 | 1 |
| Key enzyme(s) | 1: Phosphoribosyl-glycinamide formyltransferase 2: Phosphoribosyl-aminoimidazolecarboxamide formyltransferase | 1: Phosphoribosyl-glycinamide formyltransferase 2: Phosphoribosyl-aminoimidazolecarboxamide formyltransferase | 1: Phosphoribosyl-glycinamide formyltransferase 2: Phosphoribosyl-aminoimidazolecarboxamide formyltransferase | 1: Phosphoribosyl-glycinamide formyltransferase 2: Phosphoribosyl-aminoimidazolecarboxamide formyltransferase | Thymidylate synthase | Phosphoribosyl-aminoimidazolecarboxamide formyltransferase | Methionine synthase |
| C1 group donors | 1: CHO-THF 2: CHO-THF | 1: CHO-THF 2: CHO-THF | 1: CHO-THF 2: CHO-THF | 1: CHO-THF 2: CHO-THF | $CH_2$-THF | CHO-THF | $CH_3$-cobalamin, DMSP, betaine[2] |
| Accession (gene name, RefSeq, GennBank, protein) | 1: *purN*, PGA1_RS06610, PGA1_c13270, WP_014874448.1 2: *purH*, PGA1_RS16320, PGA1_c32860, WP_014881359.1 | 1: *purN*, PGA1_RS06610, PGA1_c13270, WP_014874448.1 2: *purH*, PGA1_RS16320, PGA1_c32860, WP_014881359.1 | 1: *purN*, PGA1_RS06610, PGA1_c13270, WP_014874448.1 2: *purH*, PGA1_RS16320, PGA1_c32860, WP_014881359.1 | 1: *purN*, PGA1_RS06610, PGA1_c13270, WP_014874448.1 2: *purH*, PGA1_RS16320, PGA1_c32860, WP_014881359.1 | *thyA*, PGA1_RS10490, PGA1_c21180, WP_014880433.1 | *purH*, PGA1_RS16320, PGA1_c32860, WP_014881359.1 | *bmt*, PGA1_RS06660, PGA1_c13370, WP_014879827.1 |
| Genes | 1: 33 2: 53 | 1: 33 2: 53 | 1: 33 2: 53 | 1: 33 2: 53 | 64 | 53 | 82 |
| Amount of C1 groups produced per building block[3] | 1 | 1 | 1 | 1 | 0 | 0 | 0 |
| Net C1 group requirements per cell [amol C1 group/cell] | 46.2 | 56.8 | 6.9 | 7.1 | 6.9 | 25.2 | 40.9 |

[1] Consumption of C1 groups during histidine synthesis occurs indirectly. Histidine is synthesized by condensing the adenine backbone of ATP with the ribose sugar 5-phospho-α-D-ribose 1-diphosphate (PRPP). The side product of histidine synthesis is 5-amino-1-(5-phospho-D-ribosyl)imidazole-4-carboxamide (AICAR). AICAR is channeled into the lower branch of the purine synthesis pathway to regenerate the initially consumed ATP. Thus, ATP regeneration consumes one formyl group per synthesized histidine.
histidine.

[2] Methionine synthesis in *P. inhibens* was reported to depend on the methionine synthase enzyme Bmt and a cobalamin binding protein. It was thus concluded that $CH_3$-cobalamine is the methyl group donor for Bmt[6]. However, the utilization of other methyl group donors by Bmt cannot be excluded[7].

[3] Purine synthesis consumes two C1 groups and releases one C1 group per produced building block[5]. The release of one C1 group arises from the incorporation of glycine during purine synthesis (Fig. 1C; gene 26: phosphoribosylamine—glycine ligase). Glycine is produced by converting glucose (via 3-phosphoglycerate) to serine and then to glycine, which releases one C1 group (Fig. 1C; gene 67: serine hydroxymethyltransferase).

**Table S6:** Sequencing yields of lag phase RNA-sequencing run.

| | Cycles | Yield (Gbp) | % > Q30 | Reads (M) | Reads passing filter (M) |
|---|---|---|---|---|---|
| Total | 168 | 20.77 | 90.78 | 338.1 | 249.3 |
| Read 1 | 89 | 11.01 | 90.11 | 169.1 | 124.7 |
| Read 2 | 79 | 9.76 | 91.55 | 169.1 | 124.7 |

**Table S7:** Bacterial read counts per sample of lag phase RNA-sequencing run.

| Sample ID \| replicate | Paired-end reads (after demultiplexing) | Paired-end reads (after cutadapt) | *Phaeobacter inhibens* feature counts | |
|---|---|---|---|---|
| | | | total | non-rRNA |
| Control (15 min lag) \| rep 1 | 5,173,156 | 5,172,928 | 4,912,488 | 4,786,267 |
| Control (15 min lag) \| rep 2 | 6,256,704 | 6,256,419 | 5,938,559 | 5,783,725 |
| Control (15 min lag) \| rep 3 | 5,656,899 | 5,656,722 | 5,360,067 | 5,214,107 |
| DMSP (15 min lag) \| rep 1 | 7,299,641 | 7,299,366 | 6,798,582 | 6,591,949 |
| DMSP (15 min lag) \| rep 2 | 4,585,425 | 4,585,264 | 4,232,884 | 4,090,929 |
| DMSP (15 min lag) \| rep 3 | 7,399,140 | 7,398,864 | 6,915,209 | 6,708,565 |
| Control (40 min lag) \| rep 1 | 5,816,609 | 5,816,342 | 5,537,252 | 5,402,090 |
| Control (40 min lag) \| rep 2 | 6,157,332 | 6,157,141 | 5,857,923 | 5,714,672 |
| Control (40 min lag) \| rep 3 | 5,616,052 | 5,615,798 | 5,376,958 | 5,257,722 |
| DMSP (40 min lag) \| rep 1 | 5,862,598 | 5,862,395 | 5,554,342 | 5,414,854 |
| DMSP (40 min lag) \| rep 2 | 6,068,667 | 6,068,472 | 5,711,286 | 5,562,658 |
| DMSP (40 min lag) \| rep 3 | 5,405,311 | 5,405,127 | 5,067,317 | 4,933,000 |

**Table S8:** Bacterial genomes used for analysis of methyl group-related metabolism in algal-associated and plant-associated bacteria (see fig. S18).

| **Bacteria** | **NCBI RefSeq / repository** |
|---|---|
| *Labrenzia* sp. THAF82 | GCF_009363435.1 |
| *Vibrio harveyi* SB1 | GCF_030060435.1 |
| *Marinomonas algarum* E8 | GCF_020532605.1 |
| *Pseudoalteromonas phenolica* KCTC 12086 | GCF_001444405.1 |
| *Bacillus subtilis* 168 | GCF_000009045.1 |
| *Pseudomonas koreensis* LMG 21318 | GCF_900101415.1 |
| *Pseudomonas aeruginosa* PAO1 | GCF_000006765.1 |
| *Ewingella americana* CCUG 14506T | GCF_008693045.1 |
| *Lelliottia amnigena* FDAARGOS_1445 | GCF_019355955.1 |
| *Klebsiella aerogenes* Ka37751 (previously *Enterobacter aerogenes*) | GCF_007632255.1 |
| *Sulfitobacter pontiacus* iR4 | https://doi.org/10.5281/zenodo.7520022 |

**Data S1:** *P. inhibens* feature table (genes) with results from co-cultivation RNA-sequencing run. The dataset includes bacterial gene accession numbers, functional annotations, transcript abundances (TPM normalized read counts) and results of DESeq2 differential gene expression analysis. (Fig. 1, figs. S1A, S3-S4, tables S1-S2).
Download: https://weizmann.box.com/shared/static/qg8pqrhjff5paoapsnepc00cbuv5swpq.xlsx
*(temporary link for review).*

**Data S2:** *P. inhibens* feature table (genes) with results from lag phase RNA-sequencing run. The dataset includes bacterial gene accession numbers, functional annotations, transcript abundances (TPM normalized read counts) and results of DESeq2 differential gene expression analysis. (Fig. 4A, figs. S10-S13, tables S6-S7).
Download:
https://weizmann.box.com/shared/static/5pgvd3715nuwkeygt0fkvshuee8jplgj.xlsx
*(temporary link for review).*

**Data S3:** *Emiliania huxleyi* CCMP3266 sGenome gene annotation file version 2 (GFF3 format). Download: https://weizmann.box.com/shared/static/rr1hbu2zdnmfmnmia461ogspxdiib43e.gff
*(temporary link for review).*

**Data S4:** Feature quantification obtained using Compound Discoverer. The table presents the output analysis using Compound Discoverer (v3.3) with a putative identification of metabolites. Each identified metabolite (each row) contains a sub-table under the + tab (on the left side) that specifies the feature quantification in each analyzed sample. Sample ID appears in the “Study File ID” column in each sub-table. Values in the columns “Exchange Rate [%]: 0” and “Exchange Rate [%]: 1” represent relative abundances of the molecules in their unlabeled form and with single $^{13}C$-label (M+1 isotopologue), respectively. Compounds marked with “1” in the “Tags” column were further validated using standards. The average M+1 isotope abundance [%] for *S*-Adenosylmethionine (SAM) and 5'-*S*-Methyl-5'-thioadenosine (MTA)—as reported in Fig. 4D—were calculated by averaging the values in the “Exchange Rate [%]: 1” column from samples supplemented with $^{13}C$-labeled and unlabeled DMSP, respectively. Detailed information regarding the isotope abundances of SAM and MTA can be found in the tab “SAM and MTA isotope abundance” in the table. In the tab “Features Positive Mode” samples F10, F12, F14, and F16 were supplemented with $^{13}C$-labeled DMSP and samples F2, F4, F6, F8 were supplemented with unlabeled DMSP. In the tab “Features Negative Mode” samples F2, F3, F4, and F5 were supplemented with $^{13}C$-labeled DMSP and samples F10, F11, F12, F13 were supplemented with unlabeled DMSP.
Download: https://weizmann.box.com/s/1akxr55fqpg5d7w5omnprledqnnxfpls
*(temporary link for review).*
